# FGF signaling induces disease-associated oligodendrocyte states and drives myelin pathology

**DOI:** 10.64898/2026.11.03.764025

**Authors:** Maya K. Weigel, Madeline H. Cooper, Mattia Zaghi, Jayakrishnan Gopakumar, Caldwell M. Bridges, Peggy P. Ho, Michael Schoof, Ugne Klibaite, Stacey Pedraza, Eneritz Agirre, George Tilton-Low, Nadine Bestard-Cuche, Petra Kukanja, Veronika V. Rodriguez, Graham Jones, John J. Perrino, Danielle M. Jorgens, Mark R. Terasaki, Anna Williams, Lawrence Steinman, Gonçalo Castelo-Branco, J. Bradley Zuchero

## Abstract

Myelin integrity is essential for normal central nervous system (CNS) function and is disrupted in neuroinflammatory and neurodegenerative disorders, including multiple sclerosis and Alzheimer’s disease. In these settings, myelin-forming oligodendrocytes adopt a conserved disease-associated transcriptional state, termed disease-associated oligodendrocytes (DAOs). How DAOs arise, the signals that drive their induction, and whether they actively contribute to pathology remain poorly understood. Here we show that FGF signaling instructs mature oligodendrocytes to enter disease-associated states that are sufficient to trigger myelin pathology. By screening disease-linked secreted factors, we identify FGF1 and FGF2 as potent inducers of a DAO-like state in purified oligodendrocyte cultures, characterized by transcriptional reprogramming and myelin-membrane shedding. In vivo, activating oligodendrocyte FGF signaling is sufficient to elicit DAO programs and myelin disruption, whereas inhibiting FGF signaling suppresses DAO induction in a mouse model of multiple sclerosis. Finally, analysis of post-mortem multiple sclerosis tissue reveals a concordant DAO-like state associated with increased engagement of the FGF/FGFR axis. Collectively, these studies define a regulated mechanism of DAO formation linked to myelin disruption and position mature oligodendrocytes as active contributors to myelin pathology in CNS disease.

## Introduction

Myelination is essential for rapid and precise signal conduction in the central nervous system (CNS). Myelin loss and dysfunction of myelinating oligodendrocytes are common across diverse neurological diseases including multiple sclerosis (MS) and Alzheimer’s disease, where myelin pathology contributes to disability and cognitive decline^1,2,3^. Although major upstream drivers of disease are increasingly well defined (for example, evidence implicating Epstein–Barr virus in MS risk and pathophysiology^4,5,6,7^), how these initiating events are translated into oligodendrocyte dysfunction and myelin damage remains unclear.

Recent single-cell and single-nucleus transcriptomic studies across human disease tissue and mouse models have converged on a striking observation: oligodendrocyte lineage cells adopt conserved disease-associated transcriptional programs marked by reduced expression of homeostatic myelin genes and induction of stress and immune-related pathways^8,9,10,11,12,13,14,15,16,17,18,19,20,21^. These “disease-associated oligodendrocytes” (DAOs), often marked in mice by Serpina3n expression^13,16,18,19,20^, have been proposed to represent a common cellular nexus across CNS pathologies and a potential therapeutic target for preventing or reversing demyelination^22^. However, DAOs have been defined primarily through descriptive transcriptomic atlases, leaving key mechanistic questions unresolved: Do DAOs derive from mature oligodendrocytes, from oligodendrocyte precursor cells (OPCs) undergoing perturbed differentiation, or both? What extracellular signals instruct DAO formation in vivo? And are DAO programs maladaptive—directly contributing to myelin damage—or instead protective responses to inflammation and injury?

Fibroblast growth factor (FGF) signaling has long been linked to demyelinating disease and oligodendrocyte biology. In particular, FGF2 can suppress myelin gene expression in cultured oligodendrocytes^23,24,25,26,27,28^ and perturb myelin structure in vivo^29,30,31^, and genetic or pharmacologic disruption of FGFR signaling is protective in the experimental autoimmune encephalomyelitis (EAE) mouse model of human MS^32,33,34,35^. However, the mechanistic interpretation of these findings—and whether they relate to DAO biology—remains uncertain. We hypothesized that FGFs may act not merely as nonspecific inhibitors of myelin gene expression, but as upstream instructive cues that drive mature oligodendrocytes into conserved DAO programs with functional consequences for myelin integrity.

In this study, we asked whether DAOs can be induced by defined extracellular cues and whether this transition has functional consequences for myelin. We show that FGF signaling acts directly on mature oligodendrocytes to drive entry into conserved disease-associated states and that this transition is sufficient to destabilize myelin in vivo. This framework helps explain longstanding links between FGF signaling and demyelinating disease and identifies a tractable pathway for modulating oligodendrocyte disease-state transitions.

## Results

### Disease-associated oligodendrocytes (DAOs) primarily derive from mature oligodendrocytes in EAE

The abundant presence of DAOs is well established in numerous demyelinating diseases and aging, however, the cell type of origin remains unresolved^22^. In principle, DAOs could originate from (1) mature, myelinating oligodendrocytes, (2) oligodendrocyte precursor cells (OPCs) undergoing perturbed differentiation (e.g., attempting to remyelinate following demyelination), or (3) another CNS cell type. To define which cell type DAOs originate from, we used genetic lineage tracing coupled with EAE, which models key aspects of human MS including neuroinflammation, peripheral immune cell infiltration, demyelination, and—importantly—robust induction of DAOs^13,19,36^. We tracked the fate of mature oligodendrocytes using *Plp1-CreERT;Tau-mGFP-nLacZ* reporter mice, in which tamoxifen induces recombination to express membrane-bound GFP (mGFP) and nuclear lacZ (nLacZ) alleles only in mature oligodendrocytes^37^. We confirmed that tamoxifen-induced reporter expression was 100% specific for mature oligodendrocytes, with no expression ever observed in OPCs or other cell types (**Extended Data Fig. 1a,b**). Following tamoxifen injection, >75% of all Olig2/CC1 double-positive mature oligodendrocytes efficiently recombined to express both beta-galactosidase (β-gal, product of LacZ gene) and mGFP (**Extended Data Fig. 1c,d**), allowing reliable analysis of mature oligodendrocyte fate during disease modeling.

We assessed DAO induction in the spinal cords of MOG^35–55^ EAE-induced reporter mice by quantifying oligodendrocyte lineage cell expression of Serpina3n, the primary marker of DAOs across CNS diseases^13,18,19,20,36,38^ (**Extended Data Fig. 2a–c**). At peak EAE clinical score (the time of peak DAO induction in MOG^35–55^ EAE in prior reports^13,19,36^), Serpina3n+ cells were spread throughout the spinal cord white matter, not just around lesion areas (**Extended Data Fig. 2c and 3c**), as previously reported^36^. Mature oligodendrocytes expressed high levels of Serpina3n throughout the spinal cord white matter in diseased mice, in stark contrast to mature oligodendrocytes in control mice, in which Serpina3n expression was undetectable (**Extended Data Fig. 2d**). Strikingly, we found that the majority (~87%) of Serpina3n+ oligodendrocyte lineage cells expressed β-gal, indicating that most DAOs derive from mature oligodendrocytes in this model (**Extended Data Fig. 2e**). Mature oligodendrocytes also upregulated other common DAO markers including MHC-I gene B2m and the nuclear alarmin IL33 in EAE (**Extended Data Fig. 14g,h**). The density of mature oligodendrocytes in lesion-adjacent EAE spinal cords was unchanged (**Extended Data Fig. 3b**), however, mature oligodendrocytes in EAE had hypertrophied nuclei and dysmorphic somas (**Extended Data Fig. 2f,g**). Oligodendrocytes around MS lesions have been shown to undergo similar morphologic changes^39,40^. Together, our findings establish that DAOs primarily arise from mature oligodendrocytes in EAE, and become morphologically distinct from homeostatic oligodendrocytes (**Extended Data Fig. 2h**).

### FGF1 and FGF2 are signals that switch mature oligodendrocytes into a DAO state

What signals induce mature oligodendrocytes to become DAOs? We screened candidate disease-linked, secreted proteins for DAO-inducing potential using an established primary culture system of highly pure rodent immunopanned oligodendrocytes^41^ (**Fig. 1a**). Candidate proteins included inflammatory cytokines (e.g., IFN-γ, TNF-a, IL-17), mediators of neurodegeneration (e.g., ApoE4, Amyloid-β), and growth factors with potential roles in disease (e.g., EGF, BDNF, FGFs). After differentiating oligodendrocytes from OPCs for 6 days, the resulting oligodendrocyte cultures are ~99% pure (MOG+, MBP+), with gene expression that closely resembles mature oligodendrocytes in vivo, making the system a powerful tool to model myelin and oligodendrocyte cell biology^42,43^. As a proxy for DAO induction, we quantified expression changes in established DAO markers Serpina3n and Vimentin (Vim)^18^ as well as loss of myelin basic protein (MBP), as downregulation of homeostatic mature myelin genes is a common feature of DAO activation^18^. Interestingly, we identified the fibroblast growth factor family as signaling molecules sufficient to drive induction of DAO markers (**Fig. 1b–h**) and downregulation of mature oligodendrocyte genes (**Fig. 1b–d,i**), including MBP as previously reported for FGF2^23,24,25,26,27^. Of note, Interferon gamma (IFN-γ), which has been shown to drive oligodendrocyte lineage cells into a specific interferon-responsive DAO subclass^13,38^, did not lead to increased expression of DAO markers. We focused on FGF1 and FGF2 (hereafter, FGF1/2) since they both activate all FGF receptors^44^, strongly induced a DAO-like state in culture (**Extended Data Fig. 4a,b**), and have been linked to glial reactivity^45^ and demyelinating disease^46,47,48,49,50^.

**Figure 1.**
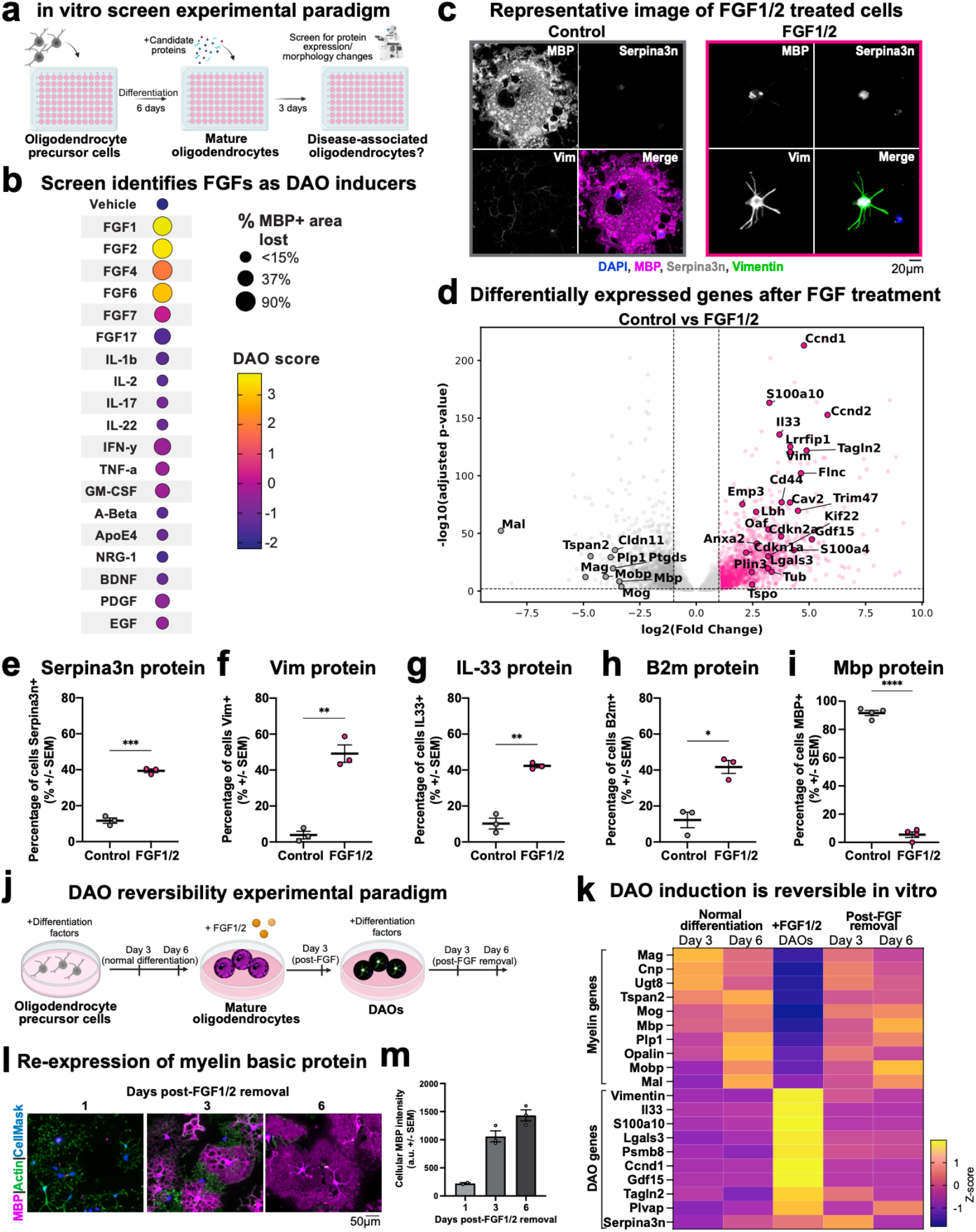
Identification of FGF1 and FGF2 as potent inducers of a DAO state in culture. a, Cartoon schematic of plating scheme for screen. Mature (6-day differentiated) oligodendrocytes treated with candidate molecules for 3 days. b, Dot-plot showing results of screen for DAO-inducing molecules. Dot size represents percent of MBP+ area lost and color represents composite z-score of Vimentin and Serpina3n expression. c, Representative image of immunostaining for MBP, Vimentin and Serpina3n in control and FGF treated oligodendrocytes. d, Volcano plot visualization of bulk RNA-seq of control (20ng/ml 0.2% BSA + heparin, 72 hours) and FGF1/2 treated (10ng/ml FGF1 + 10ng/ml FGF2 + heparin, 72 hours) oligodendrocytes shows that FGF1/2 induces coordinated DAO marker programs (not merely loss of myelin genes); n = 3 biological replicates. (e-i) Quantification of immunostaining represented by percentage of positive cells. Percentage of cells expressing each marker ± SEM: e, Serpina3n – Control: 11.69 ± 1.49%, FGF1/2: 39.40 ± 0.98%; f, Vimentin – Control: 3.85 ± 2.1%, FGF1/2: 49.16 ± 4.84%; g, IL33 – Control: 10.23 ± 2.99%, FGF1/2: 42.28 ± 1.0%; h, B2m – Control: 12.26 ± 4.31%, FGF1/2: 41.67 ± 3.58%, n = 3 biological replicates; i, MBP – Control: 91.59 ± 1.81%, FGF1/2: 5.49 ± 1.97%; n = 4 biological replicates. Also see Extended Data Fig. 4d. j, Experimental paradigm for DAO reversibility experiment shown in k. Timepoint conditions for bulk RNA-seq were day 3 differentiated, day 6 differentiated, FGF1/2 treated, day 3 post-FGF1/2 removal, day 6 post-FGF1/2 removal oligodendrocytes. k, Heatmap showing expression of mature oligodendrocyte/myelin genes (top) and disease-associated oligodendrocyte genes (bottom) in cultured oligodendrocytes for timepoints shown in j; n = 3 biological replicates. l, Representative images showing oligodendrocytes previously treated with FGF1/2 1, 3, and 6 days after replating in differentiation media in the absence FGF. m, Quantification of cellular MBP intensity immunostaining shown in l, MBP intensity per cell (a.u.) ± SEM; 1 day post-FGF removal: 222.2 ± 15.15; 3 days post-FGF removal: 1,059 ± 97.67; 6 days post-FGF removal 1,424 ± 99.22, n = 3 biological replicates. Statistical measurements (p value) were determined by paired, one-tailed t-test, p values: *p < 0.05; **p < 0.01; ***p < 0.001; ****p < 0.0001.x

To test in an unbiased manner how FGF1/2 affects mature oligodendrocytes, we performed bulk RNA-seq of fully differentiated oligodendrocytes treated with either FGF1/2 or a vehicle control for 3 days. FGF1/2 induced a dramatic upregulation of many previously reported DAO genes^13,18,20^ (e.g., *Serpina3n, Il33, Vim, Gdf15, Tagln2, S100a10*) while markers of mature, fully differentiated oligodendrocytes (e.g., *Mbp, Mog, Mag, Plp1)* were downregulated (**Fig. 1d,i**). In contrast, FGF1/2 did not induce expression of established marker genes of OPCs, astrocytes, microglia, or other CNS cell types (**Extended Data Fig. 4c**). We validated these gene expression changes using immunostaining (**Fig. 1c,e–i**) and unbiased proteomics (**Extended Data Fig. 4e– g**). Quantification confirmed increased protein expression of many key DAO markers including Serpina3n, IL33, B2m, and Vimentin (**Fig. 1e–h and Extended Data Fig. 4d**) and decreased expression of mature oligodendrocyte proteins including MBP (**Fig. 1i**). Interestingly, this transcriptional reprogramming of cultured oligodendrocytes in response to FGF1/2 was reversible upon removal of FGF from the culture media (**Fig. 1j–m**). Although FGF2 has been reported to suppress myelin gene expression in culture^23,24,25,26,27^, it was unclear whether this reflects a defined disease-associated oligodendrocyte state. Our data show that FGF1/2 induces a coordinated DAO transcriptional and proteomic program rather than an isolated loss of myelin markers. Together, these results indicate that FGF1/2 are sufficient to induce mature oligodendrocytes to enter a DAO-like state, and establish a robust model for studying DAOs in culture.

### FGF signaling is sufficient in vivo to generate conserved DAO classes

Given our results that activating FGF signaling induces oligodendrocytes to express DAO markers in culture, we next asked whether FGF1/2 are sufficient to generate bona fide DAO classes in vivo. Reactive astrocytes upregulate and are the main source of FGF1 and FGF2 in disease and injury^45,46,51,52,53,^. Thus, to mimic such disease states, we designed astrocyte-specific^54^ adeno-associated viruses (AAVs) to drive secretion of FGF1 and FGF2 from astrocytes in the mouse CNS (**Fig. 2a**). Astrocyte-specific tdTomato (red fluorescent protein) served as a control for potential nonspecific effects of viral infection as well as to mark the injection site. We first validated that FGF AAVs were sufficient to induce secretion of functional FGF from primary cultured astrocytes (**Extended Data Fig. 5a,b**). In vivo, we validated the astrocyte-specificity of our AAVs (**Extended Data Fig. 5c**) and confirmed expression of FGF1/2 using RNAscope in situ hybridization with custom probes to detect ectopically expressed (mouse codon-optimized) *Fgf1* and *Fgf2* transcripts (**Extended Data Fig. 5d**). We typically observed astrocyte tdTomato expression restricted to a ~1-2 mm^3^ volume under these conditions.

**Figure 2.**
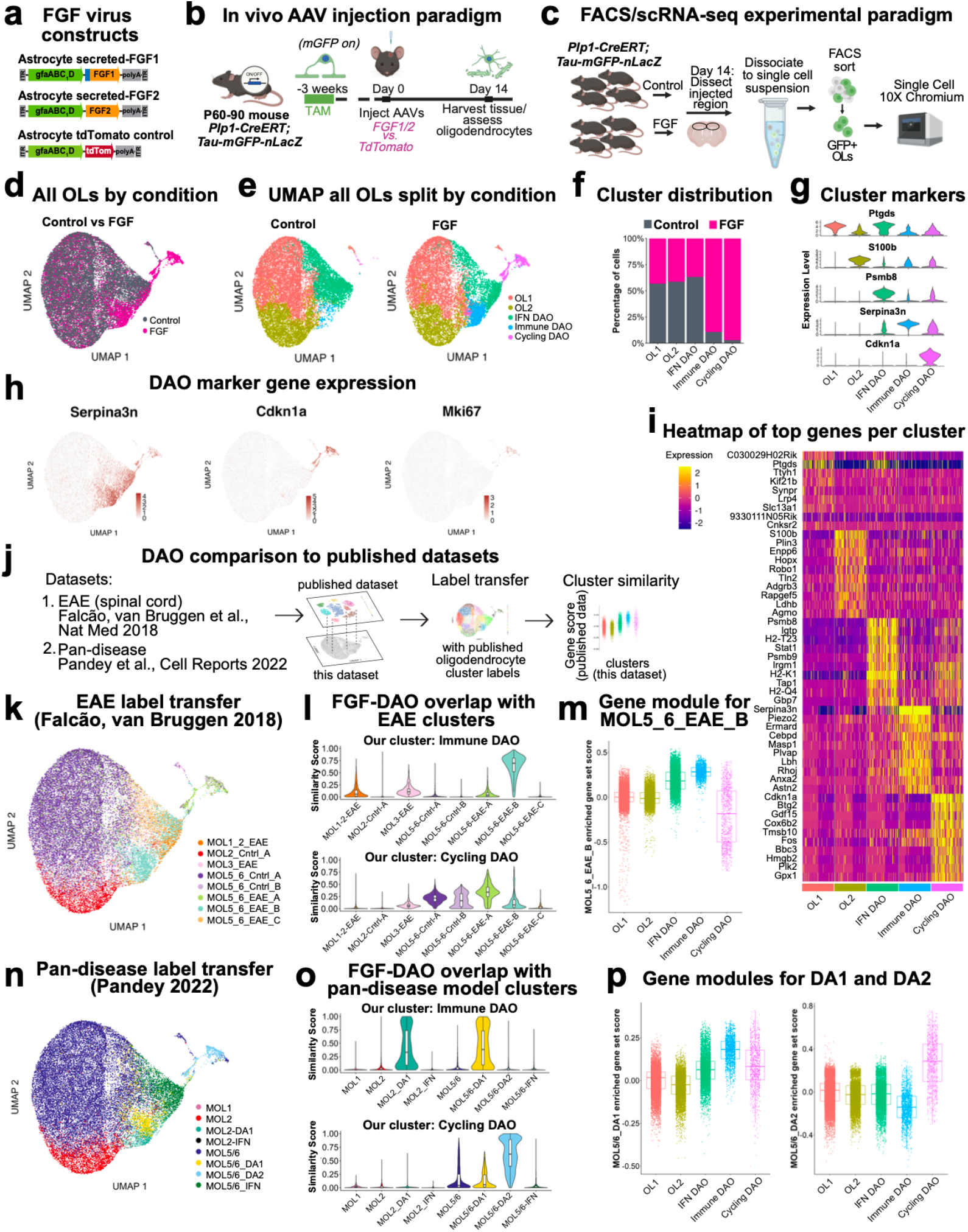
FGF induces mature oligodendrocytes into DAO states in vivo. a, Schematics of constructs packaged into AAVs to force secretion of FGF1 or FGF2 from astrocytes or express tdTomato in astrocytes. b, Schematic showing experimental paradigm for injection of AAVs into reporter mice. Mice were injected with either a cocktail of AAVs encoding FGF1, FGF2, and tdTomato (FGF1/2-AAV), or AAV concentration-matched tdTomato (control-AAV) 3 weeks after tamoxifen and assessed via scRNA-seq (Fig. 2) or immunostaining (Fig. 3) after 2 weeks. c, Schematic showing tissue and cell processing for FACS and scRNA-seq. Tissue punches from 4 mice were pooled for each sample. Total number of cells is 15,383 for controls and 13,045 for FGF. d, Uniform manifold approximation and projection (UMAP) plot showing all mature oligodendrocytes sequenced from integrated control-AAV and FGF1/2-AAV injected samples with colors representing condition (Grey = Control; Pink = FGF). e, UMAP visualizations of all mature oligodendrocytes in Control (left) and FGF (right) samples with colors representing different mature oligodendrocyte cell clusters. f, Cluster distribution showing proportion of each cluster per condition (Grey = Control; Pink = FGF). g, Violin plots depicting expression of the top differentially expressed gene in each cluster across all other cell clusters. h, UMAP visualizations with expression of key DAO genes (Serpina3n, Cdkn1a, Mki67) shown in red. i, Unbiased heatmap of top genes expressed in each mature oligodendrocyte cluster (Yellow = upregulated; Pink = downregulated). j, Workflow for single-cell dataset comparison of scRNA-seq data from FGF1/2-overexpression (this dataset) to EAE dataset13 and mouse pan-disease dataset (lysolecithin, cuprizone, and Alzheimer’s disease models18). k, UMAP depicting Falcão, van Bruggen et al., Nature Med 201813 EAE cell clusters mapped to this study’s dataset using label transfer. l, Similarity scores between FGF-induced Immune DAOs (top) and Cycling DAOs (bottom) compared to all cell clusters from Falcão, van Bruggen 2018 EAE dataset. X-axes designate oligodendrocyte clusters/subtypes described in Falcão, van Bruggen et al.13. m, Gene set scores derived from comparing genes from each cluster in this dataset to genes from Falcão, van Bruggen 2018 “MOL5/6_EAE_B”. A higher score indicates a given cluster is more highly enriched for genes in the MOL5/6_EAE_B population. n, UMAP depicting Pandey et al., Cell Reports 202218 pan-disease cell clusters mapped to this study’s dataset using label transfer. o, Similarity scores between FGF-induced Immune DAOs (top) and Cycling DAOs (bottom) compared to all cell clusters from Pandey 2022 pan-disease dataset. X-axes designate oligodendrocyte clusters/subtypes described in Pandey et al.18. p, Gene set scores derived from comparing genes from each cluster in this dataset to genes from Pandey 2022 “MOL5/6_DA1” and “MOL5/6_DA2”. A higher score indicates a given cluster is more highly enriched for genes in the DA1 or DA2 population.

To determine whether FGF1/2 are sufficient to induce DAO formation in vivo, we stereotactically injected either a cocktail of AAVs encoding FGF1, FGF2, and tdTomato (FGF1/2-AAV), or concentration-matched AAV encoding tdTomato (control-AAV) alone as a control, into the cingulum of adult *Plp1-CreERT;Tau-mGFP-nLacZ* reporter mice (**Fig. 2b**). Two weeks following injection, we isolated mGFP-positive oligodendrocytes from the injection area by FACS and performed single-cell RNA sequencing (scRNA-seq) on isolated cells using the 10x Genomics platform (**Fig. 2c**). After filtering low-quality reads (**Extended Data Fig. 6a**) and integrating control and FGF samples, we confirmed that the majority of cells (>90%) expressed canonical markers of mature oligodendrocytes (*Mag, Plp1, Mog*)^55^, with few contaminating cells (**Extended Data Fig. 6b–d**). Re-clustering only the mature oligodendrocytes revealed five distinct populations (**Fig. 2e and Extended Data Fig. 6e**), which we then compared to oligodendrocytes from disease models with robust DAO formation^13,18^ (**Fig. 2j**). Cells in all five clusters expressed mature myelin genes and did not express canonical OPC genes (**Extended Data Fig. 7g,h**). Three of the five mature oligodendrocyte clusters were found in both control and FGF samples (**Fig. 2d,f**), including two major populations of homeostatic mature oligodendrocytes, which we call “OL1” (marked by *Ptgds*) and “OL2” (*Plin3, S100b*) (**Fig. 2g and Extended Data Fig. 6f**), similar to previously described “MOL5/6” and “MOL2”^13,18,55^ (**Extended Data Fig. 8a,b**). A third shared population of mature oligodendrocytes matched previously reported interferon-responsive oligodendrocytes (**Extended Data Fig. 8a,b** “IFN DAO” cluster), with expression of genes like *Igtp, Stat1*, and *Psmb*9 (**Fig. 2i and Extended Data Fig. 6f**). It is likely that interferon-responsive oligodendrocytes are induced from an injection site response or viral response given that they exist in both control-AAV and FGF1/2-AAV injected samples (**Fig. 2e,f**). We therefore treat the shared IFN DAO cluster as a procedure-associated interferon response and focus subsequent analyses on the FGF-specific DAO populations.

Two DAO clusters were only found in the presence of FGF1/2 and were not observed in control-AAV samples despite the same surgical and viral context, indicating that their induction is ligand-dependent rather than a nonspecific injection response (**Fig. 2f**). The first cluster (which we designate “Immune DAOs”) highly expressed *Serpina3n* and other genes characteristic of DAOs found across disease models (**Fig. 2g–i and Extended Data Fig. 7c**). These Immune DAOs are highly similar to previously described “MOL5_6_EAE_B,” the Serpina3n-expressing DAO population in EAE spinal cord^13^ (**Fig. 2k–m and Extended Data Fig. 8c**), as well as previously described “DA1,” the Serpina3n-expressing DAO population in other mouse models of demyelination including lysolecithin, cuprizone, and Alzheimer’s disease models^18^ (**Fig. 2n–p and Extended Data Fig. 8d**). Gene Ontology (GO) term^56,57^ and KEGG enrichment analyses^58,59^ suggest the cells in this cluster are highly metabolically active (**Extended Data Fig. 6g**) and enriched in biological processes present in many neurologic diseases (**Extended Data Fig. 6h**). The second cluster (which we designate “Cycling DAOs”) was enriched in genes involved in the cell cycle and cell stress response (**Fig. 2g–i and Extended Data Fig. 6g,h and 7d**) including *Cdkn1a, Ccnd1*, and Gadd45 family members *Gadd45a, Gadd45b, and Gadd45g*. While Cycling DAOs did not strongly map to any one population of mature oligodendrocytes in EAE (**Fig. 2l and Extended Data Fig. 8c**), they were highly similar to the previously described DAO subtype “DA2” found across other disease models^18^ (**Fig. 2n–p and Extended Data Fig. 8d**). Interestingly, many genes associated with neuroprotective functions (e.g., *Slc16a1*^60^, *Kcnj10*^61,62^, *Fth1*^63^) were also downregulated in Cycling DAOs (**Extended Data Fig. 6i**). In an independent bulk RNA-seq experiment of sorted GFP-expressing mature oligodendrocytes following the same experimental paradigm (**Fig. 2a,b**), we confirmed FGF1/2-induced upregulation of the same DAO genes identified by scRNA-seq (**Extended Data Fig. 6j**).

Validating our transcriptomic findings, AAV-mediated FGF1/2 overexpression induced mGFP/β-gal+ mature oligodendrocytes to express Serpina3n protein (**Fig. 3c,d**) as well as other markers enriched in DAOs including nuclear alarmin IL33 (**Fig. 3e and Extended Data Fig. 9a**), complement protein C4b (**Fig. 3f and Extended Data Fig. 9b**), and MHC-I subunit B2m (**Fig. 3g and Extended Data Fig. 9c**). In response to FGF1/2, a subset of mature oligodendrocytes also expressed p21 (gene product of Cdkn1a; ~30% of oligodendrocytes) (**Fig. 3h,i and Extended Data Fig. 9d**) as well as the mitosis marker Ki67 (~15% of oligodendrocytes) (**Fig. 3j,k and Extended Data Fig. 9e**), consistent with expression of these genes in the Cycling DAO population by scRNA-seq (**Fig. 2h**). Paralleling oligodendrocyte Ki67 expression in vivo, we found that in culture, ~25% of FGF1/2-treated oligodendrocytes incorporated thymidine analog EdU over a 24-hour time period, supporting that mature oligodendrocytes in a DAO state are capable of cell-cycle re-entry (**Extended Data Fig. 11a,b**), as previously shown in response to FGF2^23,25,26^. Nevertheless, these oligodendrocytes do not revert to OPCs, since they do not express canonical OPC markers (**Extended Data Fig. 7h**). Of note, Ki67+ mature oligodendrocytes have also been observed in histology of MS tissue^64,65^, suggesting that a similar DAO state may also occur in human disease.

**Figure 3.**
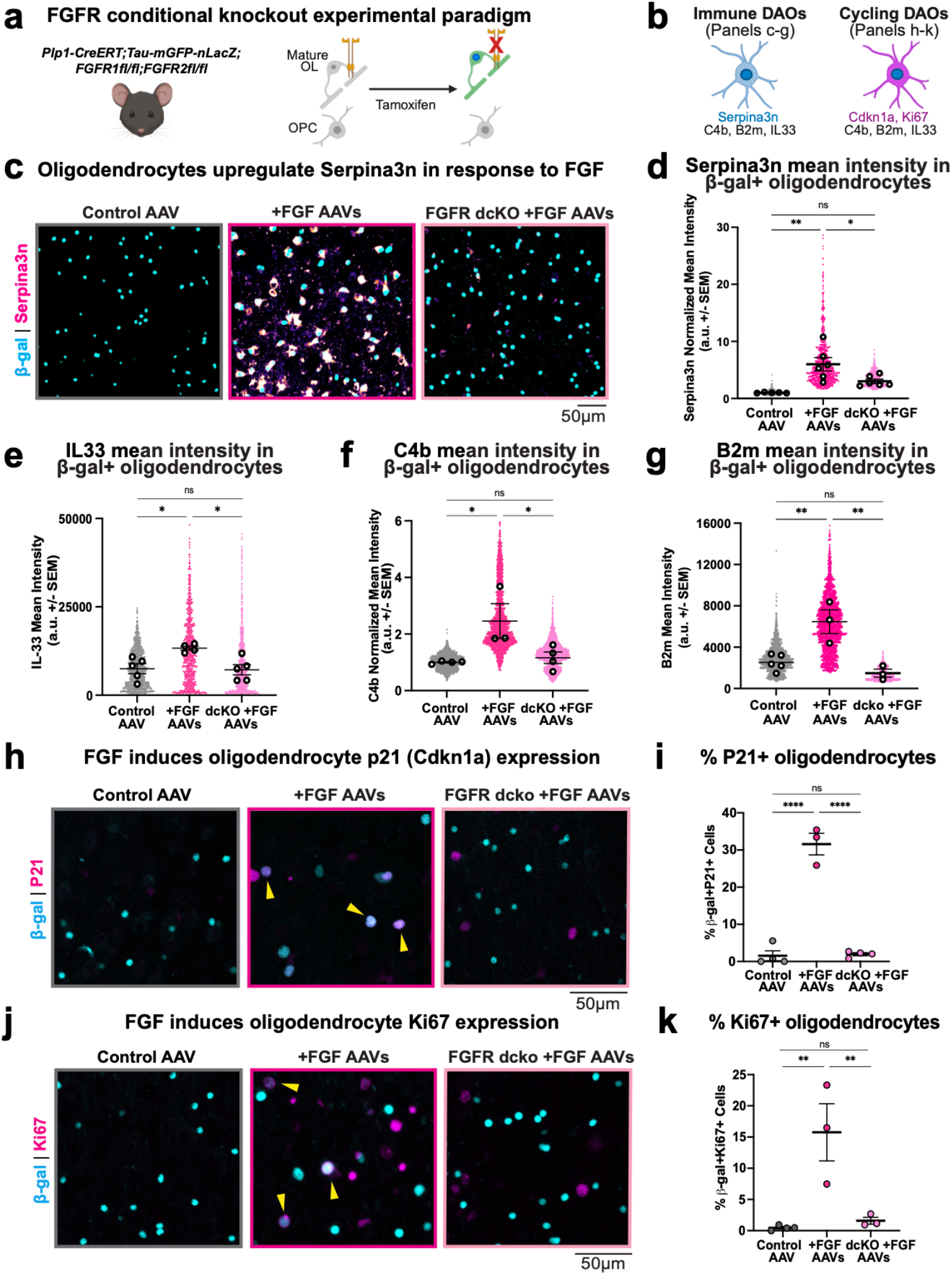
FGF-mediated DAO induction requires FGFR signaling in mature oligodendrocytes. a, Schematic introducing conditional dcKO reporter mice. b, Cartoon depicting the two major FGF-induced DAO classes, Immune DAOs and Cycling DAOs with key validated marker expression, Serpina3n and Cdkn1a/Ki67, respectively. c, Representative immunostaining for Serpina3n and β-gal in reporter mice or dcKO reporter mice injected with control-AAVs or FGF1/2-AAVs. d, Quantification of c, Serpina3n mean intensity around β-gal+ cells, normalized to control. Large black dots represent biological replicates, superplot of small colored dots are individual cell measurements. Serpina3n mean ± SEM; Control: 1.0 ± 0.03 a.u.; FGF: 6.0 ± 1.15 a.u.; 3.01 ± 0.37 a.u, n = 5-6 biological replicates. e, Quantification of IL33 mean intensity in β-gal+ cells ± SEM; Control: 7493 ± 1396 a.u.; FGF: 13322 ± 591.2 a.u.; dcKO+FGF: 7201 ± 1429 a.u, n = 4-5 biological replicates. Also see Extended Data Fig. 9a. f, Quantification of C4b mean intensity around β-gal+ cells, normalized to control ± SEM; Control: 1.0 ± 0.03 a.u.; FGF: 2.46 ± 0.61 a.u.; dcKO+FGF: 1.16 ± 0.20 a.u, n = 3-4 biological replicates. Also see Extended Data Fig. 9b. g, Quantification of B2m mean intensity around β-gal+ cells ± SEM; Control: 2523 ± 344 a.u.; FGF: 6470 ± 1154 a.u.; dcKO+FGF: 1484 ± 389.1 a.u, n = 3-4 biological replicates. Also see Extended Data Fig. 9c. h, Representative immunostaining for p21 (Cdkn1a) and β-gal in reporter mice or dcKO reporter mice injected with control-AAVs or FGF1/2-AAVs. Yellow arrowheads indicate double positive cells. i, Quantification of h. Percentage of β-gal+ cells expressing p21 +/-SEM. Control: 1.56 ± 1.34%; FGF: 31.59 ± 2.9%; dcKO+FGF: 1.97 ± 0.4%, n = 3-4 biological replicates. Also see Extended Data Fig. 9d. j, Representative immunostaining for Ki67 and β-gal in reporter mice or dcKO reporter mice injected with control-AAVs or FGF1/2-AAVs. Yellow arrowheads indicate double positive cells. k, Quantification of j. Percentage of β-gal+ cells expressing Ki67 +/-SEM. Control: 0.45 ± 0.19%; FGF: 15.76 ± 4.59%; dcKO+FGF: 1.59 ± 0.55%, n = 3-4 biological replicates. Also see Extended Data Fig. 9e. Statistical measurements (p value) were determined by 1 way ANOVA with Tukey’s multiple comparison test, p values: *p < 0.05; **p < 0.01; ***p < 0.001; ****p < 0.0001; ns = not significant.

To determine whether DAO induction was due to direct action of FGF1/2 on oligodendrocytes, we conditionally knocked out FGF receptors FGFR1 and FGFR2 (the sole FGF receptors expressed by homeostatic mature oligodendrocytes^66,67,68,55^) while simultaneously marking recombined cells (“dcKO reporter mice”: *Plp1-CreERT;Tau-mGFP-nLacZ;FGFR1fl/fl;FGFR2fl/fl*) (**Fig. 3a and Extended Data Fig. 10a,b**). Comparing dcKO reporter mice to control reporter mice following FGF1/2-AAV injection, we found that dcKO reporter mice had reduced induction of Immune DAO markers Serpina3n (**Fig. 3c,d**), IL33 (**Fig. 3e and Extended Data Fig. 9a**), C4b (**Fig. 3f and Extended Data Fig. 9b**), and B2m (**Fig. 3g and Extended Data Fig. 9c**), as well as reduced induction of Cycling DAO markers p21 and Ki67 (**Fig. 3h–k and Extended Data Fig. 9d,e**). We noted increased microglial Iba1 expression in response to FGF1/2, which was similar between FGFR dcKO mice and controls (**Extended Data Fig. 10c**). Thus, oligodendrocyte FGFR1/2 is required for DAO marker induction even in the presence of a comparable microglial response, arguing against an indirect mechanism mediated solely by altered innate immune activation. Together, these data show that FGF1/2 drives DAO induction through oligodendrocyte-autonomous FGFR1/2 signaling.

### FGF-driven DAO state induction is sufficient to trigger myelin pathology

Although DAOs have been identified in the context of demyelinating disease, it is still unclear whether DAO induction and myelin pathology are causally linked. We therefore asked whether FGF1/2-mediated DAO induction also affects myelin and oligodendrocyte homeostasis. To test this, we examined oligodendrocyte morphology and myelin architecture following FGF1/2 exposure in culture and in vivo, and asked whether these phenotypes require oligodendrocyte FGFR signaling. In culture, we found that oligodendrocytes responded to FGF1/2 with dramatic “shedding” of their stereotypical flat, MBP-rich membrane sheets (**Fig. 4a–c and Extended Data Fig. 11c,d and Supplementary Video 1**). Further, these DAO-like cells became motile and appeared to engulf their shed membranes (**Supplementary Video 2**). This dynamic behavior was accompanied by dramatic changes to the actin cytoskeleton (**Extended Data Fig. 11e,f**), with extension and retraction of actin-rich processes (**Supplementary Video 3**). This process of myelin membrane shedding and cytoskeletal rearrangement is reminiscent of how Schwann cells initiate myelin loss during peripheral nerve injury and repair, in which they activate actin polymerization to fragment their myelin sheaths before aiding in clearance of resulting myelin debris^69,70,71,72^. Oligodendrocyte morphological changes in response to FGF1/2 were similar to the changes we observed with DAO formation in EAE, including nuclear hypertrophy in culture (**Extended Data Fig. 11g**), and nuclear and somatic hypertrophy in vivo (**Fig. 4d–f**). These morphological changes were diminished in FGFR dcKO reporter mice, suggesting again that the effects of FGF signaling on oligodendrocytes are direct and cell-autonomous. Thus, in addition to altering gene and protein expression, FGF-induced DAOs enter a morphologically distinct cell state.

**Figure 4.**
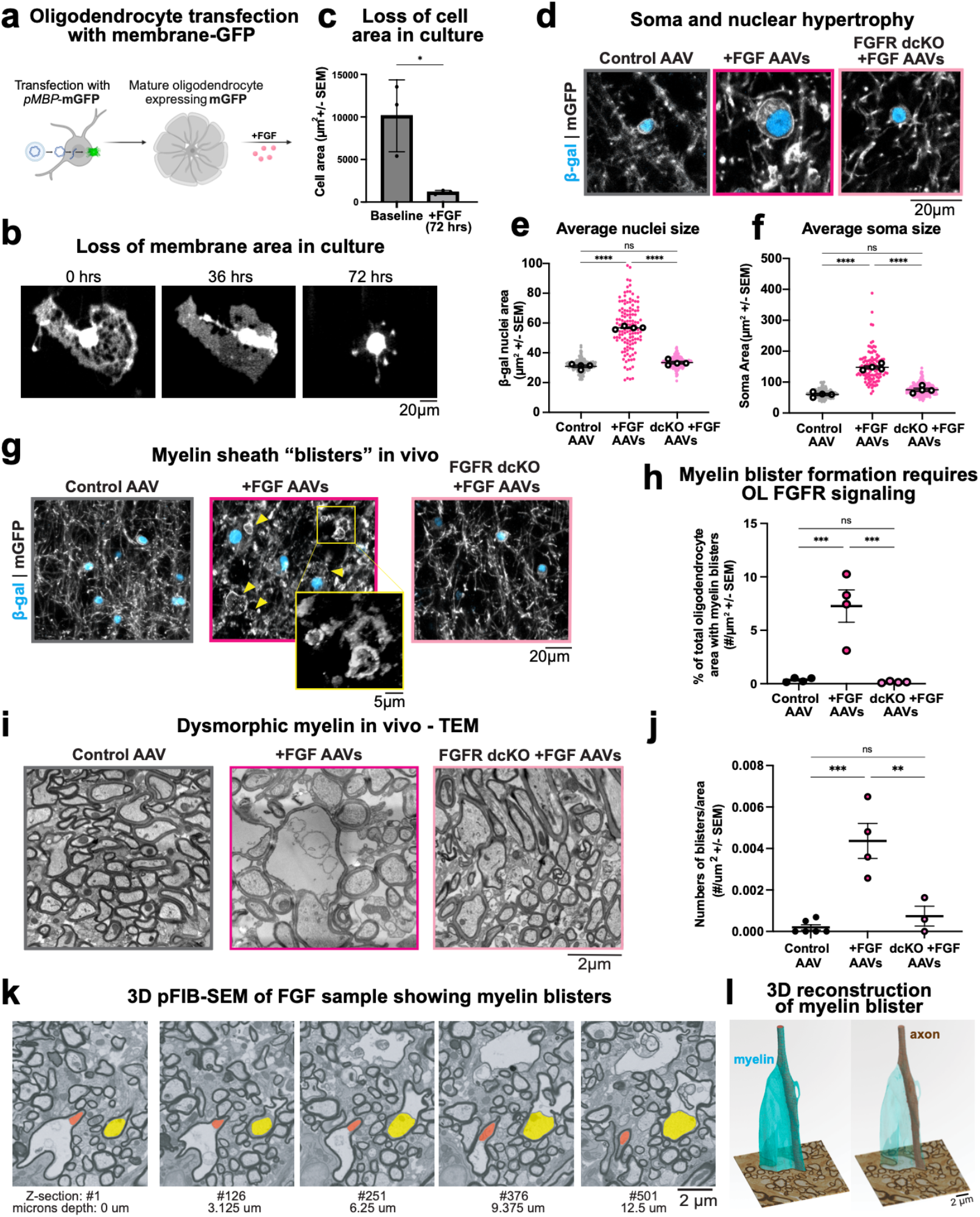
FGF-mediated DAO induction triggers oligodendrocyte and myelin pathology. a, Cartoon schematic showing transfection of cultured oligodendrocytes transfected with an MBP-promoter driven membrane-bound GFP (mGFP), differentiated for 6 days, then treated with FGF1/2. b, Representative images from timelapse imaging of FGF1/2-treated mGFP-expressing oligodendrocyte starting at day 0 of FGF treatment, showing transition from a cell with flat pancake-morphology to complete loss of myelin membrane sheets after 72hrs of treatment. Also see Supplementary Video 1. c, Quantification of change in cell area during live imaging of individual cultured oligodendrocytes over 72hrs of FGF1/2 treatment, n = 3 biological replicates. d, Representative images of oligodendrocyte nuclei and soma morphology in deep cortical layers of lineage tracing mice 14 days after infection with control-AAV (left), FGF1/2-AAV (middle), or FGF1/2-AAV into FGFR1/2 dcKO mice (right). e, Quantification of β-gal nuclei size. Large black dots represent biological replicates, superplot of small colored dots are individual cell measurements. Nuclear area ± SEM: Control 30.95 ± 0.85µm2, FGF1/2 56.82 ± 0.50µm2, dcKO FGF1/2 33.39 ± 0.87µm2, n = 4 biological replicates. See also Extended Data Fig.12a for confirmation that nuclear size measured via β-gal is equivalent to measurements made using DAPI staining. f, Quantification of GFP+ soma size. Soma area ± SEM: Control 60.02 ± 3.92µm2, FGF1/2 147.7 ± 4.94µm2, dcKO FGF1/2 75.20 ± 5.32µm2, n = 4 biological replicates. g, Representative images of myelin sheath morphology in deep cortical layers of lineage tracing mice stained with anti-GFP-488. Dysmorphic myelin “blisters” in FGF1/2 sample indicated with yellow arrowheads. Inset shows one representative blister zoomed in. h, Quantification of frequency of myelin blister formation show in g. Percentage of total oligodendrocyte area ± SEM: Control 0.35 ± 0.1%, FGF1/2 7.27 ± 1.5%, dcKO FGF1/2 0.16 ± 0.33%, n = 4 biological replicates. i, Representative electron micrographs of deep cortical layers in mice 14 days after injection with control-AAV (control, left), FGF1/2-AAV (middle), or FGF1/2-AAV into FGFR1/2 dcKO mice (right). j, Quantification of frequency of myelin blisters found in TEM. Number of blisters per um2 ± SEM: Control 0.20×10-3 ± 0.13×10-3, FGF1/2 4.37×10-3 ± 0.84×10-3, dcKO FGF1/2 0.74×10-3 ± 0.48×10-3 blisters/µm2, n = 4-6 biological replicates. k, pFIB-SEM slices (every 3.125 µm) through block of tissue from FGF1/2 sample. Pseudocolors highlighting 2 distinct axons with normal-appearing compact myelin progressing to myelin blister with space between axon and myelin sheath. Also see Extended Data Fig. 13a and Supplementary Videos 4-6. l, 3D reconstruction from pFIB-SEM of myelin blister in FGF1/2 sample shown in k. Pseudocolors highlight myelin (teal) and underlying axon (brown). Statistical measurements (p value) were determined by 1 way ANOVA with Tukey’s multiple comparison test, p values: *p < 0.05; **p < 0.01; ***p < 0.001; ****p < 0.0001; ns = not significant.

Does DAO induction by FGF1/2 also perturb myelin in vivo? Previous reports have suggested that intraventricular injection of recombinant FGF2 in rodents causes myelin disruption in adjacent regions of the brain^29,30,31^, but it is unclear whether this is due to direct action of FGF2 on oligodendrocytes. To directly test whether FGF-driven oligodendrocyte disease state induction is sufficient to produce myelin pathology in vivo, we used astrocyte-targeted AAVs to elevate FGF1/2 locally in the adult mouse brain (**Fig. 2a,b**) and combined this with oligodendrocyte-specific FGFR1/2 deletion (**Fig. 3a**). Two weeks following FGF1/2-AAV injection targeted to the cingulum of reporter mice, we found pronounced disruption to myelin sheath architecture. This included loss of sheath linearity, characterized by the formation of myelin swellings which we refer to as “blisters” (**Fig. 4g,h**). These blisters occurred within the FGF-expressing region where lineage-traced oligodendrocytes robustly induced DAO markers (**Fig. 3**), linking oligodendrocyte state change and myelin disruption in the same tissue microenvironment. Similar myelin sheath blistering has been reported in other models of demyelinating disease^73,74^ and in human MS, where it is thought to represent an early precursor to demyelination^73,75^. By co-staining for GFP (to label mGFP-positive myelin sheaths) and DAPI (to label all nuclei), we confirmed that these blisters were not aberrant myelination of cell bodies (**Extended Data Fig. 12b**). This myelin disruption phenotype was exacerbated after two months of exposure to FGF1/2, resulting in increased loss of GFP-positive myelin sheaths (**Extended Data Fig. 12c,d**).

Using transmission electron microscopy (TEM) to visualize tissue ultrastructure, myelin blisters were apparent in the FGF1/2 samples as myelinated axons with large empty spaces between the axon membrane and myelin sheath (**Fig. 4i,j**). Using plasma-focused ion beam scanning electron microscopy (pFIB-SEM) to create 3D reconstructions and trace along axons with myelin blisters, it was apparent that myelin blisters arose from normally myelinated axons (**Fig. 4k,l and Extended Data Fig. 13a–d and Supplementary Videos 4 and 5**). FGFR signaling in oligodendrocytes was necessary for this effect, as evidenced by the lack of myelin disruption and blister formation in control mice (**Extended Data Fig. 13e–g and Supplementary Video 6**) or in FGFR dcKO mice in response to FGF1/2 (**Fig. 4g–j and Extended Data Fig. 12c,d**). Because the same oligodendrocyte FGFR1/2 deletion that suppresses DAO marker induction also prevents blistering, these data support a model in which FGF/FGFR-driven DAO state switching is upstream of myelin pathology.

### Therapeutic targeting of FGF signaling attenuates DAO formation

In the EAE demyelination model, it has been shown that ablation or inhibition of FGFR signaling in oligodendrocytes ameliorates disease progression and reduces demyelination and inflammation^32,33,34,35^, but the mechanism driving this protective effect remains unclear. Is FGF signaling required for DAO induction in EAE, and could therapeutically targeting the FGF/FGFR signaling axis provide clinical benefit by preventing DAO induction? We found that *FGF2* was elevated at the time of symptom onset in EAE (**Extended Data Fig. 14a**), occurring before peak induction of DAOs^36^, consistent with FGF having an instructive role in DAO formation. As a first proof of concept, pharmacological inhibition of FGFRs using CNS-penetrant pan-FGFR inhibitor infigratinib blocked the ability of FGF1/2 to induce oligodendrocytes to enter a DAO-like state in culture (**Extended Data Fig. 14b–d**). Further, pharmacological inhibition of FGFR signaling in EAE decreased DAO formation, shown by reduced expression of DAO markers Serpina3n, B2m, and IL33 in oligodendrocytes as well as maintenance of homeostatic oligodendrocyte morphology (**Extended Data Fig. 14f–j**), and ameliorated clinical symptoms as previously shown^35^ (**Extended Data Fig. 14k**). These data suggest that attenuation of DAOs may contribute to the protective effects of FGFR inhibition in EAE, supporting pharmacologic modulation of DAO programs as a therapeutic strategy. Although systemic FGFR inhibition may affect multiple CNS and immune cell types, the reduction of DAO markers specifically within oligodendrocytes supports oligodendrocyte state modulation as a plausible component of its protective mechanism.

### Oligodendrocyte states in human MS align with FGF-induced DAOs

Finally, we asked whether human MS shows evidence consistent with involvement of FGF signaling in DAO-like states. Similar to EAE and other mouse models of demyelinating disease, oligodendrocytes in human MS undergo dramatic gene expression changes into disease-associated states^8,9,10,21,76^, and this occurs across lesion types and even in normal-appearing white matter (NAWM)^8,21^. Could FGF be a signal that triggers DAO formation in MS? In parallel with prior work showing that FGF2 levels are increased in cerebrospinal fluid (CSF) and brain tissue of MS patients^46,48,77^, we analyzed the largest snRNA-seq MS dataset to date^21^ and found that astrocytes from MS tissue expressed more *FGF2* compared to astrocytes from control brain tissue (**Fig. 5a**), specifically in NAWM (**Extended Data Fig. 15a**). We also confirmed that all oligodendrocyte subtypes identified in MS express FGFR1 and FGFR2, and expression of these receptors was increased in the major DAO population, Oligo_G, both when assessed as the average of all cells (**Fig. 5b**) and when considered at an individual sample level (**Extended Data Fig. 15b**). These findings suggest increased potential for astrocyte-to-oligodendrocyte FGF signaling in human MS.

**Figure 5.**
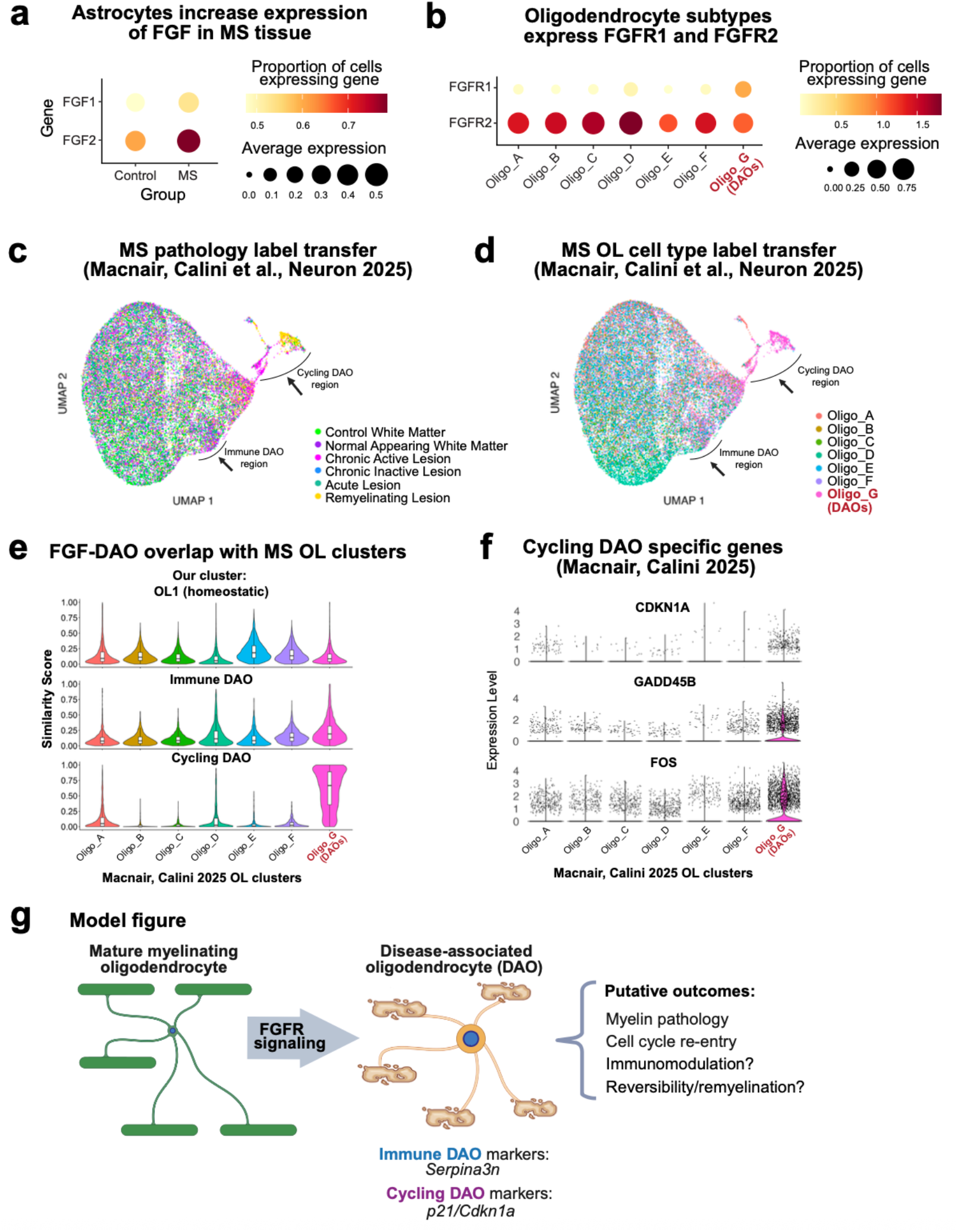
The FGF/FGFR axis is elevated in human MS, and oligodendrocyte states align with FGF-induced DAOs. a, Dot plot of FGF1 and FGF2 expression in astrocytes in MS tissue vs. control from Macnair, Calini, et al. 2025. Color scale represents the average expression of a given gene, and the size of the dot represents the proportion of cells that express that gene. b, Dot plot of FGFR1 and FGFR2 expression across oligodendrocyte clusters in MS tissue vs. control from Macnair, Calini, et al. 2025. Color scale represents the average expression of a given gene, and the size of the dot represents the proportion of cells that express that gene. c, UMAP with label transfer of pathology labels from Macnair, Calini, et al. 2025 mapped onto UMAP of mouse cells from FGF1/2 overexpression experiment (see Fig. 2). d, UMAP with label transfer of oligodendrocyte clusters from Macnair, Calini, et al. 2025 mapped onto UMAP of mouse cells from FGF1/2 overexpression experiment (see Fig, 2). e, Violin plot showing similarity score of mature oligodendrocyte clusters from MS (x-axis) to homeostatic OL1 (top), Immune FGF cluster (middle), and Cycling FGF cluster (bottom) from our FGF-expressing mice. f, Violin plots showing expression of Cycling DAO genes CDKN1A, GADD45B, and FOS across mature oligodendrocyte clusters found in MS tissue. Black dots represent expression in individual cells. g, Model Figure: FGF signaling induces mature, myelinating oligodendrocytes to become DAOs, characterized by myelin pathology and changes to cellular function—similar to human MS.

How closely do FGF1/2-induced DAOs in the mouse brain resemble disease-associated oligodendrocyte states observed in human MS? We compared the oligodendrocyte clusters from our FGF1/2 overexpression model to data from human post-mortem MS tissue to assess similarity (**Extended Data Fig. 15c**). We found oligodendrocytes in NAWM shared the most transcriptional similarity with our Immune DAOs, and oligodendrocytes in chronic active lesions and remyelinating lesions were most similar to our Cycling DAOs (**Fig. 5c**). When stratifying by the oligodendrocyte subtypes identified in the original dataset, Cycling DAOs showed the most striking similarity with subtype Oligo_G (**Fig. 5d,e**). Oligo_G was identified as the DAO population in MS tissue based on high expression of *CDKN1A* (**Fig. 5f**) and *TNFRSF12A*, and is most abundant in white matter MS samples, including demyelinated lesions and NAWM^21^. Oligo_G differentially expressed other inflammatory markers such as *CEBPD* and *ANXA2*, similar to our mouse model Immune DAOs (**Extended Data Fig. 7b,c**), as well as markers indicative of cell-cycle regulation such as *FOS, GADD45B*, and *BTG2*, similar to our mouse Cycling DAOs (**Fig. 5f and Extended Data Fig. 7b,d**). Together, these data show that MS oligodendrocyte states are concordant with FGF-induced DAOs. Combined with increased astrocyte *FGF2* and oligodendrocyte *FGFR1/2* expression in MS tissue, these results are consistent with enhanced FGF/FGFR axis engagement in human MS.

Overall, we uncover FGF signaling as a mechanism that induces disease-associated oligodendrocytes in culture and in vivo in mice, and we find that human MS exhibits oligodendrocyte states and FGF/FGFR expression patterns concordant with this program. Our data support a model in which oligodendrocytes respond to FGF by entering DAO states characterized by dramatically altered gene expression, morphology, and functionality (**Fig. 5g**), setting the stage for future studies into how manipulating this pathway affects oligodendrocyte participation in disease.

## Discussion

Disease-associated oligodendrocytes (DAOs) have emerged as a common feature of CNS pathology, yet their cellular origin, inducing signals, and functional consequences have remained unclear. Since prior work showed that OPCs can enter a disease-associated state in response to IFN-γ^13^, DAO induction could theoretically be a result of perturbed OPC differentiation. Instead, our work identifies fibroblast growth factor (FGF) signaling as a disease-relevant pathway that drives mature, myelinating oligodendrocytes to exit a homeostatic state and adopt DAO gene programs, resulting in myelin pathology. Using genetic lineage tracing in mouse EAE, we show that DAOs predominantly derive from mature oligodendrocytes rather than from progenitors, resolving a key question about DAO provenance. By combining gain- and loss-of-function manipulations of FGF1/2 (secreted ligands) and FGFR1/2 (receptors) with single-cell transcriptomics and histology, we show that FGF signaling is sufficient to induce DAOs that mirror those seen across demyelinating models and human MS, and that inhibiting FGF signaling reduces DAO induction and clinical burden in mice. Together, these data support a model in which disease-linked FGFs act directly on mature oligodendrocytes to drive DAO formation, and this state transition is sufficient to initiate myelin pathology (**Fig. 5g**).

Prior studies implicated FGF2 in demyelinating contexts by demonstrating reduced myelin gene expression in oligodendrocyte cultures^23,24,25,26,27^ and myelin disruption following exogenous FGF2 exposure in vivo^29,30,31^, and multiple reports showed that disrupting FGFR signaling is protective in mouse models of MS^32,33,34,78,79,80^. Our findings provide a unifying mechanistic explanation for these observations: FGF signaling directly drives mature oligodendrocytes into DAO states that are sufficient to initiate myelin pathology. This shifts the interpretation of FGF2 from a nonspecific “inhibitor of myelination” to an instructive cue for a conserved oligodendrocyte disease-state transition.

What do DAOs do, and are there functionally distinct subtypes? Based on our findings above, we propose that FGF-driven DAO formation is largely pathological. Nevertheless, our single-cell and histological analyses point to functional heterogeneity. We found that ectopic expression of FGF1/2 in vivo converts mature oligodendrocytes into distinct transcriptional states that closely match the major DAO classes reported across mouse demyelination models and human MS (**Fig. 2 and 5**). We refer to the two FGF-responsive DAO classes we identified as Immune DAOs and Cycling DAOs based on their dominant gene signatures. Immune DAOs upregulate complement components (e.g., *C4b*), alarmins (e.g., *Il33)*, and other genes consistent with immune modulation (e.g., *Masp1, Socs3, Cd63, Klk8)*. Pathway analysis predicts roles in inflammatory signaling, including interleukin-6 pathways linked to MS^81^. Notably, Serpina3n—a hallmark of Immune DAOs—was previously proposed to be neuroprotective via inhibition of granzyme B-mediated apoptosis^82^, but emerging data instead suggest pro-inflammatory, neurodegenerative functions^83^. These observations predict that Immune DAOs may amplify local inflammation and alter glial– immune crosstalk, which could also lead to secondary perturbations of myelin.

Cycling DAOs are distinguished by robust expression of cell cycle–related genes (e.g., *Mki67, Cdkn1a, Ccnd1*), which is surprising given that mature oligodendrocytes are classically thought to be post-mitotic. We observed EdU incorporation in cultured oligodendrocytes after FGF1/2 treatment (**Extended Data Fig. 11**), but we rarely observed cell division. Since many of these cell cycle genes also have established roles in DNA damage responses including p53-dependent cell cycle arrest (e.g., *Cdkn1a, Ccnd1*, and *Gadd45b*) and broader cellular stress responses (e.g., *Gpx1, Hmox1*, and *Txnip*), we interpret Cycling DAOs as representing a stressed state rather than productive proliferation. Given the lack of detectable oligodendrocyte SERPINA3 (human ortholog of Serpina3n) in MS, our results suggest that the major DAO population observed in human MS is more similar to Cycling DAOs. It is also possible that Immune DAOs and Cycling DAOs are phases along a continuum rather than two discrete populations. Longitudinal imaging or fate-mapping with DAO subtype-specific reporters will be required to resolve this question and to determine whether there are conditions under which Cycling DAOs could be induced to undergo cell division, as Repair Schwann cells do in the injured PNS^69,70,71,72^.

How general is FGF-driven DAO induction across CNS diseases? Elevated FGF ligands and receptors have been reported in MS^46,48,77^ as shown here, as well as other neurodegenerative disorders^84,85^, including Alzheimer’s disease^86,87^ and Parkinson’s disease^88^. In a recent transcriptomic analysis of human postmortem tissue from adult-onset leukoencephalopathy with axonal spheroids and pigmented glia (ALSP), oligodendrocyte activation was predicted to be induced by FGF signaling from reactive astrocytes^89^. These observations, together with our data, raise the possibility that FGF signaling represents a shared mechanism driving myelin vulnerability across diverse conditions. However, DAOs may also be induced by other cues. IFN-γ, produced by T cells, has been implicated in a distinct interferon-responsive DAO state^13,38^. Although less abundant than other DAO classes in some models^18,38^, we observed interferon-DAO (IFN DAO) signatures in both FGF-expressing and control animals, consistent with a broader response to injury and immune infiltration. Supporting this, high-titer AAV injections (as used here) have been shown to cause immune cell infiltration into the adult mouse brain, including CD4+ and CD8+ T cells^90^ which are known producers of IFN-γ^38,91^. Importantly, despite this shared interferon response, the Immune and Cycling DAO populations were specific to FGF1/2 expression and absent from control-AAV samples, supporting ligand-dependent induction of these two DAO classes. It remains to be determined whether IFN-γ alone is sufficient to induce mature oligodendrocytes to become IFN DAOs and whether this pathway produces myelin pathology akin to FGF1/2. We anticipate that multiple disease signals converge on DAOs, with their relative contributions varying by context.

What signaling pathways downstream of FGF1/2 drive oligodendrocytes to become DAOs? At the receptor level, our conditional knockouts demonstrated that FGFR1 and FGFR2 are required cell-autonomously in mature oligodendrocytes for DAO marker induction and myelin pathology in response to FGF1/2 (**Fig. 3, 4**). Downstream of FGFRs, canonical signaling pathways—including MAPK/Erk and PI3K/Akt—have known roles in oligodendrocytes^27,92,93,94^ that might be relevant for DAO induction. Intriguingly, experimentally hyperactivating Erk1/2 in mature oligodendrocytes causes myelin pathology similar to what we observed here in response to FGF1/2 expression^95^, though it remains untested whether this is due to DAO induction. Moreover, Erk signaling is elevated in DAOs in an Alzheimer’s disease knock-in model, and Erk inhibition partially ameliorates DAO-associated phenotypes and cognitive decline in this model^96^. Because multiple disease-linked signals activate Erk^97,98^, we hypothesize that Erk could be a “convergence node” for DAO induction, with pathway-specific inputs (including FGF1/2) shaping DAO subtype identity.

Finally, could FGF signaling represent a therapeutic target for preventing or reversing DAO formation in the multitude of demyelinating diseases where they arise? While inhibition of FGF signaling has been proposed as a potential strategy for treating MS^35,78^, it is important to consider that FGF1/2 have pleiotropic functions across the nervous system^44,99^ that increase the complexity of this approach. Both FGF1/2 induce proliferation of OPCs while inhibiting differentiation into myelinating oligodendrocytes^66,100,101^—the latter an orthogonal mechanism by which FGF signaling may be deleterious in demyelinating diseases. Beyond oligodendrocytes, FGF1/2 also promote neurogenesis and neuroprotection^84^ and regulate neuroinflammation^78,102^. Future work to uncover downstream signaling pathways regulating DAO induction may reveal more targeted strategies for preventing or reversing DAO formation in disease, which may ultimately complement orthogonal approaches to promote remyelination by OPCs in demyelinating disease.

In summary, our findings identify FGF signaling as a conserved pathway by which mature oligodendrocytes exit their homeostatic, myelinating identity to adopt disease-associated (DAO) states that actively contribute to myelin pathology. These findings also solve a long-standing mystery of why FGF signaling can perturb myelin and is deleterious in mouse models of MS^24,29,32,34^, bridging these older observations to the emerging concept of DAOs. Overall, understanding how oligodendrocytes respond to disease, as well as the mechanisms that regulate those responses, has broad implications for understanding disease mechanisms and for developing much-needed therapies for demyelinating diseases, including MS.

## Materials and Methods

### Animals

All procedures involving mice and rats were approved by the Institutional Administrative Panel on Laboratory Animal Care (APLAC) of Stanford University and followed the National Institutes of Health guidelines under animal protocols APLAC 32260 & 13226. Mice were group-housed in a Stanford University animal facility with a 12:12 hr light/dark cycle. Mouse rooms were kept between 18-23 degrees Celsius and 40-60% humidity. Mice were given ad libitum access to food and water. Mice were housed in plastic cages with disposable bedding. All mice were monitored by veterinary and animal care staff. Animals used in the study had not undergone prior procedures. *Plp1-CreERT* mice (B6.Cg-Tg(Plp1-cre/ERT)3Pop/J; Jax 005975) were obtained from Jax and bred to *Tau-mGFP;nLacZ* homozygote mice (B6;129P2-Mapttm2Arbr/J; Jax 021162), also obtained from Jax, to generate oligodendrocyte reporter mice. *FGFR1fl/fl;FGFR2f/fl* mice were a gift from David Ornitz and Juha Partanen and were bred to *Plp1-CreERT;Tau-mGFP;nLacZ* mice to obtain oligodendrocyte reporter dcKO mice. *FGFR1fl/fl;FGFR2f/fl* mice were sent for strain analysis using Transnetyx MiniMUGA Background Analysis, revealing they were generated on a mixed background, with only 60-70% on the C57BL/6 strain background. To induce recombination, tamoxifen (Sigma T5648, 100mg/kg) was administered via intraperitoneal injection for 3-5 consecutive days (3 days for lineage tracing, 5 days for all knockout experiments to ensure most complete recombination) when mice were between 8-12 weeks old. For sparse labelling experiments, mice were given a single dose of tamoxifen. Sprague-Dawley rats for primary oligodendrocyte harvest were ordered from Charles River Laboratories.

For all histology, animals were anesthetized with 2,2,2-tribromoethanol (Avertin; Sigma-Aldrich) administered by intraperitoneal injection at 200–250 mg/kg. Animals were then transcardially perfused with PBS and 4% paraformaldehyde (PFA) (made from 16% PFA, Electron Microscopy Sciences) and tissues were dissected and drop fixed in 4% PFA overnight for immunohistochemistry or Karlsson and Schultz (KS) fix for transmission electron microscopy.

### Confirming FGFR1 and FGFR2 knockout in oligodendrocytes

*Plp1-CreERT;FGFR1fl/fl;FGFR2f/fl* and *FGFR1fl/fl;FGFR2f/fl* littermates were injected with tamoxifen for 5 days as above. Three weeks later, mature oligodendrocytes were harvested from brains of 3 separate mice using the following procedure, adapted from Zuchero et al., 2015^43^: Mature mouse OLs were purified by positively immunopanning for MOG, after depletion of microglia by serial BSL1 plates. Genomic DNA and RNA were harvested from the final MOG plate after 10 washes, using RLT (Qiagen) with Beta-mercaptoethanol. DNA was PCR amplified using touchdown PCR protocol^103^ and HiFi Platinum Taq (Thermo Fischer) with 35 PCR cycles. All primer sequences are available upon request. For one replicate to confirm expression of recombined mRNA, mRNA was converted to cDNA (High Capacity RNA to cDNA kit, Thermo Fisher) and amplified with the same procedure.

### Experimental autoimmune encephalitis (EAE)

EAE was induced in male and female mice between the ages of 8 and 12 weeks. To induce EAE, mice were injected subcutaneously with 100 μg of MOG35–55 peptide (Genemed Sythesis Inc.) emulsified with Complete Freund’s Adjuvant (CFA, BD Difco Adjuvants) followed by intraperitoneal injection of 400 ng of Bordetella pertussis toxin (List Biological Laboratories Inc.) in phosphate-buffered saline (PBS) at the time of and 2 days after immunization. Control animals underwent the same treatment, excluding the MOG35-55 peptide. Brains and spinal cords were collected at the peak disease stage (one day after animals reached score 3, defined by the limp tail and complete paralysis of hind legs). Only animals with representative disease courses and clinical symptoms were used in this study. For RNAscope, WT mice were used and sacrificed to fresh-freeze spinal cord and brain at the day of symptoms onset (limp tail).

### Administration of infigratinib

The selective pan-FGFR inhibitor infigratinib (Selleck Chemicals S2183) was dissolved in 1% sodium carboxymethyl cellulose according to the manufacturer’s instructions, aliquoted and stored at −20 C. In the two treatment regimes, ~100 μl of oral doses of infigratinib (30 mg/kg) or 100 μl of vehicle (1% sodium carboxymethyl cellulose) were administered starting at day 9 until age-matched placebo controls reached peak symptoms (last dose given 24hrs before euthanasia). The study dose of 30 mg/kg of infigratinib was chosen based on prior studies of infigratinib in EAE^35^.

### Isolation of oligodendrocyte precursor cells

Primary oligodendrocyte precursors were purified by immunopanning from P5-P7 Sprague Dawley rat brains as previously described^41^. Oligodendrocyte precursors were typically seeded at a density of 150,000–250,000 cells/10 cm dish and allowed to recover for 4 days in culture before lifting cells via trypsinization and distributing for transfection, proliferation, or differentiation assays. All plasticware for culturing oligodendrocyte precursors were coated with 0.01mg/ml poly-D-lysine hydrobromide (PDL, Sigma P6407) resuspended in water. All glass coverslips for culturing oligodendrocyte precursors were coated with 0.01mg/ml PDL, which was first resuspended at 100Å~ in 150mM boric acid pH 8.4 (PDL-borate). To proliferate primary oligodendrocyte precursors, cells were cultured in serum-free defined media (DMEM-SATO base medium) supplemented with 4.2 μg/ml forskolin (Sigma-Aldrich, F6886), 10 ng/ml PDGF (Peprotech, 100-13A), 10 ng/ml CNTF (Peprotech, 450-02), and 1 ng/ml neurotrophin-3 (NT-3; Peprotech, 450-03) at 37 °C with 10% CO2. To induce differentiation, cells were switched to DMEM-SATO base media containing 4.2 μg/ml forskolin (Sigma-Aldrich, F6886), 10 ng/ml CNTF (Peprotech, 450-02), 40 ng/ml thyroid hormone (T3; Sigma-Aldrich, T6397). Cells from one individual pup considered one biological replicate.

### Oligodendrocyte culture assays

#### 96-well plate screen

Oligodendrocyte cultures were grown in differentiation media for 6 days on 96-well glass bottom culture dishes (Cellvis, P96-1.5H-N) before treatment with 10ng/ml of each of the following proteins: 0.2% BSA, FGF1 (Peprotech, 400-29A), FGF2 (Peprotech, 450-33), FGF4 (Thermo, 450-57), FGF6 (Thermo, 100-30), FGF7 (Thermo, 450-60), FGF17 (Thermo, 100-27), IL-1b (Thermo, 400-01B), IL-2 (Thermo, 400-02), IL-17 (Thermo, 210-17), IL-22 (Thermo, 210-22), IFN-γ (Thermo, 400-20), TNF-a (Thermo, 400-14), GM-CSF (Thermo, 400-23), Amyloid-Beta (Biotechne, 1428), ApoE4 (Thermo, 350-04), NRG-1 (Biotechne, 396-HB), BDNF (Thermo, 450-02), PDGF-AA (Thermo, 100-13A), EGF (Thermo, 400-25). For all FGFs, 2ng/ml heparin (Sigma H3149-10KU) was included. After 3 days, cells were fixed in 4% PFA for 15 minutes.

#### FGF1/2 treatment of oligodendrocyte cultures

Oligodendrocyte cultures were grown in differentiation media for 6 days before treatment with 10ng/ml recombinant FGF1 (Peprotech, 400-29A), 10ng/ml recombinant FGF2 (Peprotech, 450-33), and 2ng/ml heparin (Sigma H3149-10KU) or 20ng/ml BSA as a control for 3 days. For immunostaining, cells were grown on glass coverslips and fixed in 4% PFA for 15 minutes. For bulk RNA-seq, cells were grown on 6-well plastic dishes and collected in buffer RLT (Qiagen, 74134). For EdU assay, cells were grown on glass coverslips and treated with 10μM EdU (Thermo C10640) 48 hours after FGF1/2 treatment and fixed 24 hours after. For proteomics, oligodendrocytes were grown on 10cm dishes, cell lysate was collected by scraping plate in 300µl RIPA buffer (Thermo 89900) with protease/phosphatase inhibitors (Thermo 78440). Lysate was spun for 10min at 4C and supernatant collected and stored at – 80C.

#### Replating for reversibility experiment

Oligodendrocytes were grown in differentiation media for 6 days on 6-well plastic dishes before treatment with FGF1/2 for 72 hours as detailed above. Cells were then lifted from the culture dish using 0.05% Trypsin diluted in EBSS and centrifuged at 100 x g for 10 minutes to pellet. Cells were then resuspended in differentiation media and plated on glass coverslips for imaging or 6-well dishes for bulk RNA-seq.

#### Live-cell imaging of primary oligodendrocytes responding to FGF treatment

Oligodendrocytes were grown in 8-well chamber dishes (Ibidi 80827) or 35mm dishes with No. 1.5 glass coverslips (MatTek Corporation P35G-1.5-20-C) coated with PDL-borate for live imaging. Media was replaced with FluoroBrite DMEM-SATO (made with Fisher Scientific A1896701) supplemented with differentiation factors for 2 h at 37 °C, 10% CO2 before imaging. Time-lapse imaging of cells was performed on an Opterra II Multipoint Swept Field Confocal outfitted with a humidified, temperature-controlled microscope enclosure (Okolab microscope enclosure, H201-Temperature Unit (set to 37 °C), CO2 controller (set to 10% CO2), HM-Active Vibration Free Humidity Controller with humidity sensor and temperature-controlled tube). Imaging was performed using the 20x/0.75 NA objective and Perfect Focus to prevent z-plane drift during imaging. Time-lapse sequences of live cells stained with Memglow-700 (Cytoskeleton, Inc) or expressing mClover-caax were acquired every 15min. One image in a region away from cells was also imaged to be used for background subtraction to correct for uneven illumination. Lattice lightsheet images of live oligodendrocytes expressing LifeAct-mRuby3 and soluble mClover3 were acquired on a Zeiss Lattice Lightsheet 7 with a full-stage incubation chamber set to 37 °C and 10% CO2. Imaging was performed using a 13.3x/0.4 NA excitation lens, 44.83x/1.0 detection lens and pco.edge 4.2 camera every 5 min. ZEN 3.4 was used for acquisition and deskewing of acquired volumes.

### Bulk RNA-seq of cultured oligodendrocytes

RNA from cells collected in buffer RLT was extracted using RNeasy Plus Mini Kit (Qiagen 74134) and eluted in 30µl water. RNA quality and quantity were assessed using Agilent 2100 Bioanalyzer system, samples with high RNA integrity numbers (>8) were used for further analysis. KAPA/Roche Stranded mRNA-Seq kit KK8420 (Cat # 07962193001) was used to prepare poly-A enriched mRNA libraries. Samples were sequenced using Illumina Hiseq 2 x 75 platform at Stanford Genomics Center. Raw FASTQ files were trimmed using Trimmomatic (v0.32) and paired reads were aligned to the rat reference genome (Rnor_6.0) using Salmon (v1.4.0) which produced transcript-level quantifications in transcripts per million (TPM). Transcript-level quantifications were imported into R using the tximport package. Counts were summarized to the gene level based on Ensembl gene annotations and differential expression analysis was performed using DESeq2.

### Mass Spectrometry of Cultured Oligodendrocytes

10µg of protein from cultured oligodendrocyte samples were resuspended to a final volume of 40µl in PBS with 2% SDS then mixed with 80µl of SP2 binding solution (75% EtOH, 3.75% Formic Acid) and hydrophobic magnetic beads (1:1 mixture of Cytiva Sera-Mag beads, cat# 45152105050250 and 65152105050250). After incubation for 15 minutes with shaking (750 RPM), samples were applied to a multiscreen HTS-HV 0.45-µm 96-well filter plate with PVDF membrane (Merck Millipore) and centrifuged at 1000g for 2 minutes to remove unbound material. Samples were then washed 4x with 70% ethanol. Next, the digestion buffer composed of 100 mM HEPES pH 8, 1.25 mM TCEP, 5 mM chloroacetamide and trypsin (Sequencing Grade Modified Trypsin, Promega) was added to the bound protein in the filter plate. The ratio of trypsin to protein was 1:25 w/w. Trypsin was thawed and added to the ice-cold digestion buffer just before addition to the precipitates. Tryptic digestion was carried out overnight at 37C under mild shaking (500 rpm). After digestion, samples were eluted, first by centrifugation (1000g, 1 minute) and then by addition of 2% DMSO (10ul) followed by centrifugation. The eluted peptides were then dried using a vacuum concentrator. Each sample was then resuspended in 10µl of water before TMT labeling. For all samples, 10µl of TMT10 reagents at a concentration of 7.3 µg/µl in acetonitrile were added to each well. The labeling reaction was carried out for 1 hr at room temperature under shaking (750rpm) before being quenched by 5 µl of 5% hydroxylamine for 15 min. Samples belonging to the same TMT experiment were then pooled and dried using a vacuum concentrator. Pooled TMT-labeled samples were then resuspended with 250 μL OASIS Buffer A (0.05% FA in MS grade H2O) and then resuspended and acidified using 15% FA until at pH 2. Samples were then desalted using a vacuum manifold and an Oasis PRiME HLB 96-well µElution Plate (Waters, 3 mg Sorbent per well). Desalted and dried samples were subjected to high-pH reversed-phase fractionation using Pierce High pH Reversed-Phase Peptide Fractionation columns (cat# 84868). Columns were conditioned with 100% MeOH, 100% ACN, and 20 mM ammonium formate buffer (pH 10). Samples were dissolved in 250 μL of dilution buffer (20 mM ammonium formate, pH 10) and loaded onto conditioned columns. Following sample loading and washing, peptides were eluted using a step gradient of 2-100% ACN in 20 mM ammonium formate across 48 fractions, which were then pooled into 24 fraction groups for analysis (fraction 1 recombined with fraction 25, etc). Fractions were dried by vacuum centrifugation, resuspended in 0.2% formic acid, and loaded for LC-MS/MS analysis.

#### Mass Spectrometry Data Acquisition and Processing

Samples were analyzed by LC-MS/MS using a Dionex UltiMate 3000 RSLCnano System coupled to a Thermo Scientific™ Q Exactive Orbitrap mass spectrometer with Nanospray Flex source. The HFX was operated in positive ion mode with a spray voltage of 1.5 kV and a capillary temperature of 275 °C. Fractionated samples were reconstituted in 0.2% formic acid in water and were loaded to a trap column (Thermo Scientific™ Acclaim™ PepMap™ C18 column, 2 cm x 100 µm ID, 5 µm) and separated on an analytical column (μPAC™ Neo HPLC Column (110 cm)). Solvent A is 0.1% formic acid in water and solvent B is 100% acetonitrile in water with 0.1% formic acid. In a 185 minute method, the gradient was ramped from 5% to 7% at the 10 minute mark, 28% at the 156.5 minute mark, 95% at the 161 to 170 minute mark, then back down to 5% at the 170.1 minute mark for the remainder of the method. TMT labeled peptides were analyzed by data-dependent acquisition mode. For Full MS Scan, resolution was set to 120,000, AGC target of 3e6, Maximum IT of 50 ms, and a scan range of 350 to 1500 m/z. For dd-MS2, Resolution was set to 45,000, AGC target of 1e5, Maximum IT of 120 ms, Loop count of 15, isolation window at 0.7 m/z, fixed first mass of 110 m/z, and NCE of 31. Raw mass spectrometry files were processed using a standardized computational pipeline. Raw files were first converted from .raw format to .mzML format using MSConvertGUI with default parameters and the peakPicking and zeroSamples filters set. Peptide identification and quantification were performed using FragPipe software (v22.0) with the TMT10 workflow. Protein identification was performed against the rat reference proteome database (UniProt, downloaded on May 11, 2025). Fixed modifications were carbamidomethyl on cysteines and TMT10 on lysine. Variable modifications included acetylation on the protein N-terminus, oxidation of methionine, and TMT10 on peptide N-termini. Protein-level quantification was achieved by grouping peptides to their corresponding proteins. No normalization or log2 transformation was applied during the initial processing, preserving raw abundance relationships for downstream statistical analysis. The resultant abundance_protein_None.tsv and protein.tsv files were used in downstream data analysis. Abundance analysis and fold change calculations were performed using custom R scripts. In brief, identified proteins were first filtered for sufficient coverage at the peptide level (psm >1) and replicate level (observed across all TMT10 mass channels). Median normalization was then applied to all samples to account for non-biological experimental variability in the theoretical 10µg input amount. First, intensity values were Log2 transformed then the median was calculated across all samples (median_all) and within each sample (median_sample). Each intensity value (intensity_log2) was then scaled accordingly (median_all – median_sample + intensity_log2). For each identified protein, Log2 fold change and the significance (t-test) of change was calculated.

### Transfection of OPCs

Rat OPCs were trypsinized and dissociated from tissue culture dishes and centrifuged at 90 × g for 10 min. 250,000 OPCs were gently resuspended in 20 μl of nucleofector solution (Lonza P3 Primary Cell 4D-Nucleofector V4XP-3032) with 400 ng of pMBP-EGFP-caax plasmid (Addgene #190155)^104^. Cells were loaded into a 16-well cuvette and electroporated in a Lonza 4D-Nucleofector X Unit (AAF-1003X) assembled with a 4D-Nucleofector Core Unit (AAF-1002B) using pulse code DC-218. Electroporated cells rested for 10 min at room temperature. 80 μl of antibiotic free DMEM-SATO media was added to each cuvette and cells were gently triturated. Each well of 250k cells was distributed onto No. 1 glass coverslips coated with PDL-borate for differentiation timepoints and technical replicates (up to six coverslips from a single transfection). Each coverslip was half-fed with freshly supplemented DMEM-SATO media every two to three days.

Cloning of FGF1 and FGF2 overexpression AAVs The pGFAabc1d-tdTomato construct was purchased from AddGene (#44332). The pGFAabc1d-FGF1 and FGF2 constructs were assembled by InFusion cloning (Takara Bio) to anneal two DNA fragments in a pAAV vector backbone. The astrocyte-specific gfaABC1D promoter that has been widely-used in vivo^54^. The parent plasmid (pGFAabc1d-tdTomato), which contains the GFAP promoter in a pAAV vector backbone, was linearized using the XhoI and XbaI restriction sites. Mouse FGF1 and FGF2 fragments were codon optimized for mouse oligodendrocyte expression using GenScript, and synthesized by TWIST biosciences (San Francisco, CA). FGF1/2 both lack a signal peptide and are secreted from cells by an unconventional mechanism^105^. A constitutively secreted form of FGF1 was previously made by fusing the signal peptide from FGF4 to the coding sequence of FGF1; this results in forced secretion of functional FGF1 from mammalian cell. We copied this design to express secreted forms of FGF1 from astrocytes. We found FGF2 secretion was higher from cultured astrocytes without the signal peptide (data not shown), and thus the FGF2 construct does not contain a signal peptide. All constructs are <4700 bp (AAV packaging limit). We used ELISAs to confirm that both result in release of soluble FGF1 (8 ng/mL) or FGF2 (12 pg/mL) into culture media from AAV-infected primary rat astrocytes, and functional assays confirmed that this conditioned media is sufficient to induce mature OLs to downregulate mature OL marker MBP and upregulate DAO marker genes, similar to using recombinant FGF1/2. Serotype AAV-2/5 was prepared by the Stanford Neuroscience Gene Vector and Virus Core, and stored at −80 °C until use.

### Isolation and treatment of astrocytes

Astrocytes were purified by immunopanning from P6 Sprague Dawley rat forebrains and cultured as previously described^106^. Cortices were blunt dissected and enzymatically digested using papain at 37°C and 10% CO_2_. Tissue was then mechanically triturated with a 5 mL serological pipette at room temperature to generate a single-cell suspension. The suspension was filtered in a 70 μm nitex filter and negatively panned for microglia (CD45; BD Pharmingen 554875 for mouse, BD Pharmingen 553076 for rat), endothelial cells (BSL I, Vector Labs L-1100), and oligodendrocyte lineage cells (O4 hybridoma, in house) followed by positive panning for astrocytes (for mouse: HepaCAM, R&D Systems MAB4108; for rat: ITGB5, Thermo, 14-0497-80). Astrocytes were removed from the final positive selection plate by brief digestion with 0.025% trypsin and plated on poly-d-lysine coated tissue culture plates. Astrocytes were cultured in defined, serum-free medium containing 50% neurobasal, 50% DMEM, 100 U/mL penicillin, 100 μg/mL streptomycin, 1 mM sodium pyruvate, 292 μg/mL L-glutamine, 1× SATO, 5 μg/mL of N-acetyl cysteine, and 5ng/mL HBEGF (Peptrotech, 100-47). For conditioned media experiments, astrocytes were cultured for 7-14 days in defined serum-free media on 10cm culture plates before the addition of AAVs. Plates from the same prep received either an AAV containing tdTomato or AAVs containing FGF1 and FGF2 (described above). 1 week after infection, conditioned media was collected and concentrated in a 10kD spin column (Cytiva 28-9323-60) for ELISA analysis of FGF secretion using FGF1 ELISA kit (Abcam ab223587) or FGF2 Quantikine ELISA kit (R&D systems, MFB00).

### Stereotactic AAV injections

High titer AAVs (described above, 1×10^12^-1×10^14^ vg/mL) serotype AAV-2/5 were prepared by the Stanford Neuroscience Gene Vector and Virus Core, and stored at −80 °C until use. AAVs to express tdTomato control or tdTomato together with FGF1 and FGF2 were injected bilaterally into the cingulum of isofluorane-anesthetized reporter mice (1.75μL hightiter AAV; from bregma: x=1mm, y=1mm, z=1.4mm) using Hamilton syringe (Hamilton 80308, 25mm) and syringe pump set to 200nl/mL and 2min waiting period before and after AAV injections to allow tissue equilibration. Expression can spread into the contralateral hemisphere through the corpus callosum, so we did not use contralateral injections as controls.

### Immunostaining

Primary antibodies used in this study were as follows: rat anti-MBP (abcam ab7349, 1:10000 cultured cells, 1:1000 in vivo), goat anti-Serpina3n (Bio-Techne AF4709, 1:500), goat anti-IL33 (Bio-Techne AF3626, 1:500), rabbit anti-Vimentin (abcam ab92547, 1:1000), rabbit anti-beta2 microglobulin (abcam ab75853, 1:500), chicken anti-Beta-Galactosidase, LacZ (Aves Labs BGL1010, 1:1000), chicken anti-GFP (abcam ab13970, 1:2000), rabbit anti-GFP (Invitrogen A-11122, 1:2000), rabbit anti-Ki67 (Thermo PA519462, 1:1000), rabbit anti-P21 (abcam 188224, 1:500), rat anti-complement C4 (Invitrogen MA1-40047, 1:100), rabbit anti-Olig2 (Sigma AB9610 1:500), rabbit anti-Iba1 (Wako Chemicals 019-19741, 1:1000), rabbit anti-ASPA (GeneTex GTX113389, 1:1000, requires antigen retrieval), mouse anti-APC (CC1) (EMD Millipore OP80-100UG, 1:50). Secondary antibodies used in this study were highly cross-absorbed Alexa Fluor 488-, 594-, or 647-labeled secondary antibodies (Thermo Fisher) or Dylight 405 (Jackson ImmunoResearch) all used at 1:1000 dilution for cultured cells and 1:500 dilution in tissue.

#### Cells

Cell media was removed, and coverslips were treated with 4% PFA for 15 min at RT, followed by three washes with 1xPBS and permeabilization in 0.1% Triton X-100 in PBS for 3 min at RT. Prior to staining, cells were incubated in a blocking solution of 3% BSA in PBS for 20 min at RT. Then, primary antibodies were added in a 3% BSA solution for overnight incubation at 4 C. On the following day, the primary antibody solution was rinsed off with three washes of PBS, and then incubated with secondary antibodies in 3% BSA for 1 hr at RT. After three washes with PBS, CellMask Blue stain (1:1000) was incubated to stain all cells for 15 min at RT, followed by three additional rounds of washing with PBS. Stained cells were mounted onto microscope slides (Fisher Scientific 12-550-143) in Fluoromount G (SouthernBioTech, 0100-20). Cells were imaged by widefield epifluorescence with a Zeiss Axio Observer Z1 using the Plan-Apo 20x/0.8 NA objective. Images were acquired blinded to condition with identical illumination and acquisition conditions per biological replicate.

#### Tissue

For spinal cords, tissue was drop fixed in 4% paraformaldehyde overnight, washed with PBS, and then dropped into 30% sucrose before embedding in OCT. Fixed spinal cords were sectioned on a cryostat (CM3050S, Leica Microsystems) into 30 μm sections mounted onto Superfrost Plus (VWR) microscope slides and stored at −80 °C until subsequent use. Prior to staining, slides were warmed at 34 °C for 30 minutes, sections were circled with a hydrophobic pen (ThermoScientific 008899) and dried for another 5 minutes. Sections were blocked with 10% donkey serum in PBST (0.3% Triton X-100 in PBS) for 1 hour and then primary antibodies diluted in 1% donkey serum in PBST were applied (dilutions specified in Antibodies section) and incubated at 4 °C overnight. The next day, primary antibody was rinsed with 3 ×20 minute washes in PBS. Secondary antibodies diluted in 1% donkey serum in PBST were incubated for 2 hr at RT. Secondary antibody was washed 3 ×20 minutes in PBS. Samples were mounted in Vectashield with DAPI (Vector, H-1200) using a coverslip.

For brains, tissue was drop fixed in 4% paraformaldehyde for overnight then washed with PBS and sectioned to 40 μm on a vibratome. Free-floating sections were blocked in 3% donkey serum in PBST for 1 hour then incubated in primary antibodies diluted in 1% donkey serum in PBST overnight at 4 °C. Sections were washed 3x in PBS and then incubated in secondary antibodies diluted in 1% donkey serum in PBST overnight at 4 °C. Secondary antibody was washed 3 ×20 minutes in PBS and samples were mounted in Vectashield with or without DAPI (Vector, H-1200) using a coverslip.

For staining brain tissue for known markers of mature oligodendrocytes (CC1^107^ and ASPA^108^). The following modifications to the above protocol were made. For CC1 (APC) staining: primary antibody for 5-7days at 4°C. For ASPA staining: antigen retrieval with 40min at 60°C (at 15min at 95°C) in 1X citrate buffer (Sigma-Aldrich C9999), wash three times in PBS, then block and proceed with staining as usual.

RNA Fluorescence in situ hybridization (RNAscope)

Mice were euthanized by decapitation with isoflurane anesthesia to allow for rapid collection of tissues for RNAscope. Brain and spinal cord were dissected and immediately frozen in OCT (Thermo Fisher Scientific, 23-730-571) on dry ice. Tissues were kept at −80 °C until sectioning. For tissue sectioning, 10–12 μm thick spinal cord cross sections or coronal brain sections were collected with a Lecia CM3050 S cryostat preset to –20 °C. Sections were immediately transferred to cold Superfrost Plus (VWR, 48311-703) microscopy slides and stored at –80 °C until RNAscope assay was performed. RNAscope multiplex fluorescence kit v2 (ACDbio) assays were completed according to the manufacturer’s sample preparation manuals. Multiplex fluorescence kit v2 (ACDbio, 323270) and the HybEZTM II Hybridization system (ACDbio, 321710) were used in all RNAscope assays. Fresh frozen tissue slices were fixed for 15 minutes in 4% PFA (EMS, 15710) and serially dehydrated in ethanol washes immediately before assaying. After dehydration steps, samples were dried at room temperature for 20 minutes and then treated with pretreat IV (ACDbio, 320842) for 30 minutes. RNAscope probes were hybridized to target mRNAs, and the samples were treated with hybridization reagents according to the kit manual, then counterstained with DAPI and mounted with Prolong Gold antifade mount (Invitrogen, 10144). The following probes from ACDbio were used in this study: mFgf1-codon-O1-C1 (1256891-C1), mFgf2-codon-O1-C1 (1256901-C1), Mm-Fgf1 (466661), Mm-Fgf2 (316851).

Slides were imaged with a Zeiss LSM 800 laser scanning confocal microscope using Zeiss Zen Blue software. A 20x objective was used to obtain Z-stack images. Specimens were blinded during imaging and analysis.

### Image Analysis and Quantification

For IHC analysis of brain and spinal cord tissue, fixed and stained tissue sections were imaged with the researcher blind to condition using a Zeiss LSM 800 laser scanning confocal microscope with the Plan-Apochromat 20x objective. Images were acquired using the Zeiss Zen Blue software and quantified. 4–6 animals per condition were analyzed and the N number is specified in the Figures and Figure Legends. For EAE spinal cord imaging, at least 3 sections per animal were imaged in white matter at lesion-adjacent areas. See supplemental figure S2d for examples of imaged regions. For AAV brain tissue imaging, regions of interest were deep cortex and corpus callosum where tdTomato was present, confirming successful AAV injection and construct expression. For quantification of DAO marker expression, maximum intensity projections of each z-stack and background subtraction were done in Fiji (ImageJ, NIH). Images were subsequently exported to QuPath^109^ for automated cell detection and intensity quantification. First, β-gal+ nuclei were detected and nuclei ROI were expanded 3µm (or no expansion for nuclear markers e.g. P21, Ki67, IL33) to incorporate cell soma area. All nuclei selections were checked by a researcher blinded to condition to ensure single nuclei were selected. Images were then batch processed for intensity of DAO marker staining (e.g. Serpina3n) within each individual cell. Data were exported and processed to plot average mean intensity of all cells within an image, and replicate images were averaged as the mean intensity for that animal. For % expression analysis, intensity threshold was chosen based on visual inspection of images to determine intensity of cells that stain positive vs. negative for each respective marker, and the same threshold was applied to all cells within the same round of staining/imaging.

For RNAscope analysis, similar imaging and analysis pipeline as above was used, however images were also processed with 1pixel Gaussian Blur filter in Fiji (to increase SNR of RNA spots) before being exported to QuPath. Nuclei were detected with DAPI. Individual RNA transcripts within a 10 µm radius of each nuclei were counted with QuPath. Data were exported and processed to calculate average number of transcripts per unit area of cell.

### Preparation of single cell suspension for flow cytometry

Reporter mice previously injected with AAVs (described above) were anesthetized as previously described and perfused with cold PBS. Brains were removed, sliced to 2mm thick around the injection site on a coronal brain matrix, then injection area was removed using 2-3 punches of a 2mm diameter tissue punch. Tissue was further chopped in 2ml Hibernate A solution (HACA, Brainbits). Tissue pieces were collected in a 2ml round bottom tube and centrifuged 300g for 2 min. Supernatant was removed and 2 ml pre-warmed Enzyme mix 1 from Neural Tissue Dissociation Kit (P) (Miltenyi 130-092-628) with 5 μg/ml actinomycin D (Sigma-Aldrich A1410) was added to each tissue sample. Samples were incubated on a rotator for 15 min at 37°C in 2ml tube. Enzyme mix 2 from Neural Tissue Dissociation Kit (P) was added, sample transferred to a 15ml tube and gently triturated 30x with wide-orifice pipette tips. Samples were incubated for another 10 minutes, triturated 30x with regular pipette tips, incubated for 10 minutes, then triturated with 10ml ice-cold Hibernate A Low Fluorescence medium (HALF, Brainbits). Cells were filtered through a 70 μm filter (Miltenyi 130-098-462), then centrifuged 300g for 10 minutes at 4°C. Cell pellets were then resuspended in Debris Removal solution (Miltenyi 130-109-398) following manufacturer’s instructions. After debris removal, pellets were resuspended in 500µl HALF with 5% FBS. Cells were stained with Calcein Violet AM (Thermo C34858) and Viability Stain 700 (BD 564997) at 1ug/ml for 15 minutes, washed with 2ml HALF with 5% FBS, spun 500g for 5 minutes, and resuspended in 400µl HALF with 5% FBS. Flow cytometry was used to sort and collect GFP+ oligodendrocyte live single-cell suspensions for the scRNA-seq study.

For bulk RNA-seq, cell suspensions were immediately processed for RNA extraction by pelleting cells and resuspending in buffer RLT (Qiagen). RNA extraction was done using RNeasy Micro Kit (Qiagen 74004). RNA samples were sent to Novogene for QC, library preparation and sequencing using NovaSeq X Plus Series (PE150).

### Single cell RNA-seq library preparation and sequencing from FACS sorted cells

After sorting, cells were resuspended to 1000cells/µl in PBS and 16,500 cells were loaded onto a Chromium Next GEM Chip G. GEM generation & barcoding, post GEM–RT cleanup & cDNA amplification, and 3’ gene expression library construction were performed as described in the Chromium Single Cell 3’ Reagent Kits User Guide (10x Genomics CG000317 v3.1). Gene expression libraries were pooled together and sequenced on an Illumina NovaSeq X Plus Series flowcell. The BCL files were demultiplexed using 8 base pair 10X sample indexes and cellranger mkfastq to generate paired-end FASTQ.

Single-cell RNA-seq data analysis

#### Data pre-processing

Raw fastq data were analyzed using the CellRanger pipeline version 9.0.1 and aligned to the reference mouse genome version mm10. After generating single cell raw count matrices for each condition, they were loaded in R 4.50 and analyzed using Seurat 5.3.0. First a unique object was generated by merging together count matrices from the control (tdTomato) and treated (FGF) condition, after which low quality cells were filtered out using the following criteria: number of features > 200 and < 7000 and percentage of mitochondrial assigned counts < 5% resulting in 15,383 cells for control and 13,045 for FGF.

After selecting only high-quality cells, the count matrix was log normalized and the top 2000 variable features were calculated, data scaling was performed, and Principal Component Analysis (PCA) executed. After this step, clustering and dimensionality reduction using Uniform Manifold Approximation and Projection (UMAP) was executed and initial result visualized, revealing the necessity of performing an additional step of data integration between the 2 conditions. Following this, data were further normalized using regularized negative binomial regression (scTransform^110^), PCA was run on the SCT normalized data, and data integration was executed before clustering and dimensionality reduction using UMAP. After this step, clusters were manually inspected and annotated using known marker gene resulting in the identification of mature oligodendrocytes, microglia and OPCs.

#### Single-cell RNA-seq mature oligodendrocyte reclustering

Following major cell type identification, only mature OLs were selected for the subsequent analysis resulting in a total of 27,669 cells. After subsetting mature OLs, scTransformation, layer integration, PCA, clustering and dimensionality reduction using UMAP were executed on the data subset. Once again, clusters were inspected manually and after manual annotation 5 major clusters have been identified: OL_1, OL_2, DAO_IFN, DAO_FGF_immune (*Immune DAO*), DAO_FGF_cycling (*Cycling DAO*). Using these 5 clusters, marker genes were calculated for each one and sample composition of each cluster was computed using the FindAllMarkers function from Seurat with default settings log2fc threshold=0.1 and pvalue threshold of 0.01.

#### Functional enrichment

Functional enrichment was performed using the R package clusterProfiler using the significantly upregulated marker genes in each cluster as the input, and using them to query against the GeneOntology Biological processes dataset and the Kegg pathways dataset. Pvalue cut-off for significant categories has been set at 0.5 with Bejamini Hochberg correction for multiple tests. The results were plotted as −log10(padj) in all the figures.

#### Annotation with external mouse datasets

To determine the similarity between our cell clusters and other datasets we used the label transfer approach from Seurat^111^. Briefly the Seurat object from Falcão, van Bruggen et al., 2018 (EAE dataset, available at GSE113973) and Pandey et al., 2022 (pan-disease dataset, downloaded from Zenodo 10.5281/zenodo.6784453) were loaded into Seurat. Using the FindTransferAnchor function, the predicted identity from our dataset based on the clusters present in the external datasets was computed for each cell, and the prediction with the highest score was selected and plotted in the UMAP. For each cluster in our dataset, we plotted the average predicted score for each cell in that cluster against each cell type of interest from the external datasets as a violin plot.

To compute the gene module score, the same methods used in Pandey et al., Cell Rep, 2022 have been used here. Briefly, for each dataset analyzed, a list of upregulated and downregulated genes associated specifically to each cluster was generated. For the dataset from Pandey et al., the gene lists were obtained from the supplementary material of the paper using the DAO_fine_upregulated and DAO_fine_downregulated annotation. For the Falcão, van Bruggen et al., 2018 paper the top 100 upregulated and downregulated genes have been selected using the supplementary table from the paper containing the DifferentialExpressionOLs data using the gene ranking based on log2FC per cluster. After having selected the list of genes for each module, the score was calculated as follows: Gene expression values were log-transformed as Log2(normCount+1). For each sample, the average enrichment score was calculated as the average expression value across all the genes in the gene set. For each analysis, the average value of the control condition of the dataset was subtracted from the average of the other samples to obtain a control-centered module score. Since all the gene module scores presented in this manuscript included both upregulated and downregulated genes, a signed average score was computed by weighting all the up genes by +1 and down genes by −1 to capture both changes in a single score. OL_1 was set as the control population for this analysis for the mature OL dataset from this manuscript.

#### Human dataset analysis

To compare the mature oligodendrocyte dataset from FGF treatment and tdTomato control with human data from MS, the following strategy was applied - RNA count matrices from Macnair & Calini et al. 2025 (retrieved upon request from EGAD00001009169) were converted to M musculus (Mm10) using EnsEmbl Biomart database. Only ortholog genes with conservation score >= 90 and homology type “ortholog_one2one” between hg38 and mm10, ENSEMBL version 102, were selected for downstream analyses. The final number of genes was reduced from 59981 genes to 21259 ortholog genes in Macnair & Calini et al. 2025.

Only data from white matter OL lineage cells was used by only keeping nuclei labelled as white matter and as “Oligodendrocytes” or “OPCs + COPs”. Then the subset was downsampled to 3000 nuclei per “type_fine” label, cell type subtypes from the original publication. Prediction of the human cell types was performed on the mouse data using Seurat v5 label transfer, FindTransferAnchors (cca) with 1703 intersecting variable genes and using “lesion_type” as a reference For the gene module score, analysis of the top 100 upregulated and downregulated cluster from the mouse mature OL dataset was converted to the ortholog gene using Homologene R package. Afterwards the gene module score has been calculated using the same methods detailed above, setting as control population Oligo_D for the Macnair & Calini dataset.

### High pressure freezing with freeze substitution for electron microscopy

Mice were anesthetized and perfused with ice cold PBS then 4% PFA. Brains were immediately and carefully dissected out, placed on an epifluorescence microscope to confirm tdTomato+ area, then 1.5mm tissue punch was taken of the tdTomato+ area including cortex and corpus callosum. Tissue punches were then post-fixed in cold KS-fixative at 4 °C. 100 µm thick vibratome sections from previously fixed brain material were stained by using an osmium-thiocarbohydrazide-osmium (OTO) method^112,113^ in combination with microwave-assisted processing, followed by high pressure freezing and freeze substitution (HPF-FS), as previous described^114^. Briefly, samples were OTO stained, incubated with 2% aqueous uranyl acetate overnight, then subjected to HPF, followed by super quick FS^115^ with 4% osmium tetroxide, 0.1% uranyl acetate and 5% ddH2O in acetone; samples were infiltrated in increasing epon gradient then embedded and polymerized at 60 °C for 48 hours.

Transmission electron microscopy

Ultrathin sections, 90 nm, were cut using a Leica UC6 ultramicrotome (Leica Microsystems, Wetzlar, Germany) and collected onto formvar-coated copper-rhodium backed slot grids. Due to *en bloc* contrast from OTO staining, no post-stain was necessary prior to TEM imaging. Thick sections, 300 nm, were collected onto PTFE glass slide wells and stained with toluidine blue as a reference in light microscopy of the tissue. Thin sections were imaged using a Tecnai 12 120kV TEM (FEI, Hillsboro, OR, USA) and data recorded using a Gatan Rio16 CMOS camera with GWS software (Gatan Inc., Pleasanton, CA, USA) or SerialEM (University of Colorado Boulder). Samples that did not contain clear corpus callosum and adjacent cortex were excluded.

3D electron microscopy and reconstruction using plasma-focused ion beam scanning electron microscopy (pFIB-SEM)

#### Acquisition

Epon-embedded tissue blocks (see “High pressure freezing with freeze substitution” section, above) were attached to an aluminum SEM stub by superglue, trimmed to about 1 mm square, and the sides were painted with conductive graphene carbon paint (Electron Microscopy Sciences, Hatfield, PA). Serial imaging was performed with a Thermo Scientific Helios 5 Hydra Plasma-focused ion beam scanning electron microscope in the Central Microscopy Facility at the Marine Biological Laboratory, Woods Hole, MA. A standard spin mill procedure in the ASV software using oxygen plasma was used, similar to that described previously^116^ except that the number of spin angles was 5. The imaging was done using the backscatter detector, with 3 kV, 0.4 nAmp current, 1 microsecond dwell time at 10 nm per pixel.

#### Reconstructions and visualization

##### (A) Volume Preprocessing

The original TIFF volume stack from FIB-SEM imaging was loaded and resampled for consistent slice thickness. Two smaller volumes through the stack were extracted around two main regions of interest containing myelin blisters. The stacks were then independently processed and visualized using custom-written MATLAB scripts.

##### (B) Segmentation

Each stack was subjected to slice-by-slice intensity normalization followed by thresholding to detect dark pixels which were identified as myelin. Dilation was performed to fill small gaps on a per-frame basis and the final connected components of the largest object was labeled as myelin. Axon identification was performed by filling the interior of each ‘hole’ in the myelin, followed by a closing operation. The labeled volume was exported to the MATLAB volumeSegmenter app and each volume was manually corrected.

##### (C) Visualization

The final rescaled intensity volumes and their corresponding indices of myelin and axon labels were saved as .mat files for video generation. A custom script for myelin visualization was used to create a multi-stage 3D animation that shows views through the volume followed by rotations, and multi-layer reveals of the myelin bleb and axon.

## Data analysis and statistics

All statistical analyses were done using GraphPad Prism 10. Statistical details of all experiments can be found in the figure legends. Additional details of experimental analysis can be found under method descriptions. We pre-determined that using 3–5 biological replicates for culture or in vivo experiments were sufficient based on previously published studies^43,117^. Reported mean always represents mean of independent biological replicates. Unless otherwise stated in figure legend: for all data comparing three groups, statistical measurements (*p* value) were determined by 1 way ANOVA with Tukey’s multiple comparison test. For all data comparing only two groups unpaired two-tailed t-tests were used, or for cell culture IHC where expected directionality of the effect was predetermined paired one-tailed t-test were used to determine significance with a cut off of *p* = 0.05.

## Data availability

The data generated in this study are provided in the Supplementary Information and Source Data files. Source data are provided with this paper. The RNA sequencing data will be deposited in GEO at the time of publication. Proteomics datasets will be deposited at PRIDE prior to publication. All AAV constructs developed for this study will be deposited at Addgene and made available at the time of publication (AAV_pGFAabc1d-FGF1, AAV_pGFAabc1d-FGF2). Step-by-step protocols are available from the corresponding or first authors upon request.

## Code availability

Detailed code with step by step explanation of each analysis step can be found on GitHub.

Correspondence and requests for all other materials should be addressed to M.H.C., M.K.W., or J.B.Z. Source data are provided with this paper.

## End Matter

### Author Contributions and Notes

M.W., M.H.C, and J.B.Z. conceptualized the project. M.W., M.H.C., C.M.B., M.S., D.M.J., and J.B.Z. developed methodologies for experiments. M.W., M.H.C., J.G., C.M.B., M.S., P.P.H., G.T.-L., V.V.R., D.M.J., M.R.T., and J.B.Z. conducted experiments. M.W., M.H.C., M.Z., J.G., C.M.B., M.S., U.K., S.P., E.A., G.T.-L., N. B.-C., P.K., and J.B.Z. analyzed data. M.W., M.H.C., M.Z., J.G., C.M.B., M.S., U.K., S.P., G.T.-L., N. B.-C., J.J.P., G.C.-B., and J.B.Z. contributed to data visualization. I.H., A.W., L.S., G.C.-B., and J.B.Z. acquired funding for the project. M.W., M.H.C., and J.B.Z. wrote and edited the manuscript; all authors reviewed the manuscript. G.J., I.H., A.W., L.S., G.C.-B., and J.B.Z. supervised and mentored contributors. J.B.Z. oversaw all aspects of the work.

## Acknowledgments

We thank the members of the Zuchero laboratory for feedback and support; David Ornitz and Juha Partanen for FGFR1-flox and FGFR2-flox mice; N. Wall, A. Ponnuswami, J. Shepherd, S. Dhulipala at the Stanford Gene Vector and Virus Core (RRID:SCR_023250); Jing-Ping Lin, Daniel Reich, Jack Antel, Niamh McNamara, Inge Huitinga, Stefanie Giera, Karen Chandross, Wayne Rowe, Luke Hanlin, and Ethan Hughes for helpful discussions; and W. Zaw and B. Figueroa at Zeiss for lattice lightsheet imaging. Figure schematics were created with BioRender.com and Adobe Illustrator. We gratefully acknowledge funding from the Stanford University graduate training in Stem Cell Biology and Regenerative Medicine training grant T32-GM119995 (M.W.), Stanford Bio-X Lavidge and McKinley Interdisciplinary fellowship (M.H.C.), Stanford University Medical Scientist Training Program grant T32-GM007365 (M.H.C.), MSCA-postdoctoral fellowship (MonitorMS, 101151194) (M.Z.), and the Schmidt Science Fellows (M.S.). Electron microscopy studies were supported, in part, by ARRA Award Number 1S10RR026780-01 from the National Center for Research Resources (NCRR); their contents are solely the responsibility of the authors and do not necessarily represent the official views of the NCRR or the National Institutes of Health; the Cell Sciences imaging Facility (RRID:SCR_017787); and OneView 4K sCMOS (NIH S10 1S10OD028536-01). A.W. and N.B.C. were funded by the MS Society UK and the Medical research Council (UK). Work in G.C.-B.’s research group was supported by the Swedish Research Council (Distinguished Professor grant 2023-00324), the European Union (Horizon 2020 Research and Innovation Programme/European Research Council Advanced Grant SingleMS, grant agreement number 101096064), Knut and Alice Wallenberg Foundation (Wallenberg Scholar grant 2023-0280) and Strategic Research Area Stem Cells and Regenerative Medicine (Karolinska Institutet). M.Z. was supported by a Marie Skłodowska-Curie Actions post-doctoral fellowship. Work in J.B.Z.’s research group was supported by the National Multiple Sclerosis Society Research Grant RG-2307-42032 and Harry Weaver Scholar Award, Arnold and Mable Beckman Foundation Beckman Young Investigator Award, Stanford Knight Initiative for Brain Resilience Catalyst Award, NIH grant R01NS119823, NIH grant R21NS131999, Sanofi iAward, the McKnight Endowment Fund for Neuroscience, and the Myra Reinhart Family Foundation.

## Competing interests

J.B.Z. has received research funding from Sanofi, related to the current study. J.B.Z.’s spouse is employed by and holds equity in Denali Therapeutics, which is not involved in this work. G.C.-B is shareholder of Nexus Epigenomics. All other authors declare no competing interests.

## Materials and Correspondence

The data that support the findings of this study and step-by-step protocols are available from the lead author or first authors upon request. Bulk and single-cell RNA-seq data will be deposited at NCBI GEO prior to publication. Proteomics datasets will be deposited at PRIDE prior to publication. All AAV constructs developed for this study will be deposited at Addgene and made available at the time of publication (AAV_pGFAabc1d-FGF1, AAV_pGFAabc1d-FGF2). Correspondence and requests for materials should be addressed to M.H.C. and J.B.Z.

## Supplementary Information

- Extended Data Fig. 1–15
- Supplementary Data Tables 1–6
- Supplementary Videos 1–5

---

**Extended Data Fig. 1.**
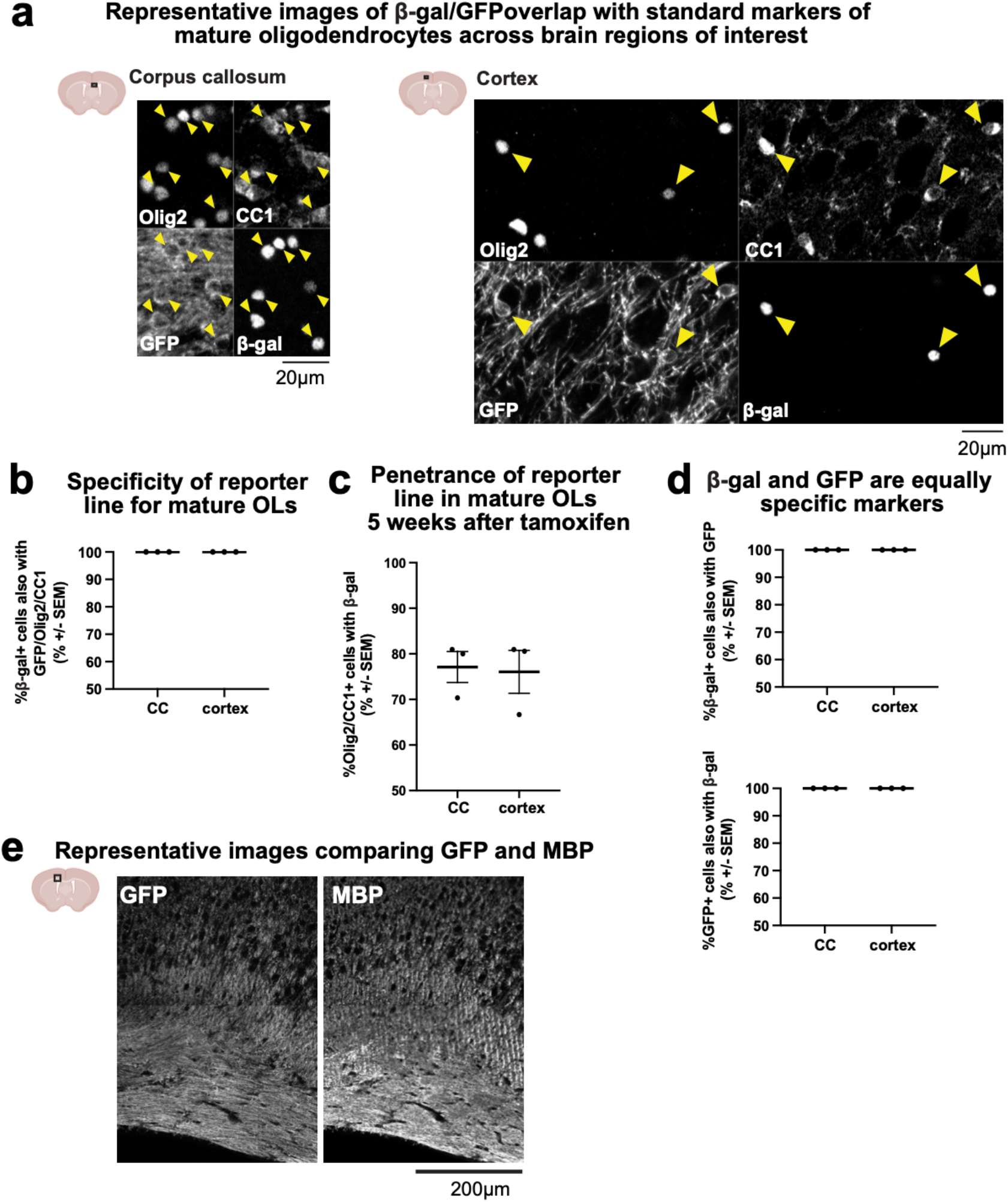
Characterization of lineage tracing mice. **a**, Representative image in cortex (left) and corpus callosum (right) of GFP+ cell soma and β-gal nuclei with double positive CC1 (mature oligodendrocytes) and Olig2 (OL lineage), confirming specificity of GFP/β-gal lineage tracing. Yellow arrow heads indicate GFP/β-gal/CC1/Olig2 positive mature OLs. **b**, Quantification of specificity of reporter line. Corpus callosum (CC): 100 ± 0.0%; Cortex: 100 ± 0.0%. **c**, Quantification of penetrance of reporter line. CC: 77.1 ± 3.39%; Cortex: 76.1 ± 4.70%. **d**, Quantification validating GFP and β-gal as interchangeable markers for mature oligodendrocyte specificity. CC: 100 ± 0.0%; Cortex: 100 ± 0.0%. *n = 3* biological replicates. **e**, Representative image of GFP and MBP, confirming GFP labeling of sheaths is similar to MBP.

**Extended Data Fig. 2.**
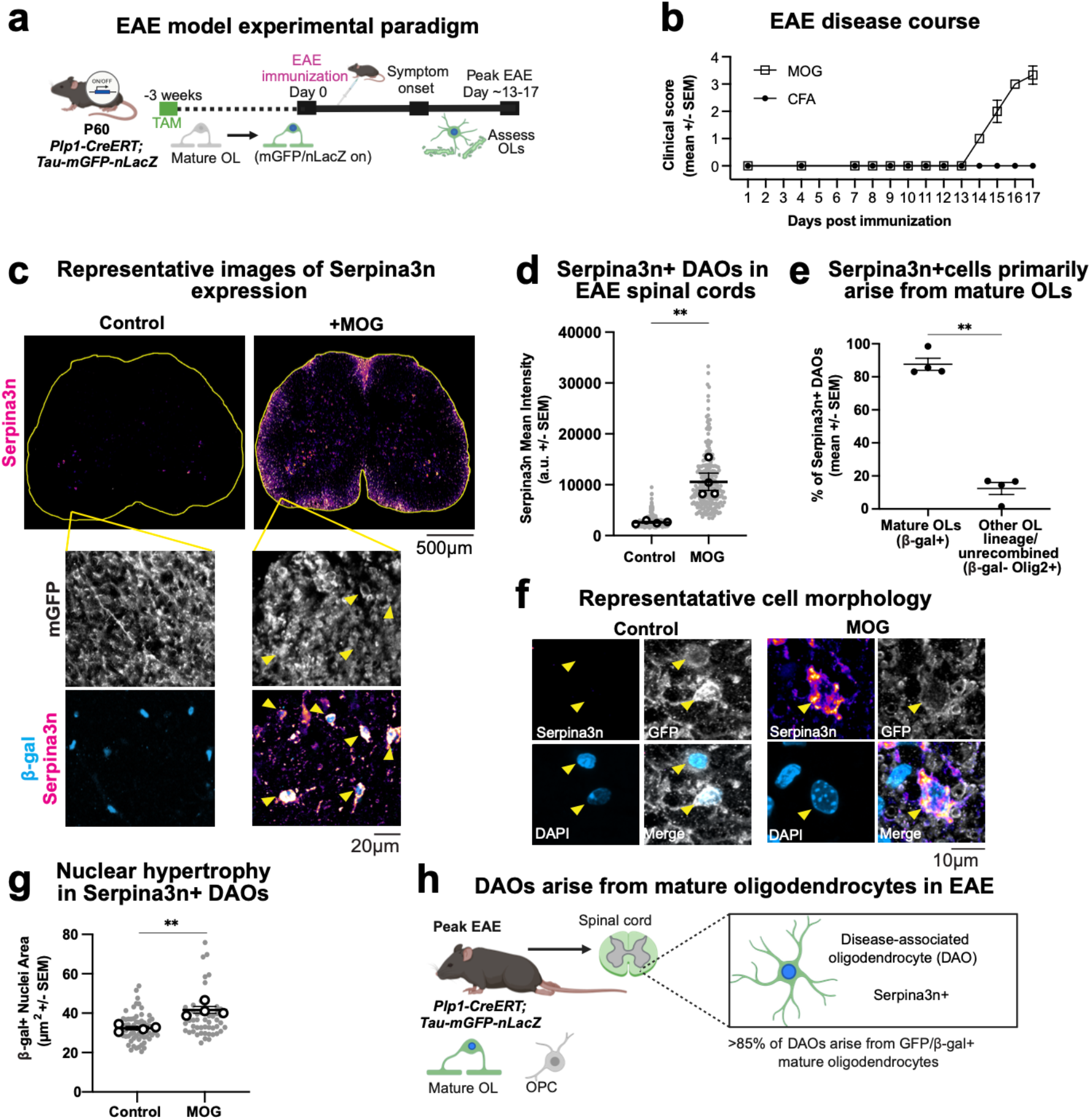
Mature oligodendrocytes become disease-associated oligodendrocytes (DAOs) **a**, Cartoon schematic of EAE experimental paradigm in reporter mice. Mice were immunized 3 weeks following tamoxifen and assessed one day past peak EAE clinical score. **b**, EAE disease course represented by clinical score in control (Complete Freund’s Adjuvant, CFA) and EAE (MOG) mice. Mean clinical score ± SEM, *n = 4* replicate mice per condition for all experiments shown in Extended Data Fig. 2. **c**, Representative stitched-tiled images of Serpina3n protein expression by immunostaining in control and MOG spinal cords. Yellow arrowheads in insets of higher magnification images highlight Serpina3n/β-gal/GFP triple positive DAOs. **d**, Quantification of Serpina3n intensity (images shown in **c**) shows increased Serpina3n expression in β-gal+ cells in MOG spinal cords compared to controls. Black dots represent biological replicates, superplot of grey dots are individual cell measurements. Serpina3n mean intensity ± SEM; Control: 2603 ± 157.1 a.u., MOG: 10571 ± 1704 a.u. **e**, Quantification of Serpina3n+ mature (β-gal+) vs other Serpina3n+ oligodendrocyte lineage cells (Olig2+ β-gal-) shows the majority of Serpina3n+ oligodendrocyte lineage cells derive from mature oligodendrocytes. Mean % of Serpina3n+ OL lineage cells ± SEM, DAOs from lineage traced mature OLs: 87.6 ± 3.67%, DAOs from other OL lineage cells (or mature OLs that did not recombine): 12.4 ± 3.67%. **f**, Representative images of normal nuclei and soma in control (left) and enlarged nuclei with hypertrophied, Serpina3n+ soma in MOG (right) spinal cords in sparse-labelled lineage tracing mice (63X maximum intensity projection). **g**, Quantification of β-gal+ cell nuclear area shows increased nuclear size in MOG spinal cords compared to controls. β-gal+ cell nuclear area ± SEM; Control: 32.45 ± 0.84 µm^2^, MOG: 41.65 ± 1.74 µm^2^. **h**, Model: Using the highly specific *Plp1-CreERT;Tau-mGFP-nLacZ* reporter mice we show DAOs primarily arise from mature oligodendrocytes. Statistical measurement (*p* value) determined by unpaired, two-tailed t-test, *p* values: \*\**p* < 0.01.

**Extended Data Fig. 3.**
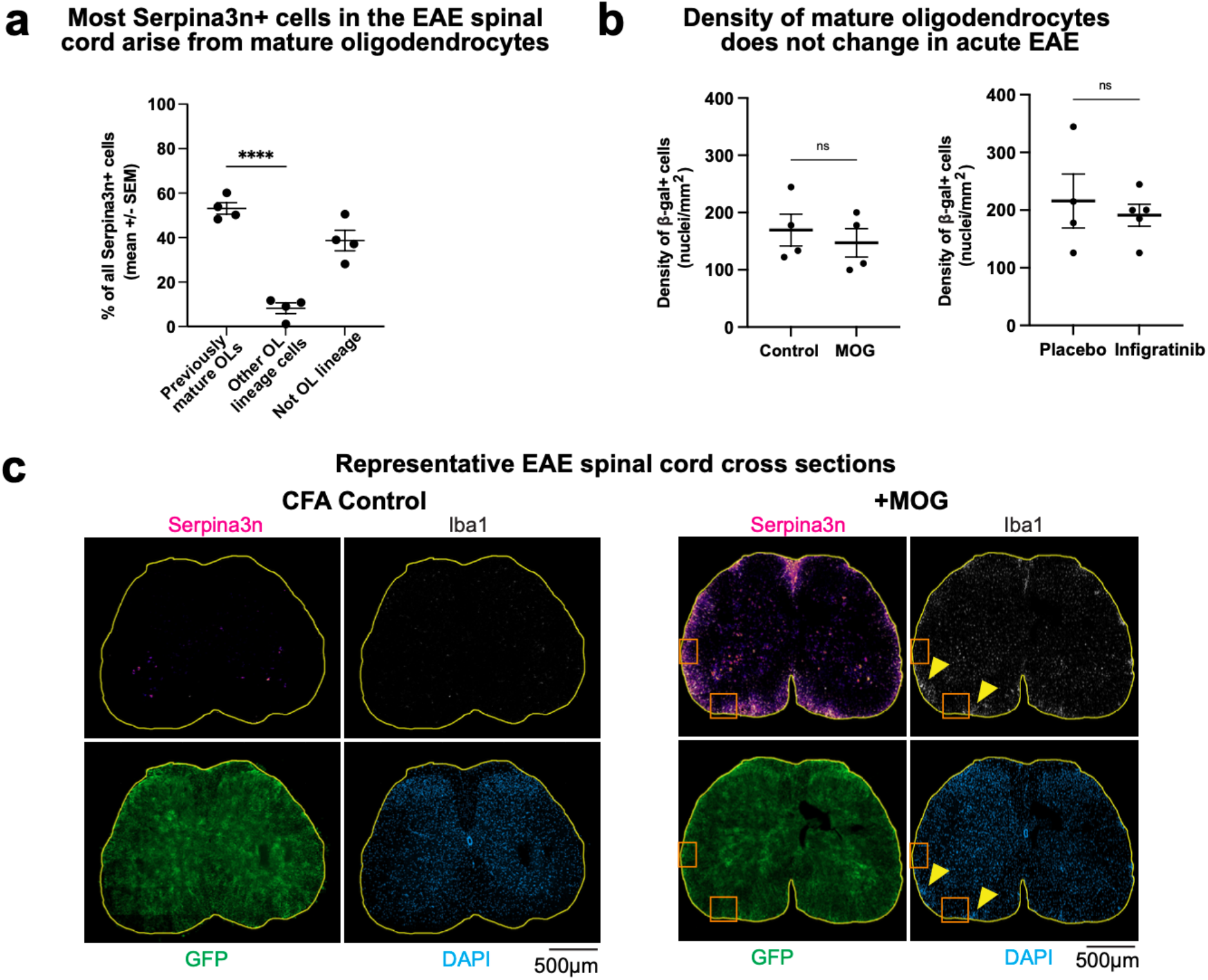
Additional characterization of EAE model. **a**, Quantification of all Serpina3n+ cells in EAE spinal cords shows the majority of all Serpina3n+ cells derive from mature oligodendrocytes. Mean % of all Serpina3n+ cells ± SEM, Serpina3n+ cells from mature OLs: 53.23 ± 2.60%, from other OL lineage cells: 8.18 ± 2.42%, from all other cells (not OL lineage): 38.69 ± 4.61%, *n = 4* replicates. **b**, Quantification of β-gal nuclei density in EAE spinal cords showing density does not change in lesion-adjacent regions imaged. Comparing CFA control vs. MOG (Extended Data Fig. 2): CFA 169.4 ± 27.73 nuclei/µm^2^, MOG: 147.2 ± 24.59 nuclei/µm^2^. Comparing MOG +placebo control vs. MOG + infigratinib (pan FGFR-inhibitor) in Infigratinib EAE experiment (Extended Data Fig. 14): Placebo 215.7 ± 46.61 nuclei/µm^2^, MOG: 191.1 ± 19.01 nuclei/µm^2^, *n = 4* replicates. **c**, Representative images of DAPI, GFP, Serpina3n, and Iba1 (microglia) protein expression by immunostaining in CFA (left) and MOG (right) spinal cords. Yellow arrowheads indicate regions considered lesions (characterized by microglia infiltration and/or high DAPI density). Orange boxes indicate example regions adjacent to lesions that were imaged and analyzed in this study. If no obvious lesions were present (e.g. in CFA example on bottom), ROIs in a similar area were chosen and imager remained blind to condition. Statistical measurements (*p* value) were determined by an unpaired, two-tailed t-test, *p* values: \*\*\*\**p* < 0.0001; *ns* = not significant.

**Extended Data Fig. 4.**
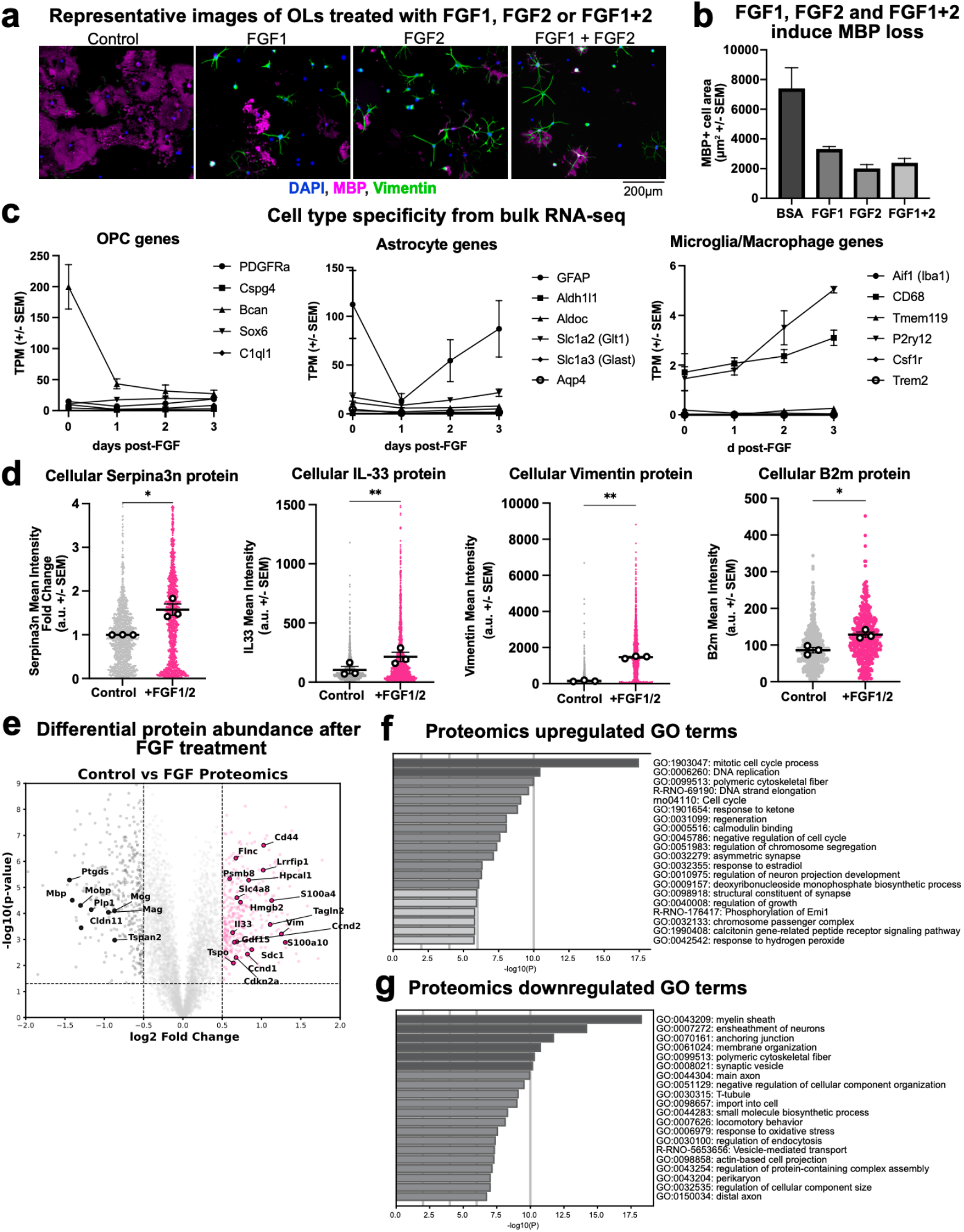
Additional characterization of FGF1/2-induced changes in culture. **a**, Representative images of cultured oligodendrocytes treated with BSA (control), FGF1, FGF2, or both FGF1 and 2 (FGF1/2) for 72hrs. Stained for DAPI (blue), MBP (magenta), and vimentin (green) illustrating that FGF1, FGF2 and the combination of FGF1 and FGF2 all similarly induce oligodendrocytes into a DAO-like state in culture with increased vimentin expression and morphological changes. **b**, Quantification of MBP+ cell area in oligodendrocytes treated with BSA (control), FGF1, FGF2, or FGF1/2; MBP area per cell (µm^2^) ± SEM; BSA (control): 7,361 ± 1,446; FGF1: 3,276 ± 207.9; FGF2: 1,974 ± 301; FGF1+2: 2,355 ± 334, *n = 1* biological replicate. **c**, Gene expression changes from bulk RNA-seq of FGF1/2 treated cultured oligodendrocytes over 72hrs showing cells are not upregulating OPC genes (left), nor more astrocyte (middle) or microglial (right) genes. Cells upregulate isolated genes typical of astrocytes (GFAP) and microglia (CD68 and P2ry12), but are not becoming another known cell type. Transcripts per million (TPM) ± SEM on y-axis, *n = 3* biological replicates. **d**, Superplots of mean intensity of DAO-gene protein expression in cultured oligodendrocytes 72hrs after FGF1/2 treatment. Large black dots represent biological replicates, superplot of small colored dots are individual cell measurements, *n = 3* biological replicates. Also see Figure 1e-h. **e**, Volcano plot visualization of unbiased proteomics of control (20ng/ml 0.2% BSA + heparin, 72 hours) and FGF1/2 treated (10ng/ml FGF1 + 10ng/ml FGF2 + heparin, 72 hours) oligodendrocytes showing differential protein abundance; *n = 4* biological replicates. **f**, GO term enrichment analysis of proteomics data of cultured oligodendrocytes treated with FGF1/2 for 72hrs compared to control oligodendrocytes (upregulated terms). **g**, GO term enrichment analysis of proteomics data of cultured oligodendrocytes treated with FGF1/2 for 72hrs compared to control oligodendrocytes (downregulated terms). Statistical measurements (*p* value) were determined by an unpaired, one-tailed t-test, *p* values: \**p* < 0.05; \*\**p* < 0.01.

**Extended Data Fig. 5.**
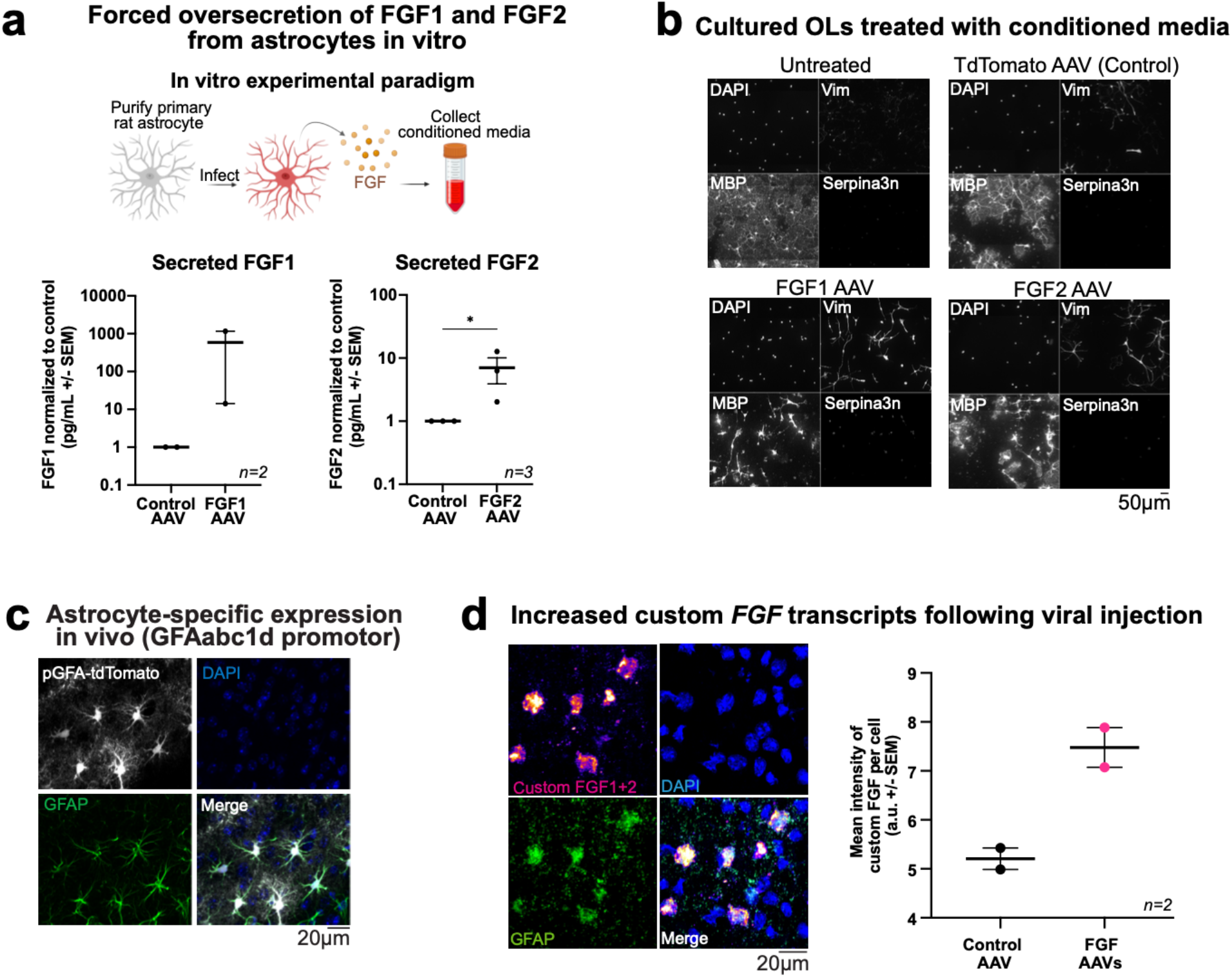
Characterization of FGF overexpression AAVs. **a**, Schematic of in vitro astrocyte infection experiment to validate function of FGF1/2 AAVs, and quantification of FGF1 (left) or FGF2 (right) in astrocytes infected with tdTomato AAV as control vs. custom FGF1 AAV or FGF2 AAV. For FGF1 secretion - Control: 1.0 ± 0.0 pg/mL; FGF1 AAV: 586.5 ± 572.4 pg/mL, *n* = 2 biological replicates. For FGF2 secretion - Control: 1.0 ± 0.0pg/mL; FGF2 AAV: 7.0 ± 3.09 pg/mL, *n* = 3 biological replicates. **b**, Representative images of cultured rat oligodendrocytes treated with conditioned media from astrocytes infected with tdTomato AAV as control vs. custom FGF1 AAV or FGF2 AAV. Loss of MBP, increase in Vimentin, increase in Serpina3n, and change in morphology confirm functionality of FGF1 and FGF2 produced with AAV overexpression paradigm. **c**, Representative image of pGFAabc1d-tdTomato (AddGene #44332) expression in astrocytes stained with GFAP in mouse brain. **d**, Representative RNAscope image and quantification of custom *FGF1* and *FGF2* transcripts in mouse brain 14 days after infection with custom FGF1/2 AAVs; Control: 5.207 ± 0.22 a.u.; FGF: 7.479 ± 0.41 a.u., *n = 2* biological replicates. Expression of custom FGF1/2 overlaps with GFAP transcript expression, confirming expression in astrocytes. Statistical measurements (*p* value) where appropriate were determined by a ratio t-test, *p* values: \**p* < 0.05.

**Extended Data Fig. 6.**
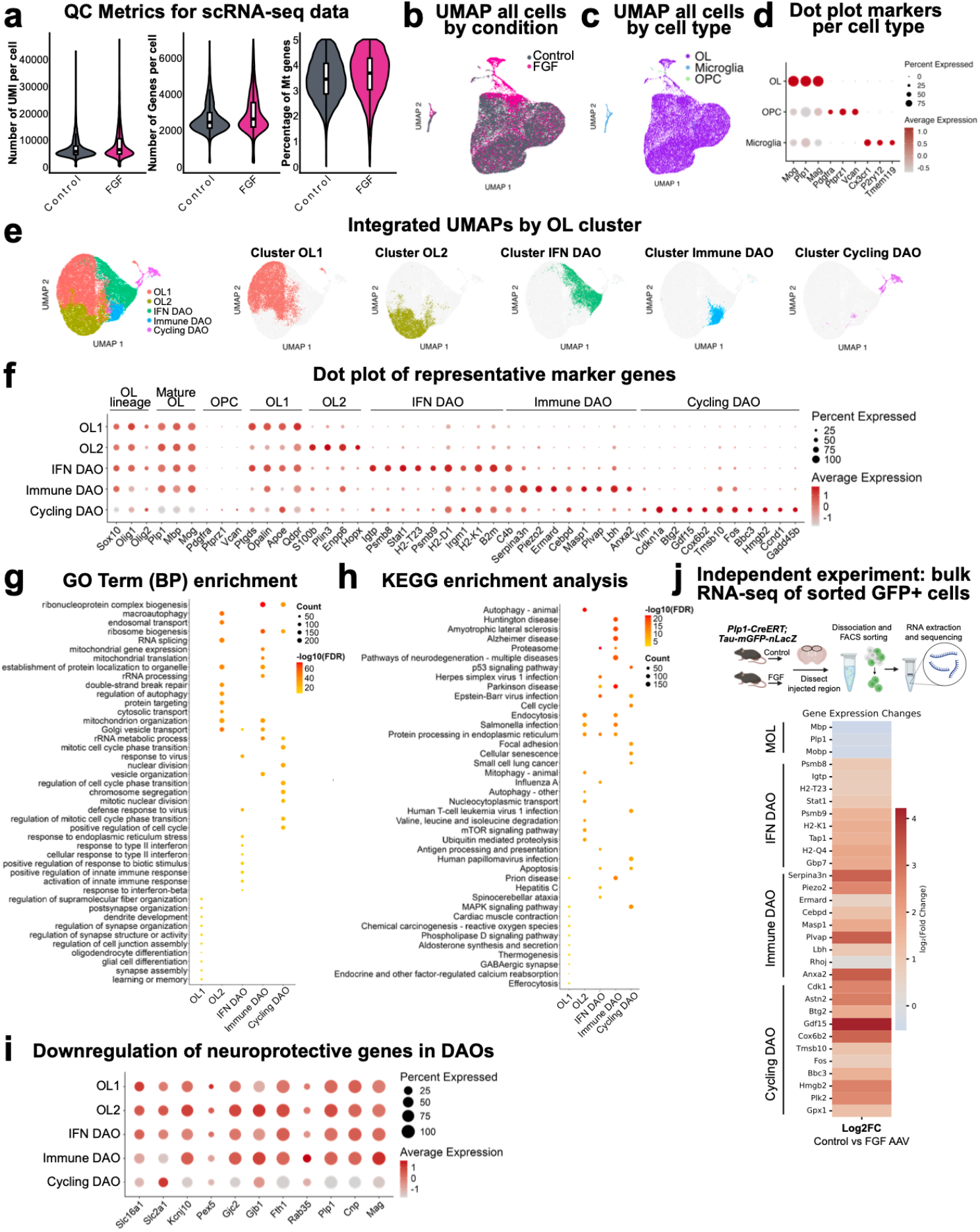
scRNA-seq reveals disease-associated oligodendrocyte clusters. **a**, Violin plots showing QC metrics (UMI counts per cell, number of genes per cell, and percentage of mitochondrial genes) for tdTomato control (grey) and FGF (pink) samples. **b**, Uniform manifold approximation and projection (UMAP) plot showing all cells integrated between control (grey) and FGF (pink) samples. **c**, Integrated UMAP of all cells colored by cell type, confirming majority mature oligodendrocytes as expected with FACS sorting of GFP+ cells: 27669 cells were mature oligodendrocytes, 102 were OPCs, and 657 cells were microglial contaminants. OPC and microglial contaminants could represent cells that phagocytosed myelin-associated GFP. **d**, Dot plot of canonical gene expression for cell types in integrated dataset illustrating clear mature oligodendrocyte population and contaminating OPCs and microglia to filter out for all further analysis. Color scale represents the average expression of a given gene, and the size of the dot represents the percentage of cells that express that gene. **e**, Integrated UMAP of all mature oligodendrocytes colored by cluster. Number of cells in each cluster: OL1: 13826 cells; OL2: 6179 cells; IFN DAO: 5267 cells; immune DAO: 1403 cells; cycling DAO: 994 cells. **f**, Dot plot of representative marker genes for each oligodendrocyte lineage cluster in integrated dataset. Color scale represents the average expression of a given gene, and the size of the dot represents the percentage of cells that express that gene. **g**, GO term enrichment analysis comparing differentially expressed genes between all mature oligodendrocyte clusters using GO Biological Process (BP) terms. **h**, KEGG enrichment analysis comparing differentially expressed genes between all mature oligodendrocyte clusters using GO Biological Process terms. **i**, Dot plot of genes associated with neuroprotective function for each oligodendrocyte lineage cluster in integrated dataset. Color scale represents the average expression of a given gene, and the size of the dot represents the percentage of cells that express that gene. **j**, Heatmap of gene expression from separate bulk RNA-seq experiment comparing pooled reporter mouse brains infected with control AAV vs. FGF1 and FGF2 AAVs for 14 days. Top genes expressed in each oligodendrocyte cluster from Figure 2i shown, confirming similar gene expression changes in independent experiment. Colors indicate log2FC of gene expression in FGF sample compared to tdTomato control.

**Extended Data Fig. 7.**
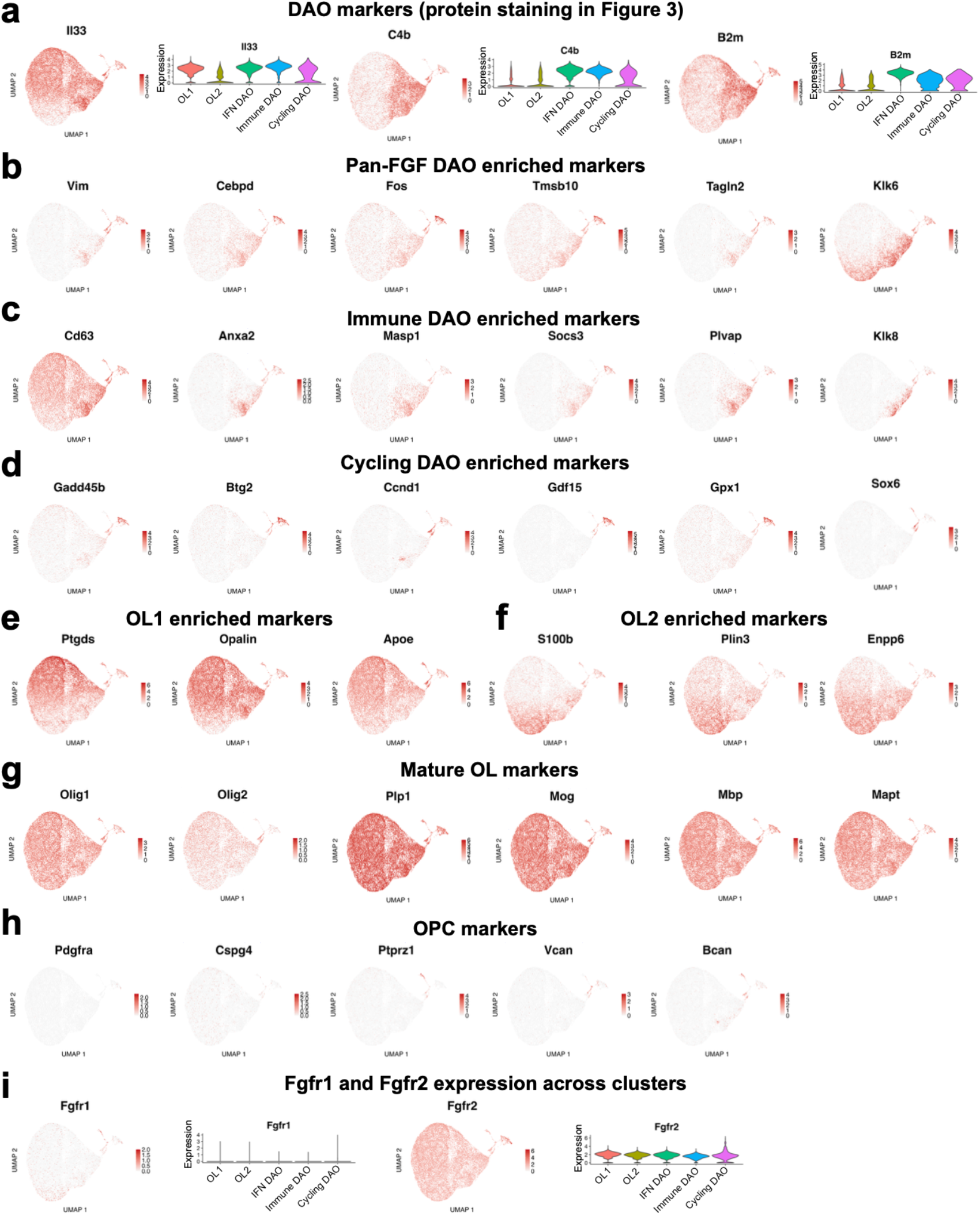
scRNA-seq cluster-specific markers. **a**, UMAP visualizations with expression of key DAO genes shared across DAO subtypes (*Il33, C4b, B2m*) shown in red. Violin plots depict expression of respective gene in each cluster across all mature OL cell clusters. **b**, UMAP visualizations with expression of genes enriched across both FGF-responsive DAO populations, Immune DAOs and Cycling DAOs (*Vim, Cebpd, Fos, Tmsb10, Tagln2, Klk6*). **c**, UMAP visualizations with expression of genes enriched in Immune DAOs (*Cd63, Anxa2, Masp1, Socs3, Plvap, Klk8*). **d**, UMAP visualizations with expression of genes enriched in Cycling DAOs (*Gadd45b, Btg2, Ccnd1, Gdf15, Gpx1, Sox6*). **e**, UMAP visualizations with expression of genes enriched in OL1 population of homeostatic OLs (*Ptgds, Opalin, Apoe*). **f**, UMAP visualizations with expression of genes enriched in OL2 population of homeostatic OLs (*S100b, Plin3, Enpp6*). **g**, UMAP visualizations with expression of mature oligodendrocyte genes illustrating expression across all clusters (*Olig1, Olig2, Plp1, Mog, Mbp, Mapt*). **h**, UMAP visualizations with expression of canonical OPC genes illustrating lack of expression across all clusters (*Pdgfra, Cspg4, Ptprz1, Vcan, Bcan*). **i**, UMAP visualizations with expression of *Fgfr1* and *Fgfr2*, the FGFRs expressed by mature oligodendrocytes illustrating expression of both receptors across all clusters. Violin plots depict expression of respective gene in each cluster across all mature OL cell clusters.

**Extended Data Fig. 8.**
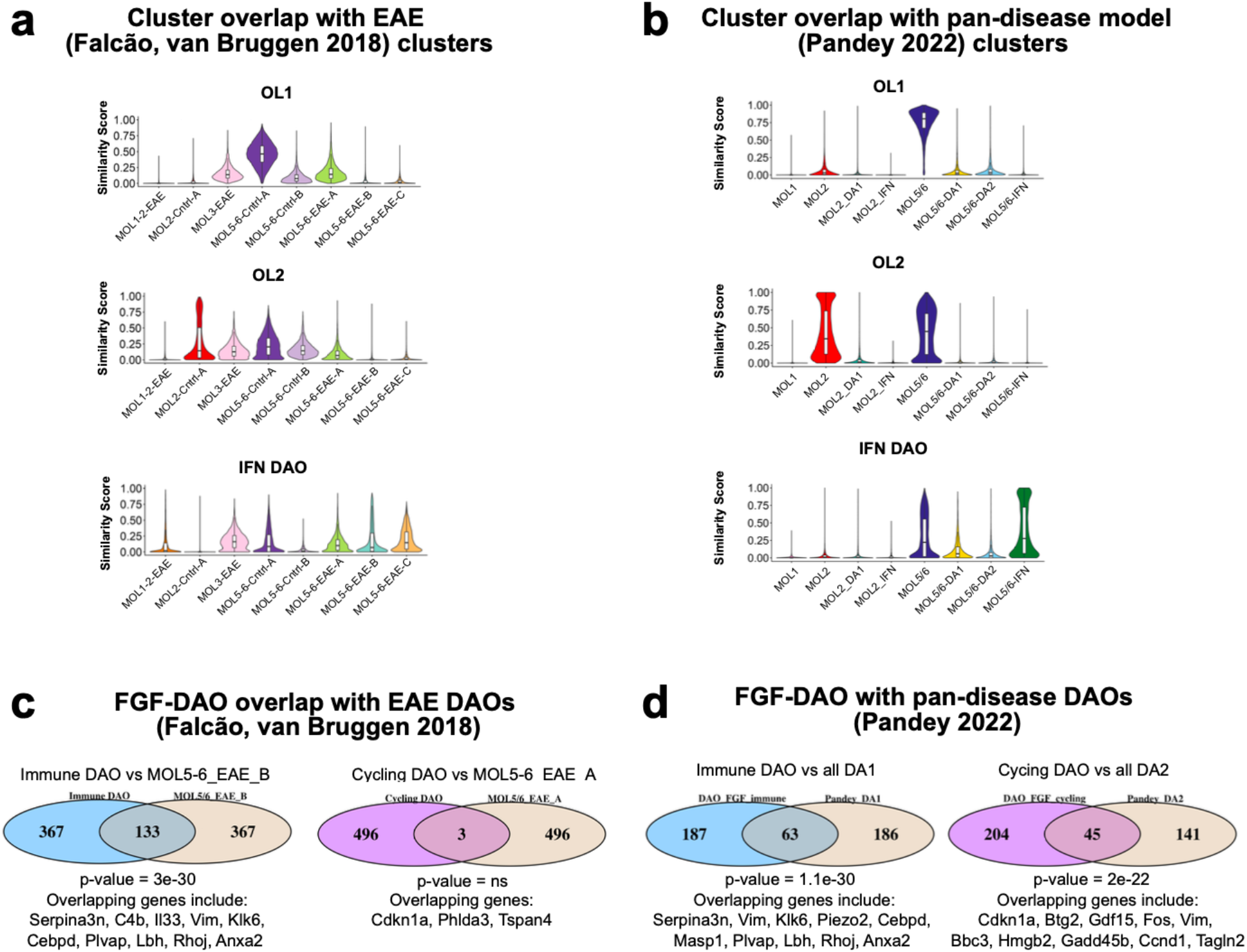
FGF-induced DAOs are similar to DAOs across disease models. **a**, Similarity scores between mature oligodendrocyte clusters not specific to FGF samples (OL1, OL2, and IFN DAO) compared to all cell clusters from Falcão, Van Bruggen 2018 EAE dataset. **b**, Similarity scores between other mature oligodendrocyte clusters not specific to FGF samples (OL1, OL2, and IFN DAO) compared to all cell clusters from Pandey 2022 pan-disease dataset. **c**, Overlap between top 500 upregulated genes in FGF-induced DAOs compared to top 500 upregulated genes in most similar population from EAE: Immune DAO vs. MOL5/6_EAE_B (left), and Cycling DAOs vs. MOL5_6_EAE_A (right) from Falcão, Van Bruggen 2018 EAE dataset. Fisher’s exact test was used to calculate the overlapping p value for each comparison. Relevant overlapping genes listed below diagram. **d**, Overlap between top 250 upregulated genes in Immune DAOs compared to top upregulated genes in DA1 (left), or comparing Cycling DAOs to top upregulated genes in DA2 (right) from Pandey 2022 pan-disease dataset. Fisher’s exact test was used to calculate the overlapping p value for each comparison.

**Extended Data Fig. 9.**
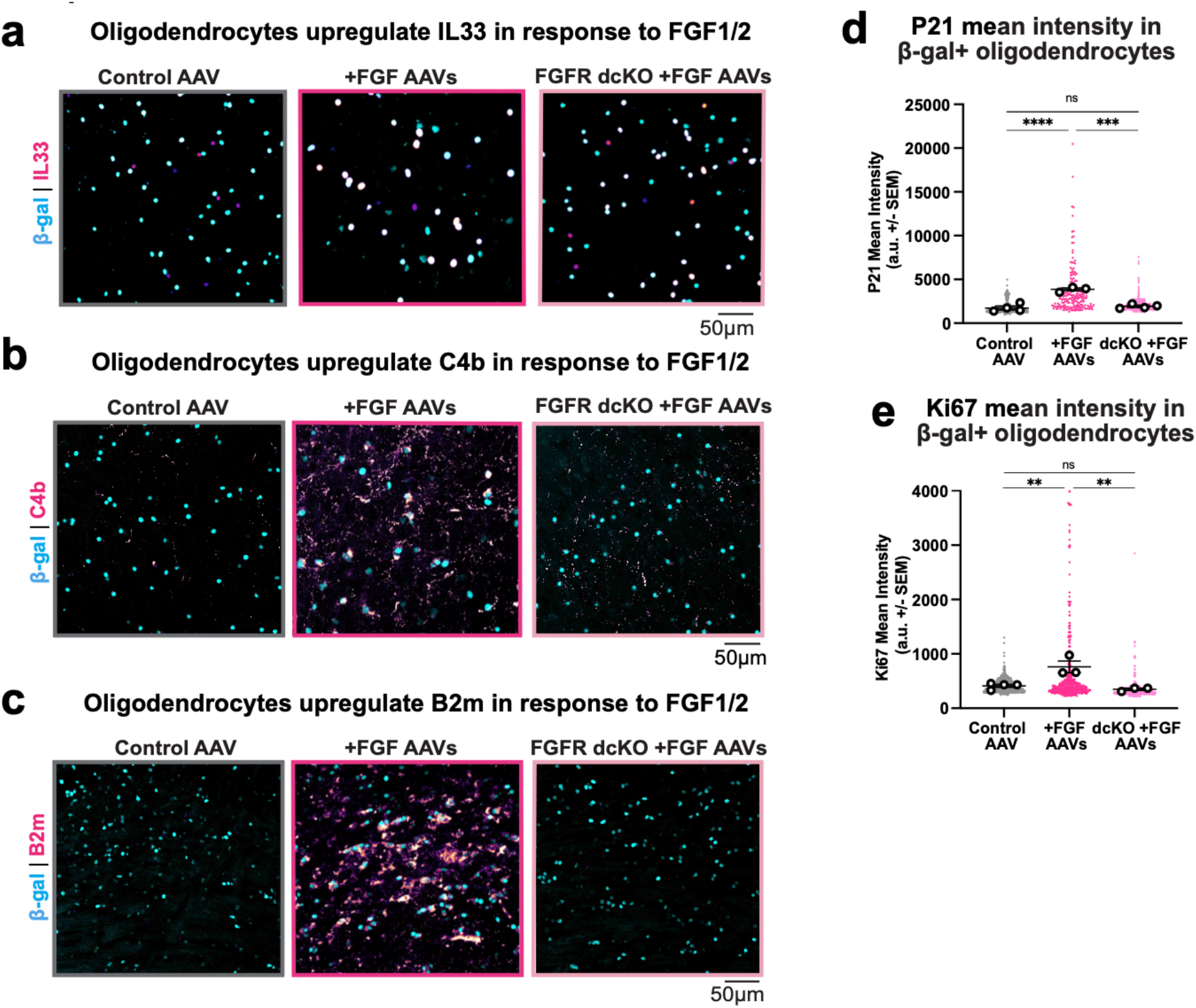
Validation of protein expression of DAO markers in response to FGF-AAVs. **a**, Representative immunostaining of IL-33 and β-gal in reporter mice or dcKO reporter mice injected with control AAVs or FGF1/2 AAVs. **b**, Representative immunostaining of C4b and β-gal in reporter mice or dcKO reporter mice injected with control AAVs or FGF1/2 AAVs. **c**, Representative immunostaining of B2m and β-gal in reporter mice or dcKO reporter mice injected with control AAVs or FGF1/2 AAVs. **d**, Quantification of P21 mean intensity in β-gal+ cells, normalized to control ± SEM; Control: 1730 ± 215.4 a.u.; FGF: 3862 ± 164.4 a.u.; dcKO+FGF: 1930 ± 117.3 a.u. **e**, Quantification of Ki67 mean intensity in β-gal+ cells, normalized to control ± SEM; Control: 411.5 ± 28.08 a.u.; FGF: 762.1 ± 106.3 a.u.; dcKO+FGF: 349.4 ± 20.22 a.u., *n = 3-4* biological replicates. Statistical measurements (*p* value) were determined by 1 way ANOVA with Tukey’s multiple comparison test, *p* values: \*\**p* < 0.01; \*\*\**p* < 0.001; \*\*\*\**p* < 0.0001; *ns* = not significant.

**Extended Data Fig. 10.**
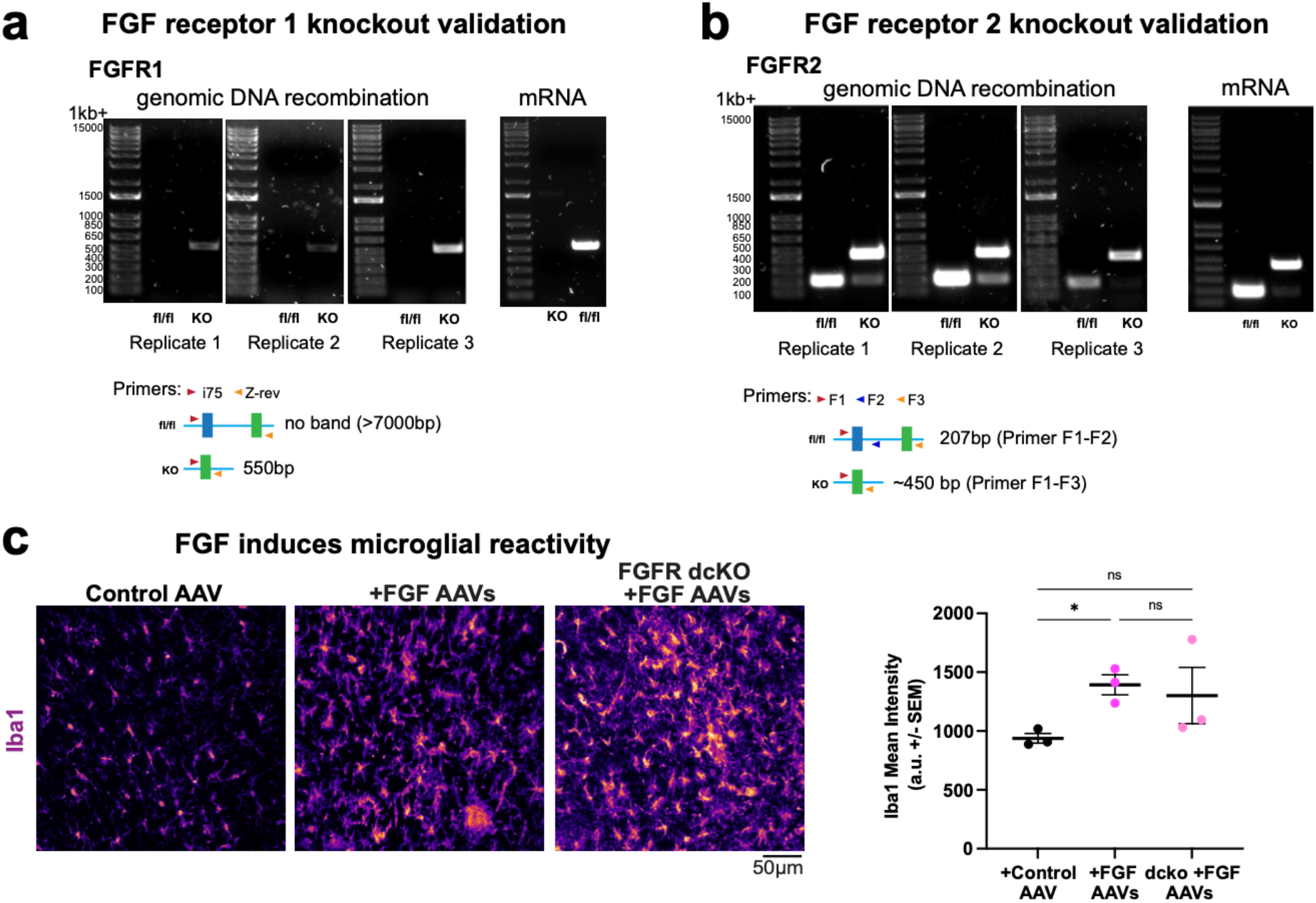
Knockout validation in *Plp1-CreERT;FGFR1fl/fl;FGFR2fl/fl* mice. **a**, Reduction of FGFR1 genomic DNA in OLs purified 3 weeks after tamoxifen recombination of FGFR1fl/fl;FGFR2fl/fl (fl/fl) and *Plp1-CreERT;FGFR1fl/fl;FGFR2fl/fl* (KO) mice. Lack of amplified DNA in fl/fl confirms intact DNA. Band ~550bp in KO confirms recombination. **b**, Same paradigm with FGFR2 genomic DNA. DNA band ~207bp in fl/fl confirms intact DNA. Band ~450bp in KO confirms recombination. Note persistent 207bp band in 2 of 3 KO replicates, indicating recombination efficiency is not 100%. DNA collected from mature oligodendrocytes from three separate mice per condition. mRNA (converted to cDNA) collected from mature oligodendrocytes from one mouse per condition. **c**, Representative immunostaining and quantification of Iba1 (microglia) in reporter mice or dcKO reporter mice injected with control AAVs or FGF1/2 AAVs, showing microglia reactivity in response to FGF is not affected by conditional FGFR knockout, Iba1 mean intensity (a.u.) ± SEM; Control: 937.9 ± 40.92; FGF: 1393 ± 85.13; dcko+FGF: 1301 ± 238.9, *n = 3* biological replicates. Statistical measurements (*p* value) were determined by 1 way ANOVA with Tukey’s multiple comparison test, *p* values: \**p* < 0.05; *ns* = not significant.

**Extended Data Fig. 11.**
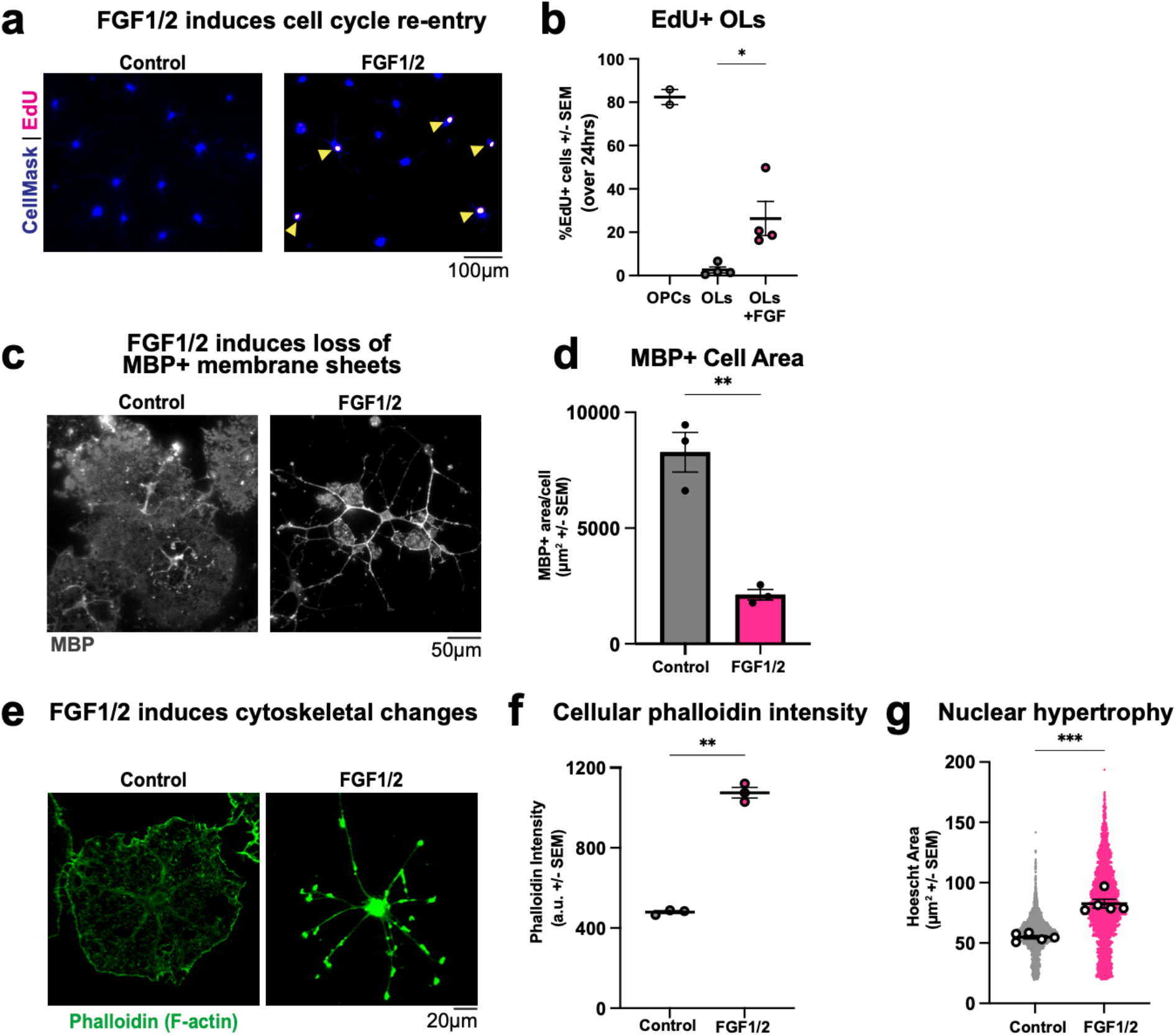
FGF alters oligodendrocyte homoeostasis in culture. **a**, Representative images of 24hrs of EdU incorporation in immunopanned primary rat oligodendrocytes treated with FGF1/2 (right) or a vehicle control (left). **b**, Quantification of EdU in a, Percentage of positive cells ± SEM, OPCs: 82.38 ± 3.5%; Control OLs: 2.59 ± 1.38%; OLs +FGF1/2: 26.31 ± 7.86%, *n = 2-4* biological replicates. **c**, Representative images of MBP staining in cultured oligodendrocytes treated with FGF1/2 (right) or a vehicle control (left). **d**, Quantification of MBP membrane loss shown in c. MBP+ area/cell (µm^2^) ± SEM; Control: 8,280 ± 856; FGF1/2: 2,121 ± 226.9, *n = 3* biological replicates. **e**, Representative images of cultured oligodendrocytes treated with FGF1/2 (right) or a vehicle control (left) stained with phalloidin to visualize actin filaments. **f**, Quantification of phalloidin staining in e. Phalloidin mean intensity (a.u.) ± SEM; Control: 479.6 ± 7.16; FGF1/2: 1075 ± 26.16, *n = 3* biological replicates. **g**, Quantification of oligodendrocyte nuclear area by Hoechst staining in control and FGF1/2-treated oligodendrocytes. Hoechst area (µm^2^) ± SEM; Control: 54.88 ± 1.45; FGF1/2: 82.58 ± 3.68, *n = 5* biological replicates. Statistical measurements (*p* value) were determined by an unpaired, two-tailed t-test, *p* values: \**p* < 0.05; \*\**p* < 0.01; \*\*\**p* < 0.001.

**Extended Data Fig. 12.**
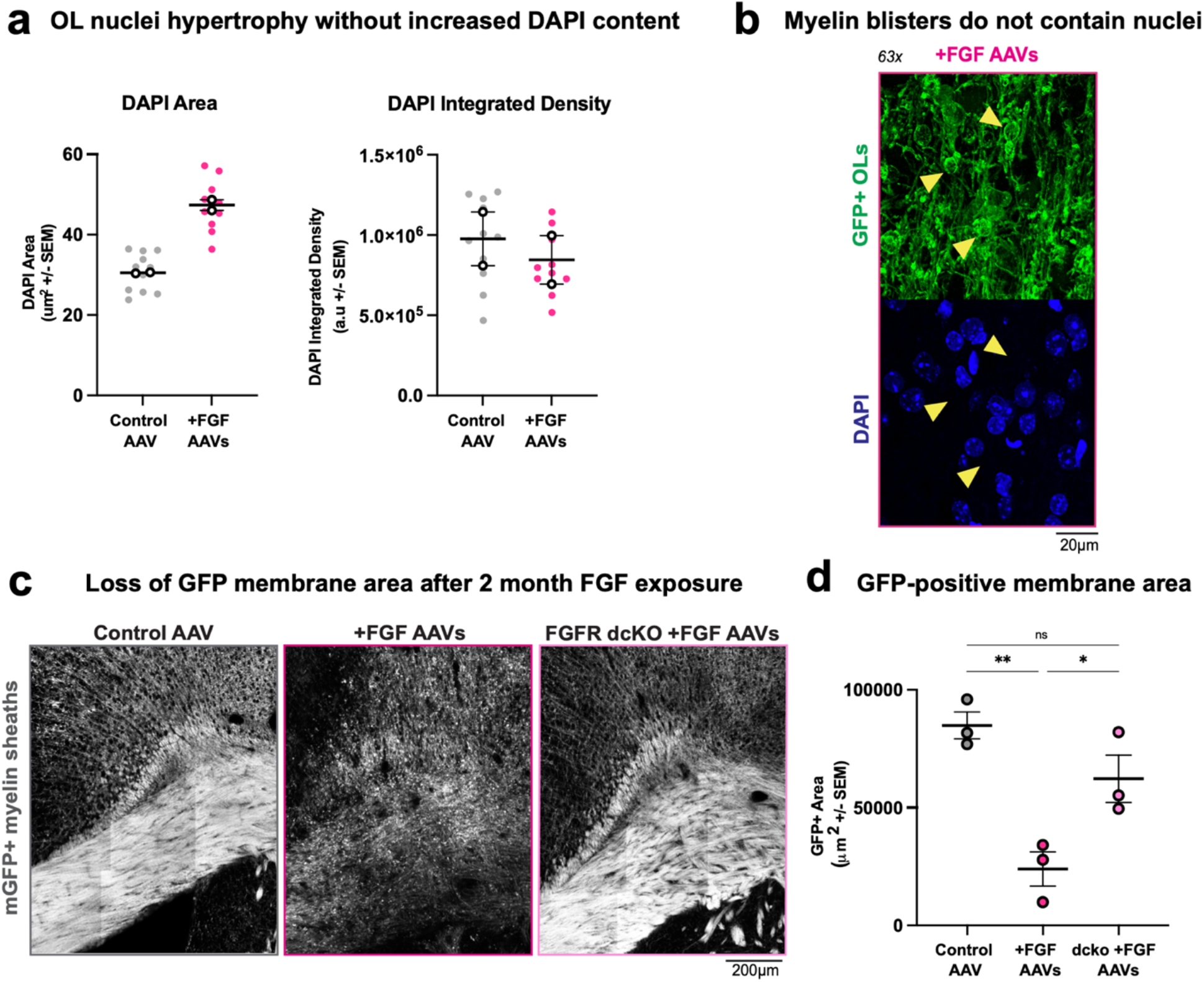
Additional characterization of oligodendrocyte and myelin pathology in response to FGF-AAVs. **a**, Quantification of oligodendrocyte nuclei size with DAPI staining in reporter mouse brains infected with control AAV vs. FGF1 and FGF2 AAVs for 14 days showing increase in nuclei size without increase in total DAPI content (integrated density), DAPI area (µm^2^) ± SEM; Control: 30.53 ± 0.13; FGF: 47.36 ± 1.32, DAPI Integrated Density (a.u.) ± SEM; Control: 976,672 ± 167,746; FGF: 845,352 ± 150,931, *n = 2* biological replicates. **b**, Representative images of myelin sheath morphology in deep cortical layers of lineage tracing mice stained with anti-GFP488 and DAPI to label all nuclei. Dysmorphic myelin “blisters” indicated with yellow arrowheads. DAPI staining shows “blisters” are not dysmyelination of other cell bodies (no underlying DAPI). **c**, Representative images of myelin sheath morphology in the corpus callosum and deep cortical layers shown by mGFP 2 months after injection with control-AAV (control, left), FGF1/2-AAV (middle), or FGF1/2-AAV into FGFR1/2 dcKO mice (right). **d**, Quantification of GFP membrane area in corpus callosum. GFP-positive area (µm^2^) in a given field of view ± SEM; Control: 84,897 ± 5,712; FGF1/2: 23,950 ± 7,269; dcKO+FGF1/2: 62,261 ± 10,014, *n = 3* biological replicates. Statistical measurements (*p* value) were determined by 1 way ANOVA with Tukey’s multiple comparison test, *p* values: \**p* < 0.05; \*\**p* < 0.01; *ns* = not significant.

**Extended Data Fig. 13.**
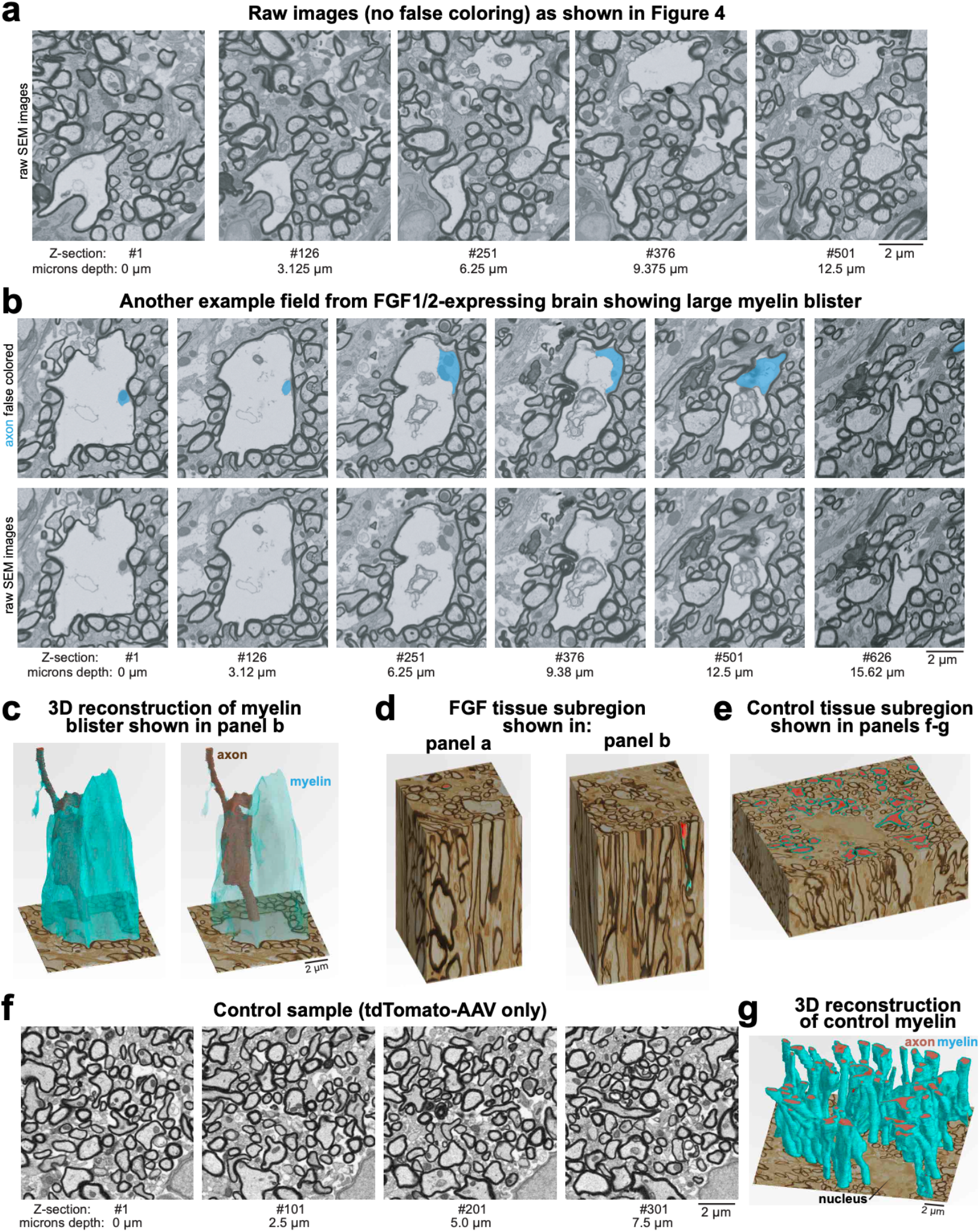
pFIB-SEM of FGF-AAV injected samples. **a**, Raw images (no pseudocoloring) of FIBSEM slices (every 3.125µm) through block of tissue from deep cortical layers in mice 14 days after injection with FGF1/2 AAVs shown in Fig. 4k. **b**, Second example of FIBSEM slices (every 3.125µm) through block of tissue from FGF1/2 sample. Pseudocolor (top) highlighting axon with normal-appearing compact myelin progressing to myelin blister with space between axon and myelin sheath. Raw images (no pseudocoloring) on bottom. **c**, 3D reconstruction from FIBSEM of myelin blister in FGF1/2 sample shown in b. Pseudocolors highlight myelin (teal) and underlying axon (brown). **d**, Reconstruction of 3D tissue block imaged to create panel a. **e**, Reconstruction of 3D tissue block imaged to create panel f-g. **f**, FIBSEM slices (every 3.125µm) through block of tissue from control (tdTomato AAV) sample. **g**, 3D reconstruction from FIBSEM of control myelin in f.

**Extended Data Fig. 14.**
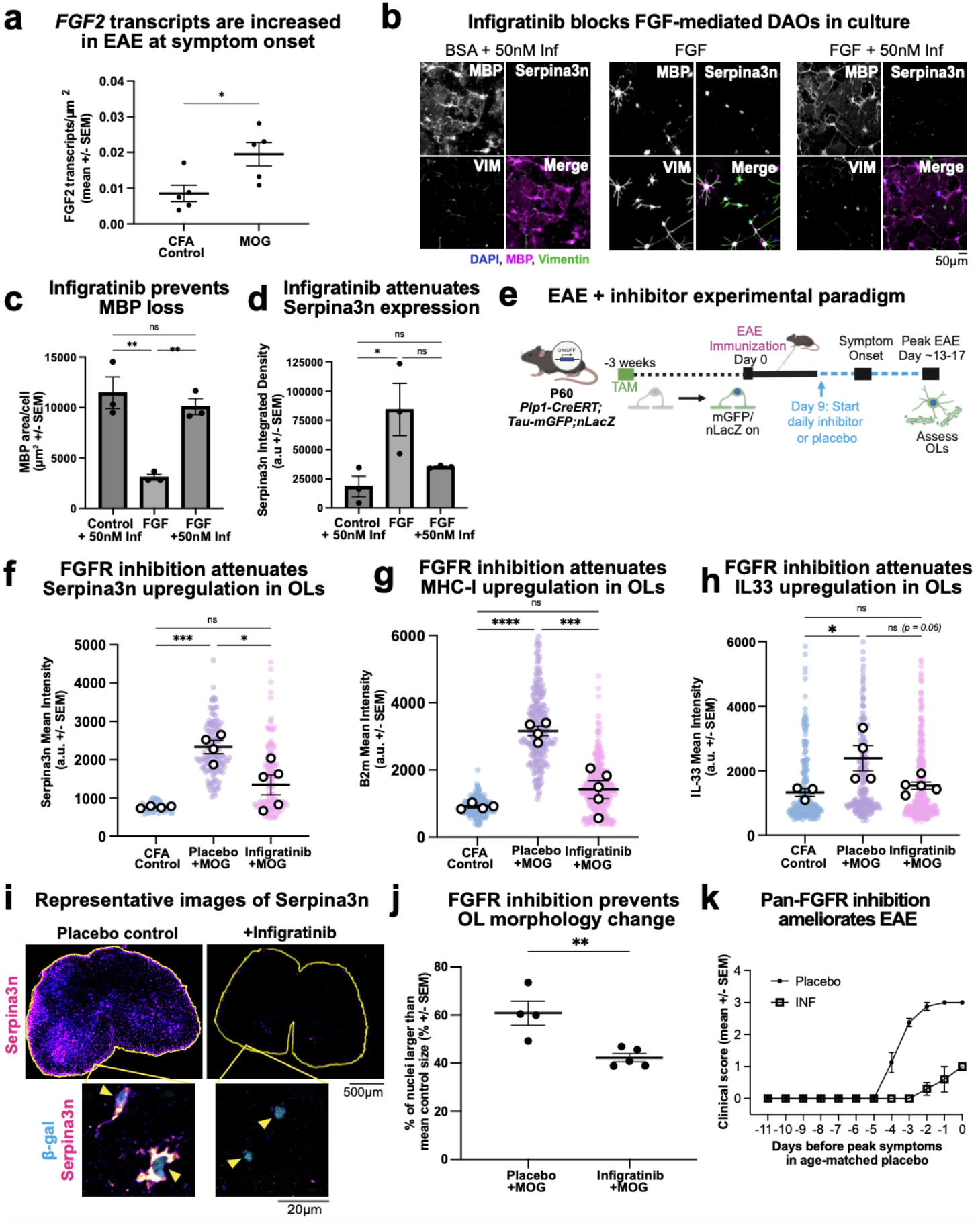
Inhibition of FGFRs reduces the formation of DAOs in EAE. **a**, Quantification of RNAscope for *FGF2* in CFA control and MOG EAE spinal cords at time of symptom onset. Number of *FGF2* transcripts per um^2^ ± SEM; CFA 0.009 ± 0.002; MOG 0.02 ± 0.003. **b**, Representative images of immunostaining for MBP, Vimentin and Serpina3n in cultured oligodendrocytes treated with BSA control (left), FGF (middle), or FGF with 50nM Infigratinib (right) for 72 hours. **c**, Quantification of MBP+ cell area in (b) showing infigratinib prevents FGF-induced loss of MBP sheets. **d**, Quantification of Serpina3n intensity in (b) showing infigratinib prevents FGF-induced expression of Serpina3n protein. **e**, Cartoon schematic of EAE with infigratinib experimental paradigm in reporter mice. Mice were immunized with MOG 3 weeks following tamoxifen, then received daily oral infigratinib or placebo starting at day 9 (before symptom onset). Tissue was harvested when age-matched placebo mice reached peak EAE score. **f**, Quantification of Serpina3n in β-gal+ cells shows reduced expression of Serpina3n in mice receiving infigratinib. Serpina3n mean intensity ± SEM: CFA: 763.7 ± 15.47 a.u.; MOG + Placebo control: 2330 ± 170.4 a.u.; MOG + Infigratinib: 1344 ± 257.4 a.u. **g**, Quantification of B2m in β-gal+ cells shows reduced expression in mice receiving infigratinib. B2m mean intensity ± SEM: CFA: 916.3 ± 44.48 a.u.; MOG + Placebo control: 3158 ± 138.6 a.u.; MOG + Infigratinib: 1417 ± 262.7 a.u. **h**, Quantification of IL33 in β-gal+ cells in EAE spinal cords shows suppression of IL33 increase in mice receiving infigratinib. IL33 mean intensity ± SEM: CFA: 1329 ± 115.6 a.u.; MOG Placebo control: 2391 ± 386.5 a.u.; MOG Infigratinib: 1535 ± 111.7 a.u. **i**, Representative images of Serpina3n expression by immunostaining in spinal cords of mice immunized with MOG and treated with either placebo (left) or infigratinib (right). Insets with yellow arrowheads highlight Serpina3n/β-gal positive DAOs. **j**, Oligodendrocyte nuclear size is restored in infigratinib treated mice. Percentage of nuclei above average control nuclei size +/SEM. Placebo: 60.87 ± 4.99%, Infigratinib: 42.27 ± 1.71%, *n* = 4 biological replicates for placebo, 5 replicates from infigratinib mice for all quantification in figure. **k**, EAE disease course represented by clinical score showing mice treated with infigratinib have fewer symptoms. Mean clinical score ± SEM, *n* = 4 biological replicates for placebo, 5 for infigratinib. Statistical measurements (*p* value) were determined by an unpaired, two-tailed t-test or 1 way ANOVA with Tukey’s multiple comparison test, *p* values: \**p* < 0.05; \*\**p* < 0.01; \*\*\**p* < 0.001; \*\*\*\**p* < 0.0001; *ns* = not significant.

**Extended Data Fig. 15.**
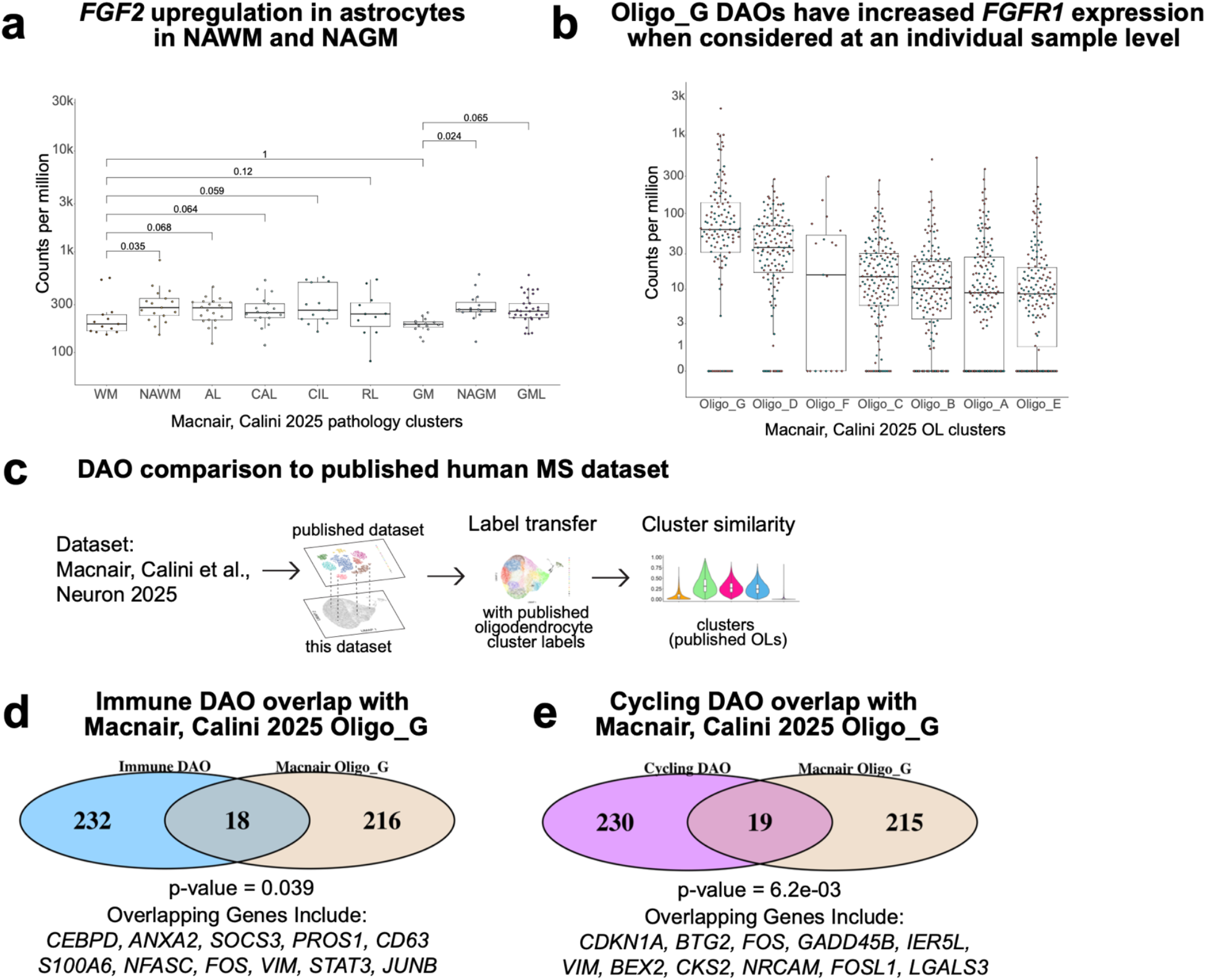
Additional human MS transcriptomic data. **a**, Box and whisker plot of FGF2 expression in astrocytes across different MS tissue lesions. FGF2 expression significantly increased (*p*<0.5) in normal-appearing white matter (NAWM) compared to control white matter (WM) and in normal-appearing grey matter (NAGM) compared to control grey matter (GM). **b**, Box and whisker plot of FGFR1 expression in distinct oligodendrocyte clusters, showing highest expression in Oligo_G DAO population. Dot color represents male vs. female patients. **c**, Workflow for single-cell dataset comparison of scRNA-seq data from FGF1/2-overexpression (this dataset) to human MS dataset from Macnair, Calini et al., 2025. **d**, Overlap between top 250 upregulated genes in Immune DAOs compared to top 250 upregulated genes in Oligo_G in human MS dataset. Fisher’s exact test was used to calculate the overlapping *p* value for each comparison. Relevant overlapping genes listed below diagram. **e**, Overlap between top 250 upregulated genes in Cycling DAOs compared to top 250 upregulated genes in Oligo_G in human MS dataset. Fisher’s exact test was used to calculate the overlapping *p* value for each comparison. Relevant overlapping genes listed below diagram.

## References

1. Stadelmann, C., Timmler, S., Barrantes-Freer, A. & Simons, M. Myelin in the Central Nervous System: Structure, Function, and Pathology. Physiol Rev 99, 1381–1431 (2019).

2. Duncan, I. D. & Radcliff, A. B. Inherited and acquired disorders of myelin: The underlying myelin pathology. Exp Neurol 283, 452–475 (2016).

3. Baecher-Allan, C., Kaskow, B. J. & Weiner, H. L. Multiple Sclerosis: Mechanisms and Immunotherapy. Neuron 97, 742–768 (2018).

4. Lanz, T. V. et al. Clonally expanded B cells in multiple sclerosis bind EBV EBNA1 and GlialCAM. Nature 603, 321–327 (2022).

5. Bjornevik, K. et al. Longitudinal analysis reveals high prevalence of Epstein-Barr virus associated with multiple sclerosis. Science 375, 296–301 (2022).

6. Hedström, A. K. Risk factors for multiple sclerosis in the context of Epstein-Barr virus infection. Front Immunol 14, 1212676 (2023).

7. Steinman, L. & Zamvil, S. S. Discoveries beyond molecular mimicry describe how EBV drives multiple sclerosis. Cell 189, 345–347 (2026).

8. Jäkel, S. et al. Altered human oligodendrocyte heterogeneity in multiple sclerosis. Nature 566, 543–547 (2019).

9. Absinta, M. et al. A lymphocyte–microglia–astrocyte axis in chronic active multiple sclerosis. Nature 597, 709–714 (2021).

10. Schirmer, L. et al. Neuronal vulnerability and multilineage diversity in multiple sclerosis. Nature 573, 75–82 (2019).

11. Grubman, A. et al. A single-cell atlas of entorhinal cortex from individuals with Alzheimer’s disease reveals cell-type-specific gene expression regulation. Nat Neurosci 22, 2087–2097 (2019).

12. Mathys, H. et al. Single-cell transcriptomic analysis of Alzheimer’s disease. Nature 570, 332–337 (2019).

13. Falcão, A. M. et al. Disease-specific oligodendrocyte lineage cells arise in multiple sclerosis. Nat. Med. 24, 1837–1844 (2018).

14. Floriddia, E. M. et al. Distinct oligodendrocyte populations have spatial preference and different responses to spinal cord injury. Nat Commun 11, 5860 (2020).

15. Shen, K. et al. Multiple sclerosis risk gene Mertk is required for microglial activation and subsequent remyelination. Cell Reports 34, 108835 (2021).

16. Zhou, Y. et al. Human and mouse single-nucleus transcriptomics reveal TREM2-dependent and TREM2-independent cellular responses in Alzheimer’s disease. Nat Med 26, 131–142 (2020).

17. Lee, S.-H. et al. TREM2-independent oligodendrocyte, astrocyte, and T cell responses to tau and amyloid pathology in mouse models of Alzheimer disease. Cell Reports 37, (2021).

18. Pandey, S. et al. Disease-associated oligodendrocyte responses across neurodegenerative diseases. Cell Reports 40, 111189 (2022).

19. Kenigsbuch, M. et al. A shared disease-associated oligodendrocyte signature among multiple CNS pathologies. Nat Neurosci 25, 876–886 (2022).

20. Hou, J. et al. Transcriptomic atlas and interaction networks of brain cells in mouse CNS demyelination and remyelination. Cell Rep 42, 112293 (2023).

21. Macnair, W. et al. snRNA-seq stratifies multiple sclerosis patients into distinct white matter glial responses. Neuron 113, 396–410.e9 (2025).

22. Castelo-Branco, G., Kukanja, P., Guerreiro-Cacais, A. O. & Rubio Rodríguez-Kirby, L. A. Disease-associated oligodendroglia: a putative nexus in neurodegeneration. Trends Immunol 45, 750–759 (2024).

23. Grinspan, J. B., Stern, J. L., Franceschini, B. & Pleasure, D. Trophic effects of basic fibroblast growth factor (bFGF) on differentiated oligodendroglia: a mechanism for regeneration of the oligodendroglial lineage. J Neurosci Res 36, 672–680 (1993).

24. Fressinaud, C., Vallat, J. M. & Labourdette, G. Basic fibroblast growth factor down-regulates myelin basic protein gene expression and alters myelin compaction of mature oligodendrocytes in vitro. J Neurosci Res 40, 285–293 (1995).

25. Grinspan, J. B., Reeves, M. F., Coulaloglou, M. J., Nathanson, D. & Pleasure, D. Re-entry into the cell cycle is required for bFGF-induced oligodendroglial dedifferentiation and survival. J Neurosci Res 46, 456–464 (1996).

26. Bansal, R. & Pfeiffer, S. E. FGF-2 converts mature oligodendrocytes to a novel phenotype. Journal of Neuroscience Research 50, 215–228 (1997).

27. Fortin, D., Rom, E., Sun, H., Yayon, A. & Bansal, R. Distinct Fibroblast Growth Factor (FGF)/FGF Receptor Signaling Pairs Initiate Diverse Cellular Responses in the Oligodendrocyte Lineage. J Neurosci 25, 7470–7479 (2005).

28. Cohen, R. I. & Chandross, K. J. Fibroblast growth factor-9 modulates the expression of myelin related proteins and multiple fibroblast growth factor receptors in developing oligodendrocytes. J Neurosci Res 61, 273–287 (2000).

29. Goddard, D. R., Berry, M., Kirvell, S. L. & Butt, A. M. Fibroblast growth factor-2 inhibits myelin production by oligodendrocytes in vivo. Mol Cell Neurosci 18, 557–569 (2001).

30. Butt, A. M. & Dinsdale, J. Fibroblast growth factor 2 mediated disruption of myelin-forming oligodendrocytes in vivo is associated with increased tau immunoreactivity. Neurosci Lett 375, 28–32 (2005).

31. Azim, K., Raineteau, O. & Butt, A. M. Intraventricular injection of FGF-2 promotes generation of oligodendrocyte-lineage cells in the postnatal and adult forebrain. Glia 60, 1977–1990 (2012).

32. Rajendran, R., Giraldo-Velásquez, M., Stadelmann, C. & Berghoff, M. Oligodendroglial fibroblast growth factor receptor 1 gene targeting protects mice from experimental autoimmune encephalomyelitis through ERK/AKT phosphorylation. Brain Pathol. 28, 212–224 (2018).

33. Rajendran, R. et al. Oligodendrocyte-Specific Deletion of FGFR1 Reduces Cerebellar Inflammation and Neurodegeneration in MOG35-55-Induced EAE. Int J Mol Sci 22, 9495 (2021).

34. Kamali, S. et al. Oligodendrocyte-specific deletion of FGFR2 ameliorates MOG35-55-induced EAE through ERK and Akt signalling. Brain Pathol 31, 297–311 (2021).

35. Rajendran, R. et al. The small molecule fibroblast growth factor receptor inhibitor infigratinib exerts anti-inflammatory effects and remyelination in a model of multiple sclerosis. British Journal of Pharmacology 180, 2989–3007 (2023).

36. Kukanja, P. et al. Cellular architecture of evolving neuroinflammatory lesions and multiple sclerosis pathology. Cell 187, 1990–2009.e19 (2024).

37. Cullen, C. L. et al. Periaxonal and nodal plasticities modulate action potential conduction in the adult mouse brain. Cell Reports 34, 108641 (2021).

38. Kaya, T. et al. CD8+ T cells induce interferon-responsive oligodendrocytes and microglia in white matter aging. Nat Neurosci 25, 1446–1457 (2022).

39. Wolswijk, G. Oligodendrocyte survival, loss and birth in lesions of chronic-stage multiple sclerosis. Brain 123 (Pt 1), 105–115 (2000).

40. Barnett, M. H. & Prineas, J. W. Relapsing and remitting multiple sclerosis: Pathology of the newly forming lesion. Annals of Neurology 55, 458–468 (2004).

41. Dugas, J. C. & Emery, B. Purification of Oligodendrocyte Precursor Cells from Rat Cortices by Immunopanning. Cold Spring Harb Protoc 2013, pdb.prot070862 (2013).

42. Aggarwal, S. et al. A size barrier limits protein diffusion at the cell surface to generate lipid-rich myelin-membrane sheets. Dev Cell 21, 445–456 (2011).

43. Zuchero, J. B. et al. CNS myelin wrapping is driven by actin disassembly. Dev Cell 34, 152–167 (2015).

44. Ornitz, D. M. & Itoh, N. The Fibroblast Growth Factor signaling pathway. Wiley Interdiscip Rev Dev Biol 4, 215–266 (2015).

45. Sofroniew, M. V. Astrocyte Reactivity: Subtypes, States, and Functions in CNS Innate Immunity. Trends Immunol 41, 758–770 (2020).

46. Thümmler, K. et al. Polarizing receptor activation dissociates fibroblast growth factor 2 mediated inhibition of myelination from its neuroprotective potential. acta neuropathol commun 7, 212 (2019).

47. Tooyama, I. et al. Acidic fibroblast growth factor-like immunoreactivity in brain of Alzheimer patients. Neurosci Lett 121, 155–158 (1991).

48. Sarchielli, P. et al. Fibroblast growth factor-2 levels are elevated in the cerebrospinal fluid of multiple sclerosis patients. Neuroscience Letters 435, 223–228 (2008).

49. Clemente, D., Ortega, M. C., Arenzana, F. J. & Castro F. de. FGF-2 and Anosmin-1 Are Selectively Expressed in Different Types of Multiple Sclerosis Lesions. J. Neurosci. 31, 14899–14909 (2011).

50. Mohan, H. et al. Transcript profiling of different types of multiple sclerosis lesions yields FGF1 as a promoter of remyelination. Acta Neuropathol Commun 2, 168 (2014).

51. Gómez-Pinilla, F., Cummings, B. J. & Cotman, C. W. Induction of basic fibroblast growth factor in Alzheimer’s disease pathology. Neuroreport 1, 211–214 (1990).

52. Messersmith, D. J., Murtie, J. C., Le, T. Q., Frost, E. E. & Armstrong, R. C. Fibroblast growth factor 2 (FGF2) and FGF receptor expression in an experimental demyelinating disease with extensive remyelination. Journal of Neuroscience Research 62, 241–256 (2000).

53. Fahmy, G. H. & Moftah, M. Z. FGF-2 in Astroglial Cells During Vertebrate Spinal Cord Recovery. Front. Cell. Neurosci. 4, (2010).

54. Shigetomi, E. et al. Imaging calcium microdomains within entire astrocyte territories and endfeet with GCaMPs expressed using adeno-associated viruses. J Gen Physiol 141, 633–647 (2013).

55. Marques, S. et al. Oligodendrocyte heterogeneity in the mouse juvenile and adult central nervous system. Science 352, 1326–1329 (2016).

56. The Gene Ontology Consortium et al. The Gene Ontology knowledgebase in 2023. Genetics 224, iyad031 (2023).

57. Ashburner, M. et al. Gene Ontology: tool for the unification of biology. Nat Genet 25, 25–29 (2000).

58. Kanehisa, M. & Goto, S. KEGG: Kyoto Encyclopedia of Genes and Genomes. Nucleic Acids Res 28, 27–30 (2000).

59. Kanehisa, M., Furumichi, M., Sato, Y., Matsuura, Y. & Ishiguro-Watanabe, M. KEGG: biological systems database as a model of the real world. Nucleic Acids Res 53, D672–D677 (2025).

60. Lee, Y. et al. Oligodendroglia metabolically support axons and contribute to neurodegeneration. Nature 487, 443–448 (2012).

61. Schirmer, L. et al. Oligodendrocyte-encoded Kir4.1 function is required for axonal integrity. Elife 7, e36428 (2018).

62. Looser, Z. J. et al. Oligodendrocyte-axon metabolic coupling is mediated by extracellular K+ and maintains axonal health. Nat Neurosci 27, 433–448 (2024).

63. Mukherjee, C. et al. Oligodendrocytes Provide Antioxidant Defense Function for Neurons by Secreting Ferritin Heavy Chain. Cell Metab 32, 259–272.e10 (2020).

64. Schönrock, Kuhlmann, Adler, Bitsch, & Brück. Identification of glial cell proliferation in early multiple sclerosis lesions. Neuropathology and Applied Neurobiology 24, 320–330 (1998).

65. Solanky, M. et al. Proliferating oligodendrocytes are present in both active and chronic inactive multiple sclerosis plaques. Journal of Neuroscience Research 65, 308–317 (2001).

66. Bansal, R., Kumar, M., Murray, K., Morrison, R. S. & Pfeiffer, S. E. Regulation of FGF Receptors in the Oligodendrocyte Lineage. Molecular and Cellular Neuroscience 7, 263–275 (1996).

67. Zhang, Y. et al. An RNA-Sequencing Transcriptome and Splicing Database of Glia, Neurons, and Vascular Cells of the Cerebral Cortex. J. Neurosci. 34, 11929–11947 (2014).

68. Zhang, Y. et al. Purification and Characterization of Progenitor and Mature Human Astrocytes Reveals Transcriptional and Functional Differences with Mouse. Neuron 89, 37–53 (2016).

69. Harrisingh, M. C. et al. The Ras/Raf/ERK signalling pathway drives Schwann cell dedifferentiation. EMBO J. 23, 3061–3071 (2004).

70. Stierli, S., Imperatore, V. & Lloyd, A. C. Schwann cell plasticity-roles in tissue homeostasis, regeneration, and disease. Glia 67, 2203–2215 (2019).

71. Clements, M. P. et al. The Wound Microenvironment Reprograms Schwann Cells to Invasive Mesenchymal-like Cells to Drive Peripheral Nerve Regeneration. Neuron 96, 98–114.e7 (2017).

72. Jessen, K. R., Mirsky, R. & Lloyd, A. C. Schwann Cells: Development and Role in Nerve Repair. Cold Spring Harb Perspect Biol 7, a020487 (2015).

73. Romanelli, E. et al. Myelinosome formation represents an early stage of oligodendrocyte damage in multiple sclerosis and its animal model. Nat Commun 7, 13275 (2016).

74. Luchicchi, A. et al. Axon-Myelin Unit Blistering as Early Event in MS Normal Appearing White Matter. Ann Neurol 89, 711–725 (2021).

75. Kedia, S. et al. T cell-mediated microglial activation triggers myelin pathology in a mouse model of amyloidosis. Nat Neurosci 27, 1468–1474 (2024).

76. Lerma-Martin, C. et al. Cell type mapping reveals tissue niches and interactions in subcortical multiple sclerosis lesions. Nat Neurosci 27, 2354–2365 (2024).

77. Su, J. J. et al. Upregulation of vascular growth factors in multiple sclerosis: Correlation with MRI findings. Journal of the Neurological Sciences 243, 21–30 (2006).

78. Rajendran, R., Böttiger, G., Stadelmann, C., Karnati, S. & Berghoff, M. FGF/FGFR Pathways in Multiple Sclerosis and in Its Disease Models. Cells 10, 884 (2021).

79. Tobin, J. E., Xie, M., Le, T. Q.Song, S.-K. & Armstrong, R. C. Reduced Axonopathy and Enhanced Remyelination Following Chronic Demyelination in Fibroblast Growth Factor-2 (Fgf2) Null Mice: Differential Detection with Diffusion Tensor Imaging. J Neuropathol Exp Neurol 70, 157–165 (2011).

80. Mierzwa, A. J., Zhou, Y.-X., Hibbits, N., Vana, A. C. & Armstrong, R. C. FGF2 and FGFR1 signaling regulate functional recovery following cuprizone demyelination. Neurosci Lett 548, 280–285 (2013).

81. Rothaug, M., Becker-Pauly, C. & Rose-John, S. The role of interleukin-6 signaling in nervous tissue. Biochim Biophys Acta 1863, 1218–1227 (2016).

82. Haile, Y. et al. Granzyme B-inhibitor serpina3n induces neuroprotection in vitro and in vivo. J Neuroinflammation 12, 157 (2015).

83. Zhu, M. et al. A paradigm shift of SERPINA3N in neurobehavioral development and brain injury. 2025.09.09.675167 Preprint at 10.1101/2025.09.09.675167 (2025).

84. Klimaschewski, L. & Claus, P. Fibroblast Growth Factor Signalling in the Diseased Nervous System. Mol Neurobiol 58, 3884–3902 (2021).

85. Turner, C. A., Eren-Koçak, E., Inui, E. G., Watson, S. J. & Akil, H. Dysregulated fibroblast growth factor (FGF) signaling in neurological and psychiatric disorders. Semin Cell Dev Biol 53, 136–143 (2016).

86. Kimura, H., Tooyama, I. & Mcgeer, P. L. Acidic FGF Expression in the Surroundings of Senile Plaques. The Tohoku Journal of Experimental Medicine 174, 279–293 (1994).

87. Mashayekhi, F., Hadavi, M., Vaziri, H. R. & Naji, M. Increased acidic fibroblast growth factor concentrations in the serum and cerebrospinal fluid of patients with Alzheimer’s disease. Journal of Clinical Neuroscience 17, 357–359 (2010).

88. Zhang, S. et al. Oligodendrocyte-astrocyte crosstalk in Parkinson’s disease mediates neuronal ferroptosis via the FGF signaling pathway. npj Parkinsons Dis. 11, 136 (2025).

89. Du, S. et al. Mutations in the human CSF1R gene impact microglia’s maintenance of brain white matter integrity. Nat Immunol 26, 1198–1211 (2025).

90. Guo, Y. et al. High-titer AAV disrupts cerebrovascular integrity and induces lymphocyte infiltration in adult mouse brain. Mol Ther Methods Clin Dev 31, 101102 (2023).

91. Paliard, X. et al. Simultaneous production of IL-2, IL-4, and IFN-gamma by activated human CD4+ and CD8+ T cell clones. J Immunol 141, 849–855 (1988).

92. Furusho, M., Dupree, J. L.Nave, K.-A. & Bansal, R. Fibroblast Growth Factor Receptor Signaling in Oligodendrocytes Regulates Myelin Sheath Thickness. J Neurosci 32, 6631–6641 (2012).

93. Furusho, M., Ishii, A. & Bansal, R. Signaling by FGF Receptor 2, Not FGF Receptor 1, Regulates Myelin Thickness through Activation of ERK1/2-MAPK, Which Promotes mTORC1 Activity in an Akt-Independent Manner. J Neurosci 37, 2931–2946 (2017).

94. Bercury, K. K. & Macklin, W. B. Dynamics and mechanisms of CNS myelination. Dev Cell 32, 447–458 (2015).

95. Ishii, A., Furusho, M., Dupree, J. L. & Bansal, R. Strength of ERK1/2 MAPK Activation Determines Its Effect on Myelin and Axonal Integrity in the Adult CNS. J Neurosci 36, 6471–6487 (2016).

96. Park, H. et al. Single-cell RNA-sequencing identifies disease-associated oligodendrocytes in male APP NL-G-F and 5XFAD mice. Nat Commun 14, 802 (2023).

97. Bell, K. A., O’Riordan, K. J., Sweatt, J. D. & Dineley, K. T. MAPK recruitment by beta-amyloid in organotypic hippocampal slice cultures depends on physical state and exposure time. J Neurochem 91, 349–361 (2004).

98. Huang, Y.-W. A., Zhou, B., Wernig, M. & Südhof, T. C. ApoE2, ApoE3, and ApoE4 Differentially Stimulate APP Transcription and Aβ Secretion. Cell 168, 427–441.e21 (2017).

99. Reuss, B. & von Bohlen und Halbach, O. Fibroblast growth factors and their receptors in the central nervous system. Cell Tissue Res 313, 139–157 (2003).

100. Besnard, F., Perraud, F., Sensenbrenner, M. & Labourdette, G. Effects of acidic and basic fibroblast growth factors on proliferation and maturation of cultured rat oligodendrocytes. Int J Dev Neurosci 7, 401–409 (1989).

101. Bansal, R., Magge, S. & Winkler, S. Specific inhibitor of FGF receptor signaling: FGF-2-mediated effects on proliferation, differentiation, and MAPK activation are inhibited by PD173074 in oligodendrocyte-lineage cells. Journal of Neuroscience Research 74, 486–493 (2003).

102. Zhai, W. et al. The fibroblast growth factor system in cognitive disorders and dementia. Front. Neurosci. 17, (2023).

103. Korbie, D. J. & Mattick, J. S. Touchdown PCR for increased specificity and sensitivity in PCR amplification. Nat Protoc 3, 1452–1456 (2008).

104. Lam, M. et al. CNS myelination requires VAMP2/3-mediated membrane expansion in oligodendrocytes. Nat Commun 13, 5583 (2022).

105. Forough, R. et al. Differential transforming abilities of non-secreted and secreted forms of human fibroblast growth factor-1. J Biol Chem 268, 2960–2968 (1993).

106. Foo, L. C. et al. Development of a Novel Method for the Purification and Culture of Rodent Astrocytes. Neuron 71, 799–811 (2011).

107. Bin, J. M., Harris, S. N. & Kennedy, T. E. The oligodendrocyte-specific antibody ‘CC1’ binds Quaking 7. J Neurochem 139, 181–186 (2016).

108. Madhavarao, C. N. et al. Immunohistochemical localization of aspartoacylase in the rat central nervous system. J Comp Neurol 472, 318–329 (2004).

109. Bankhead, P. et al. QuPath: Open source software for digital pathology image analysis. Sci Rep 7, 16878 (2017).

110. Hafemeister, C. & Satija, R. Normalization and variance stabilization of single-cell RNA-seq data using regularized negative binomial regression. Genome Biol 20, 296 (2019).

111. Stuart, T. et al. Comprehensive Integration of Single-Cell Data. Cell 177, 1888–1902.e21 (2019).

112. Friedman, P. L. & Ellisman, M. H. Enhanced visualization of peripheral nerve and sensory receptors in the scanning electron microscope using cryofracture and osmium-thiocarbohydrazide-osmium impregnation. J Neurocytol 10, 111–131 (1981).

113. Willingham, M. C. & Rutherford, A. V. The use of osmiumthiocarbohydrazide-osmium (OTO) and ferrocyanidereduced osmium methods to enhance membrane contrast and preservation in cultured cells. J Histochem Cytochem 32, 455–460 (1984).

114. Ewald, A. J. et al. Mammary collective cell migration involves transient loss of epithelial features and individual cell migration within the epithelium. J Cell Sci 125, 2638–2654 (2012).

115. McDonald, K. L. & Webb, R. I. Freeze substitution in 3 hours or less. J Microsc 243, 227–233 (2011).

116. Wang, J. et al. Reactive oxygen FIB spin milling enables correlative workflow for 3D super-resolution light microscopy and serial FIB/SEM of cultured cells. Sci Rep 11, 13162 (2021).

117. Harterink, M. et al. DeActs: genetically encoded tools for perturbing the actin cytoskeleton in single cells. Nat Methods 14, 479–482 (2017).

